# From *in silico* prediction to experimental validation: Identification of drugs and novel synergistic combinations that inhibit growth of inflammatory breast cancer cells

**DOI:** 10.64898/2025.12.10.693562

**Authors:** Esraa A. Salim, Xiaojia Ji, Michael Tarpley, Maria S. Dixon, Weifan Zheng, John E. Scott, Kevin P. Williams

## Abstract

Drug repurposing offers a promising approach for identifying novel treatments, especially for rare cancers like inflammatory breast cancer (IBC), an aggressive type with limited therapeutic options. Here, we present a comprehensive validation and verification study of compounds identified through two computational approaches: Literature Wide Association Studies (LWAS) and Gene Reversal Rate (GRR), using orthogonal cell viability assays in 2D models across IBC and non-IBC cell lines. In the SUM149 IBC cell line, repurposed compounds predicted from LWAS achieved a 70% success rate, with several showing nanomolar potency, while those predicted from GRR showed a 38% success rate. Through systematic combination screening in both 2D and 3D-spheroid models, we identified novel synergistic compound pairs targeting crosstalk between IGF1-R, EGFR and PI3K/Akt/mTOR pathways, with high synergy scores across multiple reference models. Using these combinations, western blotting analysis revealed significant suppression in the phosphorylation of key signaling proteins and downstream effectors, while wound healing assays demonstrated reduced cell migration for some combinations, suggesting effective pathway inhibition. To further validate these findings at the transcriptional level, RNA-Seq analysis in SUM149 cells confirmed that the GRR drug combinations significantly reversed the IBC gene expression signature (IBC-GES). These findings not only validated our computational predictions but also identified promising combination strategies that could potentially overcome drug resistance in IBC. Our integrated computational-experimental approach establishes a framework for systematic drug repurposing and highlights novel therapeutic combinations warranting further investigation.

## Introduction

Inflammatory breast cancer (IBC) is an aggressive and uncommon type of breast cancer [1–3]. In Western nations, IBC accounts for up to 5% of breast cancer cases [4, 5], with higher rates among African American [6] and North African populations, reaching 7-11.1% [7]. Although considered rare, IBC is responsible for 8-10% of all breast cancer deaths [8]. At diagnosis, up to 85% of IBC patients have lymph node involvement, and up to 30% show distant metastasis [9, 10]. The aggressive nature of IBC and the presence of unique characteristics in its presentation [11, 12] and progression contribute significantly to the higher rates of treatment resistance, relapse, and mortality [13]. Despite the widespread adoption of trimodality treatment for IBC, combining anthracycline- and taxane-based neoadjuvant chemotherapy (NACT), surgery, and radiation therapy [14], the 5-year and 10-year overall survival (OS) rates remain low at 55.4% and 37.3%, respectively [15, 16]. Currently, there is no Food and Drug Administration (FDA)-approved targeted therapy specific to IBC and the current treatment strategies are primarily based on findings from studies of high-risk non-IBC patients [17]. This critical gap in treatment options highlights the essential need for innovative therapeutic strategies to improve patient outcomes in rare diseases. Several drug discovery approaches can be undertaken to find novel drugs for IBC, including drug repurposing which can be a promising approach to address this critical need for new treatments.

Drug repurposing, the application of existing, market-approved drugs to new disease indications, offers significant advantages over *de novo* drug discovery [18]. This approach aims to enhance the drug discovery process, by reducing both time and cost [19, 20]. By utilizing known drugs with established safety profiles, drug repurposing can bypass intermediate steps and accelerate the path from hypothesis to clinical application [21–23]. The primary objective of drug repurposing is to identify novel associations between existing drugs and diseases through two main approaches: computational prediction and experimental validation [24]. Modern computational drug repurposing technologies seek to systemically analyze diverse data types, including genomic, proteomic, clinical health records, and scientific literature to predict potential drug-disease connections [20, 25, 26]. Our previous studies utilized two distinct computational approaches [27, 28] to identify repurposing candidates for IBC treatment. Our first approach utilized Literature Wide Association Study (LWAS) [27], which applies extensive text mining [29] as a powerful tool to identify new repurposed candidates. Our second approach used Gene Reversal Rate (GRR) [28], which employs gene expression reversal methodology [30, 31] to identify compounds that are capable of reversing cancer-associated gene expression patterns. Complementing these computational methods, experimental validation approaches involve direct testing of the predicted compounds in various disease models to enhance the likelihood of successful drug repurposing.

This current study aimed to assess the efficacy of the drug and compound candidates we had predicted for IBC from our LWAS and GRR approaches using *in vitro* screening techniques in 2D and 3D IBC and non-IBC cell-based models. We used different cell viability and proliferation assays to evaluate drug potency and efficacy, both as single agents and in combinations. Our initial findings identified anti-proliferative effects for a subset of drugs in both IBC and non-IBC cell lines, with 70% (19 of 24) of the LWAS-predicted compounds and 38% (6 of 16) of the GRR-predicted compounds showing efficacy in the SUM149 IBC cell line. To address potential drug resistance to single compounds, we used a drug combination assay matrix and synergy analysis pipeline to identify potential synergistic combinations. Three novel synergistic combinations were identified with significant effects on proliferation, spheroid integrity, protein expression, and cell migration in IBC cell models. These combinations comprising compounds predicted from our GRR study and had not been previously reported in IBC clinical trials or the literature. In addition to the phenotypic and protein-level validation, we further assessed transcriptional responses to these GRR drug combinations using RNA-Seq and observed a significant reversal of genes in the IBC gene expression signature. In this study, the synergistic GRR combinations that we identified act on insulin-like growth factor 1 receptor (IGF1-R) and epidermal growth factor receptor (EGFR) signaling, as well as downstream PI3K/Akt/mTOR signaling, suggesting a promising avenue for targeting IBC.

## Materials and methods

### Cell lines and compounds

Four breast cancer cell lines were used in this study: SUM149PT (SUM149) and SUM159PT (SUM159) (obtained from BioIVT, Westbury, NY, USA), and MDA-MB-231 and MCF-7 (obtained from American Type Culture Collection, ATCC, Manassas, VA, USA). Compounds identified through LWAS [27] (n = 24) and GRR [28] (n = 16 were commercially available out of the 19 identified) studies were purchased from Selleck Chemicals (Houston, TX, USA) and Sigma-Aldrich (St. Louis, MO, USA). All compounds were supplied as powders, reconstituted in 100% DMSO to a stock concentration of 10 mM, and stored at −20°C.

### Cell culture

SUM149 and SUM159 cells were cultured in Ham’s F-12 medium supplemented with 5% FBS, 1 µg/ml hydrocortisone, 5 µg/ml insulin, 1% of 10,000 units/mL Penicillin/Streptomycin, and 10 mM HEPES, according to BioIVT guidelines. MCF-7 cells were maintained in EMEM supplemented with 0.01 mg/ml human recombinant insulin, 10% FBS, and 1% Penicillin/Streptomycin, following ATCC protocols. MDA-MB-231 cells were cultured in Leibovitz’s L-15 Medium supplemented with 10% FBS and 1% Penicillin/Streptomycin, as recommended by ATCC. SUM149, SUM159, and MCF-7 cells were incubated at 37°C in a humidified atmosphere with 5% CO_2_. MDA-MB-231 cells were maintained at 37°C with free gas exchange and 0% CO_2_.

### High content imaging cell proliferation assay

Cell proliferation was quantified using an automated Hoechst 33342 DNA staining method, as previously described [32, 33]. Breast cancer cell lines (SUM149, SUM159, MDA-MB-231, and MCF-7) were seeded in 384-well microplates (Corning, 3764) at optimized densities of 800, 600, 1600, and 2500 cells/well, respectively. Cell suspensions were dispensed at 50 µL per well using a MultiFlo Dispenser (Agilent BioTek). Each plate included negative controls (0.1% DMSO) and positive controls (Staurosporine at 10 µM and 1 µM). After 24 hours, cells were treated with test compounds dispensed by a D300 Digital Dispenser (HP Inc., Palo Alto, CA, USA). The compounds were solubilized in DMSO to prepare 10 mM stock solutions and dispensed in a 20-point dose-response range using 1:2 serial dilutions, starting with 10 µM concentration. After 72 h of treatment, cells were stained with Hoechst 33342 using a Biomek NX Automated Workstation (Beckman Coulter, Brea, CA, USA). The dye was added at a final concentration of 10 µg/mL, and plates were incubated for 45 min at 37°C. The culture medium and dye were then aspirated, and cells were fixed with 20 µL of 4% formalin for 15 min at room temperature. After removing the fixative, 50 µL of PBS were added to each well. Cell nuclei were imaged and quantified using a CellInsight CX7 High-Content Screening (HCS) Platform (Thermo Fisher Scientific). Five fields per well were captured using a 10× objective. Data were analyzed using GraphPad Prism 9.0 (GraphPad Software, San Diego, CA, USA). All experiments were performed in triplicate technical replicates with three independent experiments, and results are presented as mean ± SD.

### Automated cell viability assay

Cell viability was evaluated in four breast cancer cell lines (SUM149, SUM159, MDA-MB-231, and MCF-7) using an automated high-throughput screening approach [34]. The cells were seeded at optimized densities (SUM149: 800 cells/well; SUM159: 600 cells/well; MDA-MB-231: 4000 cells/well; MCF-7: 2500 cells/well) in 384-well flat-bottom polystyrene microplates (Corning, Cat. #3701). Cell suspensions were dispensed using a MultiFlo Dispenser (Agilent BioTek, Santa Clara, CA, USA) in 50 µL of complete growth medium per well. Following 24 h, cells were treated with test compounds in a dose-response format as described above. Each plate contains maximum solvent control wells (0.1 % DMSO) and minimum control wells (no cells, culture media only). Cell viability was quantified 72 h post-treatment using the MTT (3-(4,5-dimethylthiazol-2-yl)-2,5-diphenyltetrazolium bromide) colorimetric assay. MTT solution was delivered to each well at a final concentration of 0.5 mg/mL using a Biomek NX Automated Workstation (Beckman Coulter, Brea, CA, USA). After a 4 h incubation at 37°C to allow formazan crystal formation, the culture medium was aspirated, and the crystals were solubilized in 40 µL DMSO during a 1 h dark incubation at room temperature. Absorbance was measured at 550 nm using a SpectraMax Microplate Reader (Molecular Devices, San Jose, CA, USA). Cell viability percentages were calculated by normalizing sample absorbance values to negative control measurements. Dose-response curves were generated, and the half-maximal inhibitory concentration (IC_50_) was calculated using GraphPad Prism 9.0 (GraphPad Software, San Diego, CA, USA). All experiments were conducted in biological triplicate, with data presented as mean ± standard deviation (SD).

### Drug combination screening

The synergistic effects of drug combinations were evaluated in 2D SUM149 cells using the Hoechst 33342 DNA staining assay. This assay was performed over five days as described in above. The combination effects were assessed in two sequential phases: primary screening followed by secondary validation screening.

#### Primary combination screening: single-dose drug combinations

Every two compounds were combined in the same well at their respective EC_25_ values to get a singular dose. The concentration that achieves 25% inhibition (EC_25_) values for each drug was calculated using the following equation, in which F is the desired response level, and H is the Hill slope:

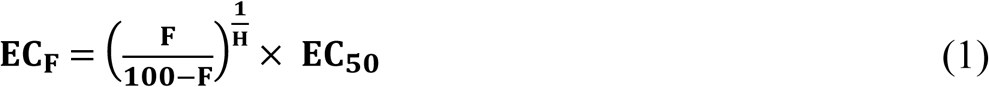

In total, 78 combinations from the LWAS study and 15 combinations from the GRR study were evaluated. Synergy analysis was performed using the Bliss excess Independence method [35] using the following equation:

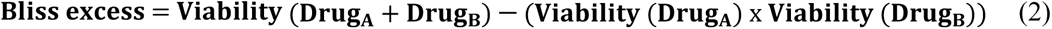

The Bliss excess score, with negative values indicating synergy, was calculated across quadruplicate technical replicates to assess statistical significance.

#### Secondary drug combination screening: matrix-designed drug treatments

An 8×8 dose-response matrix [36, 37] was designed using ½ log serial dilutions for the top five combinations from each list, based on the Bliss synergy scores from the primary screening. This was implemented using the Synergy Wizard feature in the D300 Control Software (HP Inc.). The highest concentration in each matrix was selected based on each drug’s IC_50_ value. Staurosporine (10 µM) was used as a positive control and DMSO (0.2%) as a negative control. DMSO tolerance in SUM149 cells was confirmed up to 0.8%. The experiment was conducted with three technical replicates and three independent experiments. Percent inhibition and synergy scores were calculated and analyzed using SynergyFinder3 [38] SynergyFinder+ [39] web-based tools.

### Quantification of cell viability in SUM149 spheroids

Cell viability was assessed using the CellTiter-Glo Luminescent Cell Viability Assay (Promega, Madison, WI, USA) according to the manufacturer’s instructions. Briefly, SUM149 cells were plated in 384 microplates (Corning, Cat. #3830) at 5000 cells per well and allowed to form a spheroid for 72 h at 37°C in a humidified 5% CO_2_ incubator. Spheroids were then treated with selected combinations (based on a 2D combination experiment) in an 8×8 matrix as described previously, using Staurosporine and DMSO as controls. After 72 h, plates were incubated at room temperature for 30 min. An equal volume of CellTiter-Glo reagent was added to each well, and plates were then shaken on a shaker for 5 minutes to promote cell lysis, followed by 30 min incubation at room temperature. Luminescence was measured using a PHERAstar FSX detection system (BMG LABTECH, Cary, NC, USA). Cell viability was calculated as a percentage relative to vehicle-treated control cells. Each experiment was performed in triplicate and repeated in three independent experiments. Data were analyzed using SynergyFinder+ and SynergyFinder 3 web-based tools.

### Cell death evaluation and spheroid imaging

SYTOX Red Dead Cell Stain (Invitrogen, Cat. No. S34859) was used as a qualitative tool to assess and image the spheroid shape and integrity. Spheroids were formed and treated in 384-well microplates as described above in the CellTiter-Glo assay (2.6). SYTOX Red was added to a final concentration of 40 nM immediately after treatment. Imaging was performed using an IncuCyte S3 Live-Cell Analysis System (Sartorius, Ann Arbor, MI, USA) equipped with a 10× objective. The IncuCyte system was maintained at 37°C and 5% CO_2_ throughout the experiment. Images were acquired every 6 h for 3 days, capturing both brightfield and red fluorescence (excitation 585 nm, emission 635 nm) channels. Spheroid size and fluorescence intensity were quantified using the IncuCyte S3 software. A minimum of four spheroids were analyzed per condition.

### Wound healing and cell migration

SUM149 cells were seeded at 60,000 cells per well in an interlock 96-well plate (Essen Bioscience, Ann Arbor, Michigan) and allowed to attach overnight at 37°C in a humidified atmosphere with 5% CO_2_. A uniform vertical scratch wound was created across all wells using the automated WoundMaker™ tool (Essen BioScience). Cells were gently washed with media to remove detached cells, and fresh culture medium was added before the combination treatment using a D300 Digital Dispenser (HP). Cell migration into the wound area was monitored in real-time using the IncuCyte® S3 Live-Cell Analysis System (Sartorius) as previously described [33]. Images were automatically captured at 0, 6-, 12-, 18-, and 24-h post-wounding using a 10× objective lens, enabling quantitative assessment of wound closure kinetics.

### Western blotting

SUM149 cell lysates (after 24 h drug treatment) were collected using M-PER buffer supplemented with protease/phosphatase inhibitors (Thermo Fisher Scientific, Cat. #A32959). Lysates were centrifuged (14,000 xg, 15 min, 4°C) and protein concentrations were determined by BCA assay (Pierce, Cat. #23225). Equal protein amounts (15-30 μg/lane) were loaded, and samples were electrophoresed on 4-12% Bis-Tris gels (Invitrogen, Cat. #NP0336BOX). For mTOR and p-mTOR, samples were run on 8% Tris-Glycine gels (Invitrogen, Cat. #XP00085BOX). Proteins were transferred to 0.45 μm PVDF membranes (Invitrogen, Cat. #LC2005) using XCell II™ Blot Module (Overnight, 4°C). Membranes were blocked with 5% BSA in TBS (pH 7.4) for 1 h at room temperature. Primary antibodies were diluted in 5% BSA in TBS-T (0.1% Tween-20) and incubated with membranes overnight at 4°C. All primary antibodies were used at 1:1000 dilution except anti-β-Actin and p-Akt(Ser473) which were used at 1:2000. Antibodies included: anti-β-Actin (45 kDa, LI-COR, #926-42212), AKT (60 kDa, CST, #9272), p-Akt(Ser473) (60 kDa, CST, #4060), IGF-1R (95 kDa, CST, #3027), p-IGF-1R(Tyr1135/1136) (95 kDa, CST, #3024), EGFR (175 kDa, CST, #4267), p-EGFR(Tyr1068) (175 kDa, CST, #3777), mTOR (289 kDa, CST, #2983), and p-mTOR(Ser2448) (289 kDa, CST, #2971). After TBS-T washes (3×10 min), membranes were incubated with either IRDye secondary antibodies (800CW/680RD, LI-COR, 1:20,000 in 5% BSA/TBS-T [0.2% Tween-20]/0.02% SDS) or HRP-linked antibody (1:10,000, CST, 7074) for 1 h at room temperature. Following TBS-T washes (3×5 min), membranes were imaged using Odyssey CLx (LI-COR) or iBright Systems (Invitrogen). For IGF1-R and pIGF1-R (and β-Actin control for these samples), HRP secondary antibody was used for detection, and blots imaged by iBright. Analysis was performed using Image Studio (v5.2, LI-COR) or iBright software. Band intensities were normalized to total protein stain (for LI-COR) and β-Actin (for iBright). Experiments were performed in triplicate (n=3), with significance determined as *p* < 0.05.

### RNA extraction and RNA Seq analysis

Total RNA was extracted from SUM149 cultured cells after 24 h treatment with either combinations or DMSO vehicle control, using the RNeasy Mini Kit (Qiagen, 74104), following the manufacturer’s instructions. RNA concentration and purity were assessed using a NanoDrop 2000 spectrophotometer (Thermo Fisher Scientific). The absorbance ratios A260/A280 and A260/A230 were used to evaluate RNA quality and purity. Samples with A260/A280 ratios between 1.8 and 2.1 and A260/A230 ratios above 1.8 were considered of suitable quality for RNA-Seq analysis. All samples were prepared as independent replicates (n = 3). RNA-Seq analysis was performed by Novogen (Novogene Corporation Inc., Sacramento, CA, USA) following their standard pipeline, beginning with the preparation of RNA-Seq libraries using the KAPA mRNA HyperPrep kit, which were then sequenced on an Illumina NovaSeq 6000/X Plus platform to produce 100-bp paired-end reads. Raw reads underwent quality control with FastQC; adapters and low-quality bases were trimmed using fastq-mcf and cutadapt. Potential contaminants (rRNA, tRNA, adapters, mitochondrial sequences) were filtered with Bowtie2, and the remaining reads were aligned to the human Ensembl GRCh38 (release 87) reference using STAR in two-pass mode. Gene-level counts were assigned with HTSeq and count matrices were normalized with DESeq2 in R. Differential expression was carried out in DESeq2 using contrasts named Control_vs_Case (direction reported for the case group relative to control) with significance defined as padj < 0.05 and |Log_2_fold-change| ≥ 2.

### Data processing and statistical analysis

Statistical analyses and data visualizations were performed using GraphPad Prism 9 (GraphPad Software, Inc., San Diego, CA, USA), with statistical significance set at p < 0.05 for all analyses. IC_50_ values were determined using nonlinear logistic regression analysis with a three-parameter log(inhibitor) vs. dose-response function, constrained to a zero-bottom asymptote. SynergyFinder3 and SynergyFinder+, web-based tools, were used to calculate synergy scores based on four widely used reference models: Bliss, Loewe, Highest Single Agent (HSA), and Zero interaction potency (ZIP). All data are presented as mean ± SD, unless otherwise noted.

## Results

### Assessment of LWAS and GRR drug candidates for inhibitory effects in breast cancer cell lines

To assess the efficacy of the identified drug and compound candidates from our previous LWAS [27] and GRR [28] studies, we conducted high-throughput screening (HTS) using dose-response treatments in an automated 384-well format. Two orthogonal cell-based assays were utilized to determine the efficacy and potency of each candidate: the Hoechst 33342 assay, a fluorescent DNA staining method for assessing nuclei count (as a readout for cell number) using high-content imaging [32], and the MTT assay to assess effects on cell viability. Screening was conducted across a panel of IBC and non-IBC cell lines: SUM149, SUM159, MDA-MB-231, and MCF-7.

#### Identification of LWAS candidates with effects on cell proliferation

Our previous LWAS text mining study [27], used a natural language processing-based model trained on a large corpus of cancer-related abstracts to identify diseases that co-occur with IBC, with the concept that drugs for those diseases that cluster with IBC can be potentially repurposed for IBC. This LWAS analysis identified oncology-based compounds that we predicted could be repurposed for IBC [27] and comprised 24 compounds, across various mechanistic classes: tyrosine kinase inhibitors (TKIs), antimetabolites, topoisomerase inhibitors/anthracycline antibiotics, microtubule inhibitors, and alkylating agents (**S1 Table**). In this current study, we first assessed the activity of these 24 compounds [27] in our lab-based Hoechst assay to determine effects on cell number and viability. We found significant antiproliferative activity (IC_50_ < 10 μM) for 17 compounds in SUM149, 16 in SUM159, and 15 in both MDA-MB-231 and MCF-7 cell lines (shown in heatmap format in **Fig 1A**).

**Fig 1.**
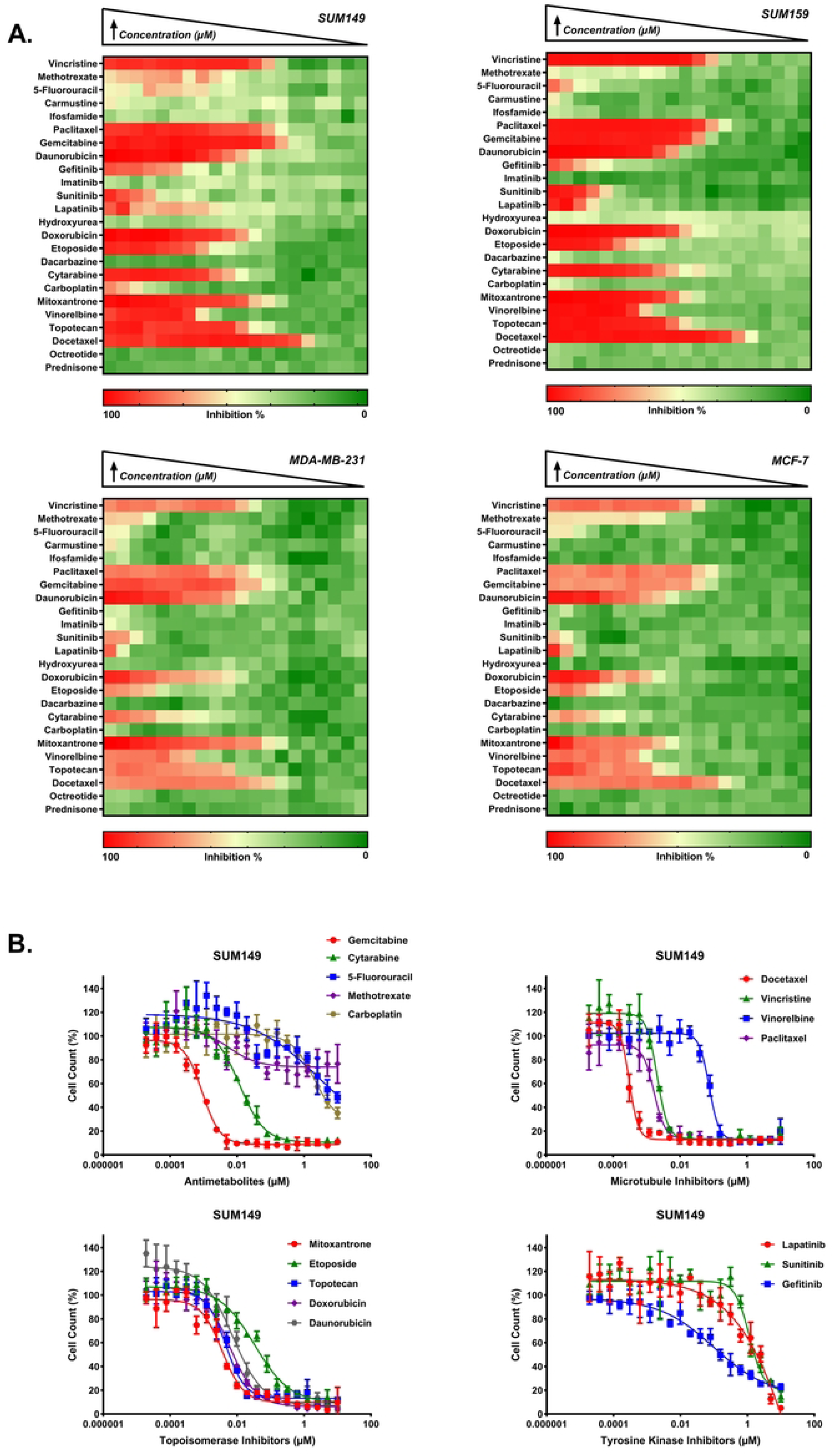
LWAS drug sensitivity profiles in IBC and non-IBC breast cancer cell line panel by Hoechst assay. (A) Heatmaps representing the percent inhibition response for a 2-fold concentration dose-response starting at 10 µM for SUM149, SUM159, MDA-MB-231, and MCF-7. Drug names are listed on the y-axis. The color scale indicates the level of inhibition, with red representing high inhibition (close to 100%) and green representing low inhibition (close to 0%). (B) Representative dose response curves (DRC) for the LWAS drugs with activity in the SUM149 cell line, showing the percent cell count relative to control against drug concentration (µM). Plots are grouped by drug mechanism: antimetabolites, microtubule inhibitors, topoisomerase inhibitors, and tyrosine kinase inhibitors. Data points and error bars represent the mean ± SD. DRCs for the LWAS drugs in SUM159, MDA-MB-231, and MCF-7 are in **S1 Fig**.

Comparative analysis across cell lines revealed a graded pattern of drug sensitivity, with SUM149 demonstrating the highest responsiveness, particularly to microtubule inhibitors and antimetabolites. Analysis of median IC_50_ values across compound classes showed decreasing sensitivity in the following order: SUM149 > SUM159 > MDA-MB-231 > MCF-7, which may correlate with these cells’ relative doubling times. Among the compound classes, microtubule inhibitors demonstrated the highest potency, with docetaxel, paclitaxel, and vincristine showing low nanomolar activity (IC_50_: 0.2-13 nM) across all cell lines. Topoisomerase inhibitors represented the second-highest set of potent compounds, with doxorubicin and daunorubicin achieving IC_50_ values below 0.2 μM. The tyrosine kinase inhibitors lapatinib and sunitinib showed moderate micromolar potency (IC_50_: 0.57-7.3 μM and 1.4-5.3 μM, respectively), while gefitinib showed selective activity against SUM149 and SUM159 (0.4-10 μM, respectively). Within the antimetabolite class, gemcitabine yielded high potency (IC_50_: 0.001-0.29 μM) and cytarabine yielded moderate potency (IC_50_: 0.023-0.65 μM). The alkylating agent carboplatin established the lowest potency, with activity only in SUM149 (IC_50_ = 6.04 μM). The remaining compounds from the LWAS screen (n = 7) showed minimal antiproliferative effects against SUM149 (IC_50_ > 10 μM). Representative dose-response curves (DRC) for SUM149 are shown in **Fig 1B**, while DRCs for SUM159, MDA-MB-231, and MCF-7 cell lines are in **S1 Fig**. Complete IC_50_ values are summarized in **Table 1**.

**Table 1.**
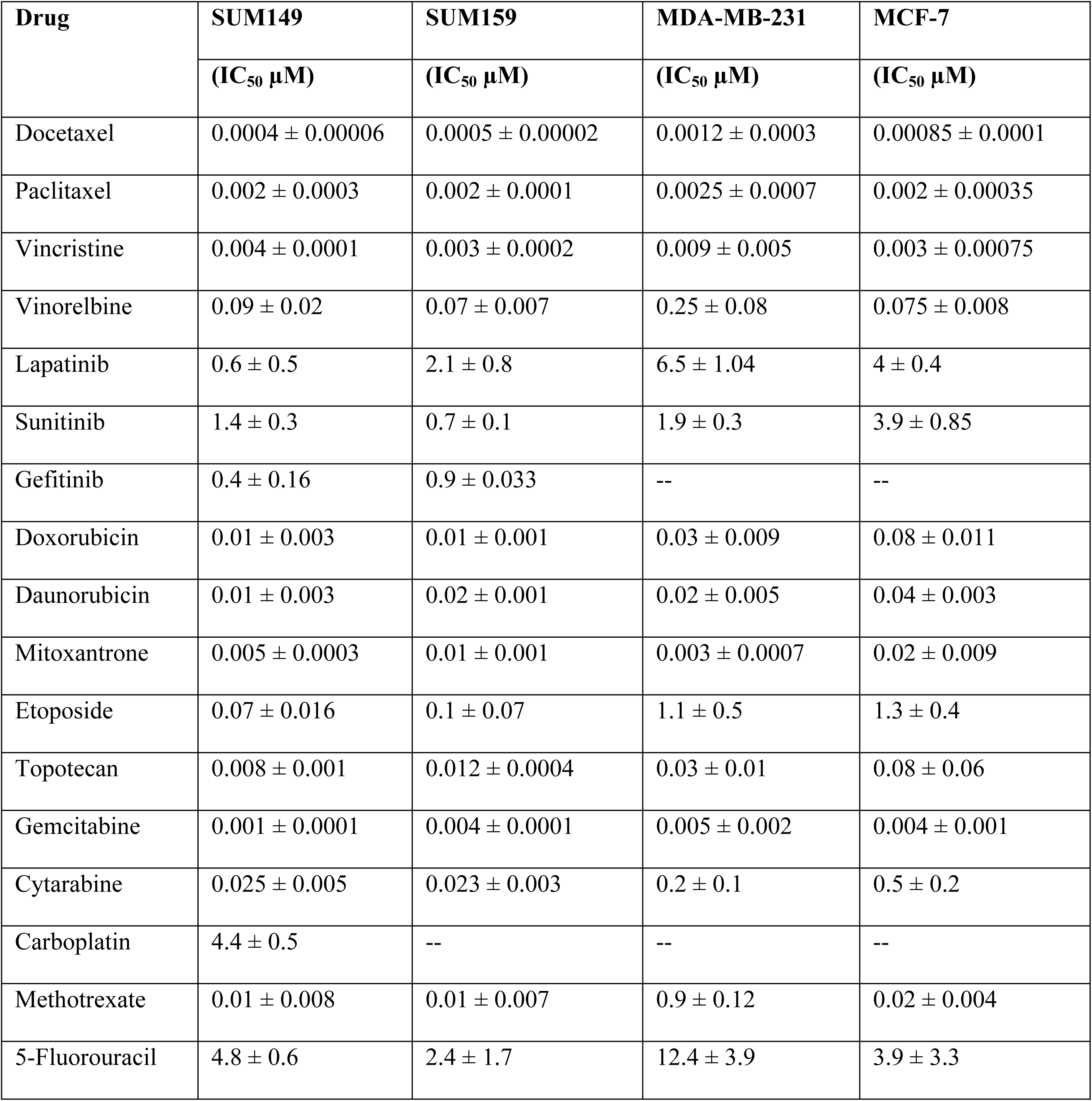
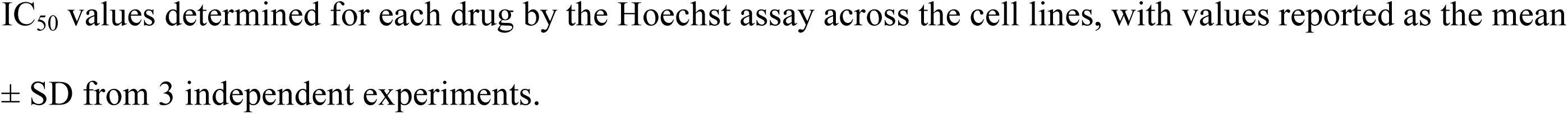
Efficacy of LWAS-predicted drugs and compounds in four breast cancer cell lines: SUM149, SUM159, MDA-MB-231, and MCF-7.

#### Identification of GRR candidates with effects on cell proliferation

In our previous GRR analysis, we used a compiled IBC-gene expression signature (IBC-GES) to query the LINCS database for drugs that significantly reversed expression of the majority of IBC-GES genes and identified 19 drugs [28]. Here, we evaluate the anti-proliferative activity of these GRR candidates (16 of the 19 were commercially available), using the Hoechst assay in the same four cell lines described above. The GRR candidates showed differential anti-proliferative activity in each cell line, with significant effects (IC_50_ < 10 μM) observed for 6 compounds in SUM149, SUM159 and MDA-MB-231, and 5 in MCF-7 cells (represented as heatmaps in **Fig 2A**). Although the GRR list comprises both oncology and non-oncology compounds from diverse drug classes (**S2 Table**), the active compounds were primarily tyrosine kinase inhibitors, with Akt Inhibitor-IV (AKTIV) being an exception. In SUM149, tipifarnib displayed the highest potency (absolute IC_50_ = 0.075 μM), followed by tyrphostin AG-1478 and AKTIV, which showed comparable potency (absolute IC_50_ ≈ 0.2 μM). Tipifarnib showed a biphasic DRC, and upon calculating the relative IC_50_ in SUM149, it shifted to 7 ± 0.2 nM (10-fold increase in potency). Tyrphostin AG-1478 showed a low efficacy curve (indicating antiproliferative activity), its relative IC_50_ in SUM149 shifted to 0.05 ± 0.01 μM (4-fold increase in potency). Temsirolimus displayed varying potency across the 4 cell lines with an absolute IC_50_ of 4.3 μM in SUM149, compared to much higher potency in SUM159, MDA-MB-231, and MCF-7 cell lines (IC_50_ range: 0.2-0.6 nM). Temsirolimus also showed a biphasic DRC in SUM149 cells, with a relative IC_50_ value of 0.1 nM, which is comparable to other cell lines but with lower efficacy (approximately 60% maximum inhibition). MDA-MB-231 showed unique sensitivity to nicardipine (projected IC_50_ = 16.7 μM), while showing reduced sensitivity to the other compounds compared to other cell lines. Tyrphostin AG-1478 showed no inhibitory activity in MCF-7, unlike other cell lines. Representative DRCs for SUM149 are shown in **Fig 2B**, while DRCs for SUM159, MDA-MB-231, and MCF-7 cell lines are in **S2 Fig**, showing a range of responses. The IC_50_ values for the GRR’s compounds are summarized in **Table 2**.

**Fig 2.**
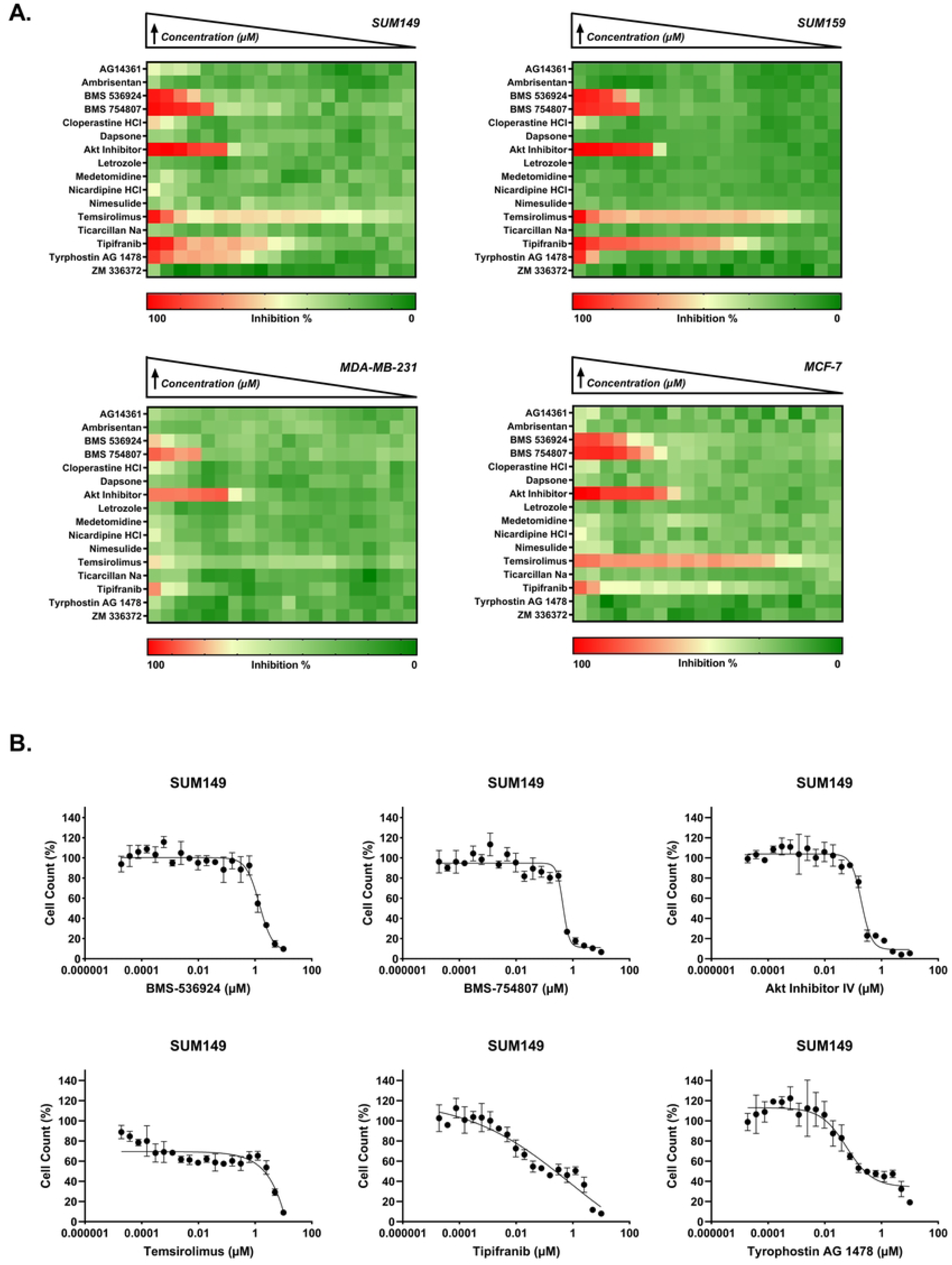
GRR drug sensitivity profiles in IBC and non-IBC breast cancer cell line panel by Hoechst assay. (A) Heatmaps representing the percent inhibition response for a 2-fold concentration dose-response starting at 10 µM for SUM149, SUM159, MDA-MB-231, and MCF-7. Drug names are listed on the y-axis. The color scale indicates the level of inhibition, with red representing high inhibition (close to 100%) and green representing low inhibition (close to 0%). (B) Representative dose-response curves for active inhibitors BMS-536924, BMS-754807, Akt Inhibitor IV, temsirolimus, tipifarnib, and tyrphostin AG 1478 in the SUM149 cell line, showing the percent cell count relative to control against drug concentration (µM). Data points and error bars represent the mean ± SD. DRCs for SUM159, MDA-MB-231, and MCF-7 are in **S2 Fig**.

**Table 2.**
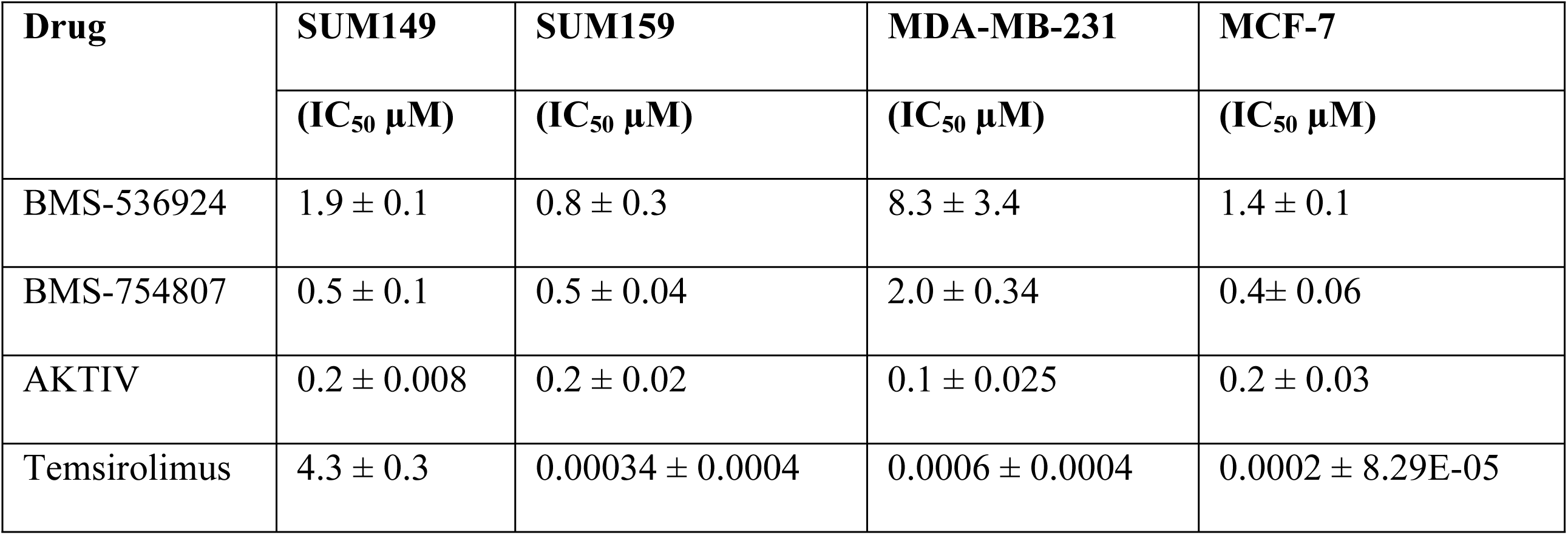
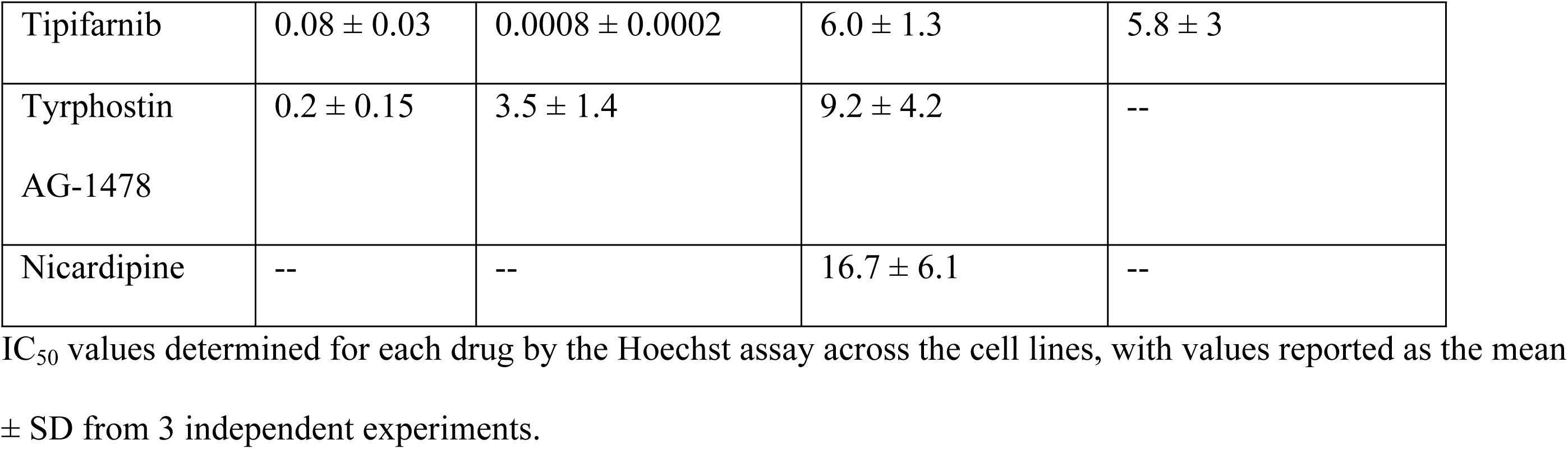
Efficacy of the GRR-predicted drugs and compounds in four breast cancer cell lines: SUM149, SUM159, MDA-MB-231, and MCF-7.

#### Identification of LWAS and GRR candidates with efficacy in the MTT cell viability assay

We also assessed the LWAS and GRR compounds in an orthogonal cell-based assay (MTT) to assess effects on cell viability. Testing of the compounds in the MTT assay yielded IC_50_ values and DRCs comparable to those of the Hoechst assay for both the LWAS (**S3 Fig**) and the GRR (**S4 Fig**) compounds. All the IC_50_ values for the LWAS drugs and GRR compounds in the MTT assay across the 4 cell lines are summarized in **S3 Table**. For the MTT assay, using an activity threshold of IC₅₀ < 10 µM, the LWAS list screening yielded 17 active compounds in SUM149, 16 in SUM159, 15 in both MDA-MB-231 and MCF-7, whereas the GRR panel yielded 6 anti-proliferative compounds in SUM149, SUM159 and MDA-MB-231, and 5 in MCF-7. The same compounds were active in both the Hoechst and MTT assays and maintained the same overall rank-order of potency. For LWAS compounds tested in the MTT assay, microtubule inhibitors again were the most potent class with nanomolar potency in both assays across all cell lines, and the median docetaxel IC₅₀ shifts by <1.5-fold (e.g. 0.0004 µM in SUM149 by Hoechst versus 0.0003 µM by MTT). Likewise, topoisomerase inhibitors retain mid-nanomolar potency, while antimetabolites gemcitabine and cytarabine showed only ≤ 2-fold differences between assays. For the GRR list, the potencies for each compound in both assay formats were comparable, in the range of 1- to 3-fold difference. In SUM149, temsirolimus showed an IC_50_ value of 4.3 µM in the Hoechst assay and 7.3 µM in MTT assay.

For AKTIV, BMS-536924 and BMS-754807, the fold shifts are generally below 3, except for a 7-fold change for BMS-754807 in SUM149 (IC_50_: 3.6 µM in MTT assay 0.5 µM in Hoechst assay). Tyrphostin AG-1478 shows comparable nanomolar activity in SUM149 and comparable micromolar activity SUM159 cell lines. A couple of results were not consistent between the two assay formats; tipifarnib in SUM159, where the IC_50_ was 0.8 nM by Hoechst compared to 2.1 µM by MTT, and gefitinib, which was ∼10-fold less potent in SUM159 by MTT (IC_50_: 11 µM) than by Hoechst (IC_50_: 0.9 µM). These differences might reflect the mechanism behind each assay where cytostatic effect manifests faster in the nuclear-count readout than in the metabolic assay. But the overall strong concordance between the two assays was about 85 %: individual IC_50_ pairs differed by less than 2-fold, indicating that the viability (MTT) and nuclear count (Hoechst) endpoints read out very similar anti-proliferative effects for most compounds, with SUM149 being the most responsive cell line in both formats.

### Assessing drug combinations in SUM149 for synergistic effects

The potential synergistic effects of drug combinations were assessed in the SUM149 cell line as this was our representative IBC model. The SUM149 cell line was derived from an IBC patient tumor [40], and is a well-established model for IBC [33, 34, 41–43]. Our Hoechst and MTT efficacy profiling of single compounds identified the most potent compounds from both the LWAS and GRR lists for subsequent combination studies. These drug-combination experiments were conducted using a two-tiered approach that included primary and secondary screening. In the primary screening, every two compounds were combined and tested at their respective EC_25_ values. The EC_25_ value was chosen to achieve a modest effect on cell viability, thereby minimizing toxicity and avoiding one drug overwhelming the response. In this case, each drug can affect cellular activity without killing most cells, creating an opportunity to observe an extra effect when the drugs are combined [44]. The 17 active LWAS inhibitors were reduced to 13 by excluding less potent compounds with similar mechanisms of action (MOA), yielding a total of 78 unique two-drug combinations. Likewise, combining the 6 GRR inhibitors gave 15 unique two-drug combinations. The Bliss excess scores [35] were calculated, and a one-sample t-test was used to determine the statistical significance. Combinations that showed statistical significance were subsequently validated in the secondary combination screening phase using 8×8 dose-response matrices in 2D cell models and then confirmed in 3D spheroid models to enable a more comprehensive assessment. Synergy evaluation was performed using SynergyFinder3 [38] and SynergyFinder+ [39] online tools. The complete experimental workflow is summarized in **Fig 3**.

**Fig 3.**
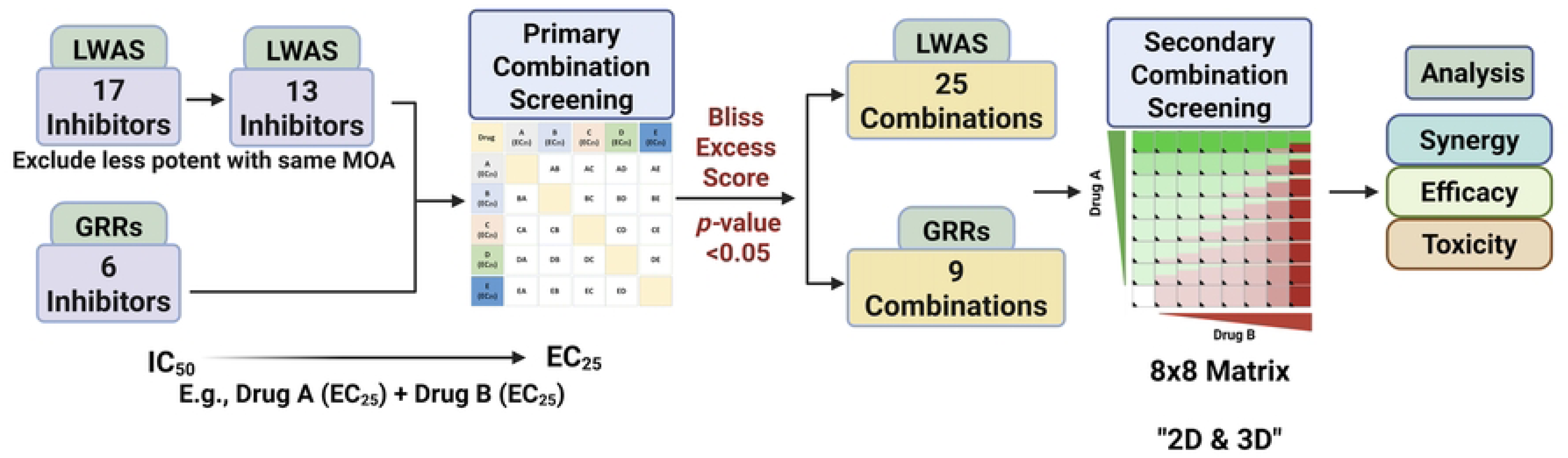
Workflow for identification and evaluation of synergistic drug combinations. The screening process began with 17 LWAS inhibitors refined to 13 LWAS inhibitors after excluding less potent compounds with similar mechanisms of action (MOA) plus 6 GRR inhibitors. Primary drug combination screening was done using drug pairs at their respective EC_25_ values. The identified combinations were selected based on Bliss excess scores and statistical significance (*p* value < 0.05), which identified 25 LWAS and 9 GRR synergistic combinations to be confirmed in the secondary screening phase, utilizing an 8×8 matrix format. The final analysis included synergy assessment and efficacy evaluation in 2D and 3D cell models.

#### Identification of key combinations using single-dose primary screening

Our initial primary screening analysis, while using only single EC_25_-dose combinations, revealed potential drug synergy based on Bliss excess score calculations, in which negative values indicate a synergistic effect when starting with cell viability data [35]. Quantitative analysis of the primary combination screen identified 25 synergistic combinations from the LWAS list (**Fig 4A**, 32.1% success rate, p < 0.05) and 9 synergistic combinations from the GRR list (**Fig 4B**, 60% success rate, p < 0.05). These synergistic combinations demonstrated robust negative excess Bliss scores ranging from −0.16 to −0.27.

**Fig 4.**
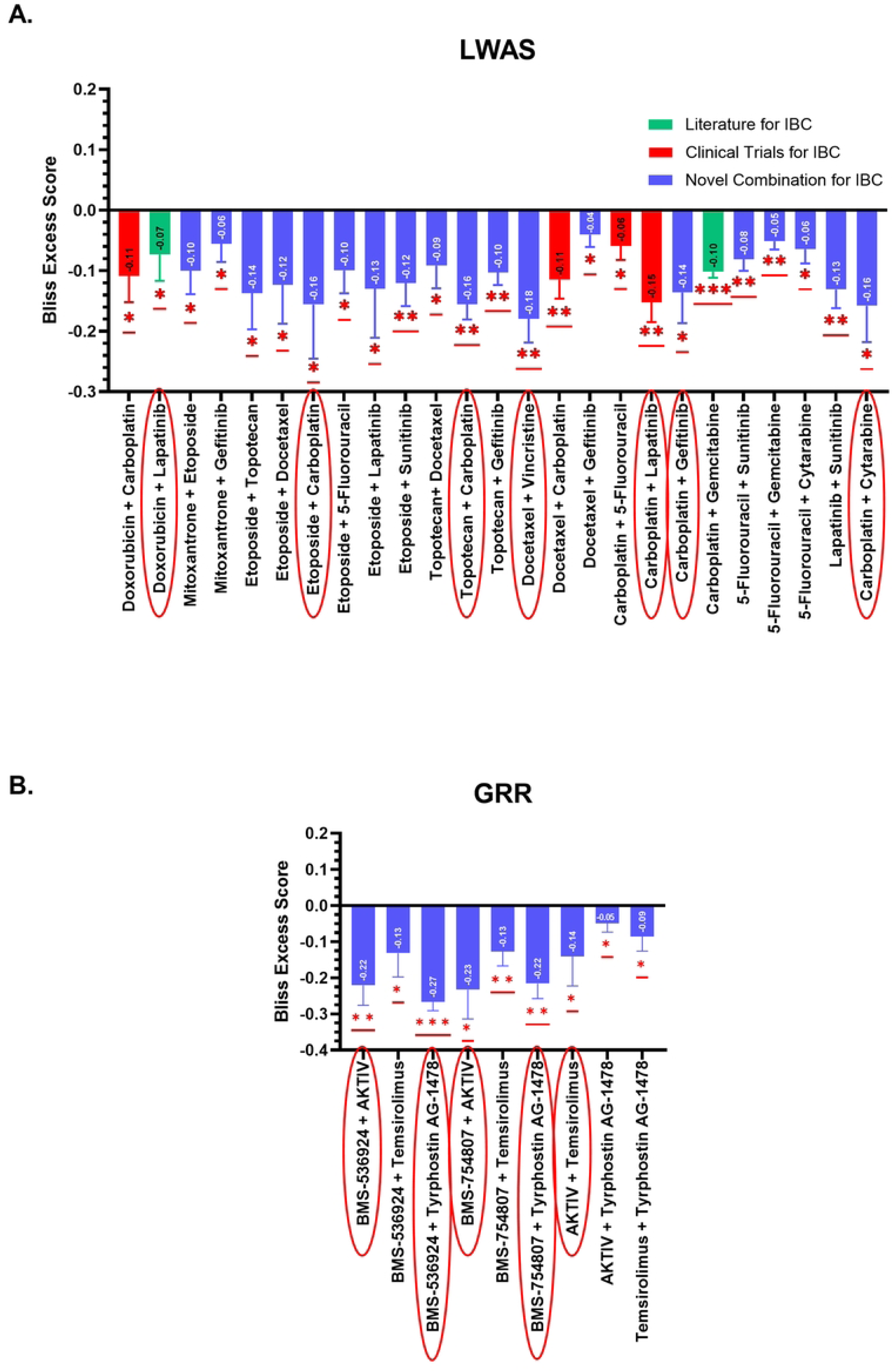
Drug combinations identified from primary combination screening based on Bliss excess scores in SUM149. Primary drug combination screening was done using drug pairs at their respective EC_25_ values. Each combination is evaluated based on percentage viability, with negative scores indicating synergism. (A) Twenty-five significantly different combinations were identified from the LWAS study and categorized as reported in the literature for IBC (green bars), tested in clinical trials for IBC (red bars), and novel combinations for IBC (blue bars). (B) Nine significantly different combinations were identified from the GRR list, all of which were novel for IBC. Red asterisks indicate statistically significant deviations centered at zero based on a t-test (*p < 0.05; **p < 0.01; ***p < 0.001). Red oval shapes highlight the selected combinations for further screening in the 8×8 matrix.

For the LWAS primary combination results, 4 of the 25 combinations (red bar color in **Fig 4**) have been previously documented in IBC-related clinical trials registered on ClinicalTrials.gov, including where carboplatin was combined with either doxorubicin (NCT05415215), docetaxel (NCT00251329 and NCT00118053), lapatinib (NCT01426880), or 5-fluorouracil (NCT01036087). These trials either included IBC patients or specifically focused on IBC populations. Further, the combination of carboplatin with gemcitabine is listed as a treatment option for IBC [45]. For the GRR primary combination screen, combinations targeting PI3K/AKT/mTOR downstream pathways, either directly or indirectly, showed high synergy scores, including BMS-536924 (a dual IGF-1R/IR tyrosine kinase inhibitor) with tyrphostin AG 1478 (a selective EGFR tyrosine kinase inhibitor); and BMS-536924 with AKTIV (an inhibitor of Akt phosphorylation). To our knowledge, the remaining LWAS combinations (bars highlighted in blue) and all the GRR pairs represent potential novel therapeutic strategies for IBC.

Drug combinations with the highest negative excess Bliss scores (highlighted by red oval shapes in **Fig 4**, along with two previously reported were selected for secondary combination screening and evaluated using 8×8 dose-response matrices.

#### Determination of synergy score based on dose-response combination matrix

For our secondary screening, we conducted an 8×8 dose-response matrix analysis (**Fig 3**) to validate promising drug combinations identified from the primary single-dose combination screening. This approach again focused on evaluating drug synergy for SUM149 cell proliferation. For each drug pair (7 for LWAS and 5 for GRR), we designed dose matrices with IC_50_ concentrations positioned centrally, with concentration ranges above and below IC_50_ at ½-log intervals (**S4 Table**). The experimental protocol involved treating SUM149 cells with drug combinations for 72 h, followed by Hoechst dye-based cell counting. Based on the results from our primary combination screening, we selected the top five combinations from both LWAS and GRR lists. Additionally, we included two other combinations from the LWAS list: one documented in the literature [46] and one found in clinical trials [47] (see **Fig. 4**). We conducted a comprehensive synergy and efficacy analysis to identify the optimal combinations. The synergy score calculation is based on four widely used reference models: Bliss, Loewe, HSA, and ZIP [48]. According to the framework proposed by Malyutina *et al* [49], our combinations were classified as synergistic (scores > 5) or antagonistic (scores < −5) when meeting these thresholds across all models as a strict criterion. For the efficacy metric evaluation, we used the Combination Sensitivity Score (CSS) developed by Malyutina *et al* [49]. CSS quantifies combination efficacy against cell proliferation by calculating the mean area under the dose-response curve when titrating one drug against the IC_50_ concentration of the second drug (CSS_1_IC50_2_ and CSS_2_IC50_1_). SynergyFinder+ was used to compute the average CSS values for all combinations. In the LWAS combination analysis, we found that none met the strict criterion for overall synergy or antagonism (all the overall synergy (δ) scores fell between −5 and 5 across all reference models, indicating additive effects) (**S5 Fig**). A closer examination of the contour plots was then undertaken, focusing on the Most Synergistic Area (MSA) [38], which represents a 3×3 dose window in the combination matrix that yields the highest positive deviation from the chosen reference model [38]. This analysis of MSA δ scores gave (δ > 5) for five (Loewe model) and seven (HSA model) of the combinations, respectively (**S5 Fig**, **Table 3**). This approach can highlight the dose ranges where the interaction is most pronounced and help in selecting concentration pairs for follow-up validation. For example, we observed that the lapatinib + doxorubicin combination gave scores of 26.2 for Loewe and 7.5 for HSA, whereas lapatinib + carboplatin showed scores of 18.6 in Loewe and 10.1 in HSA. However, for the Bliss and ZIP models, MSA δ scores for all these were < 5, reducing confidence that these combinations might be synergistic.

**Table 3.**
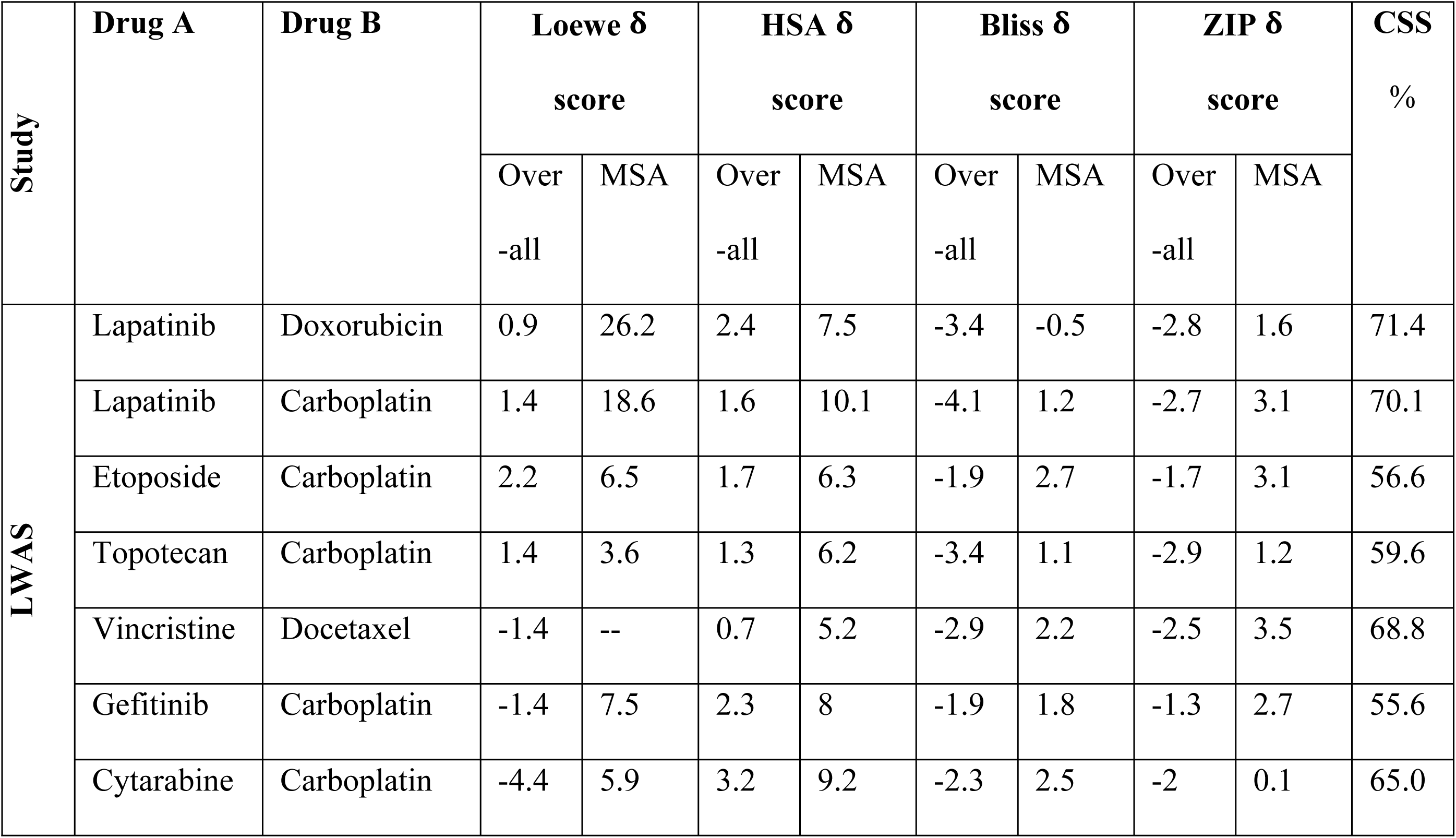
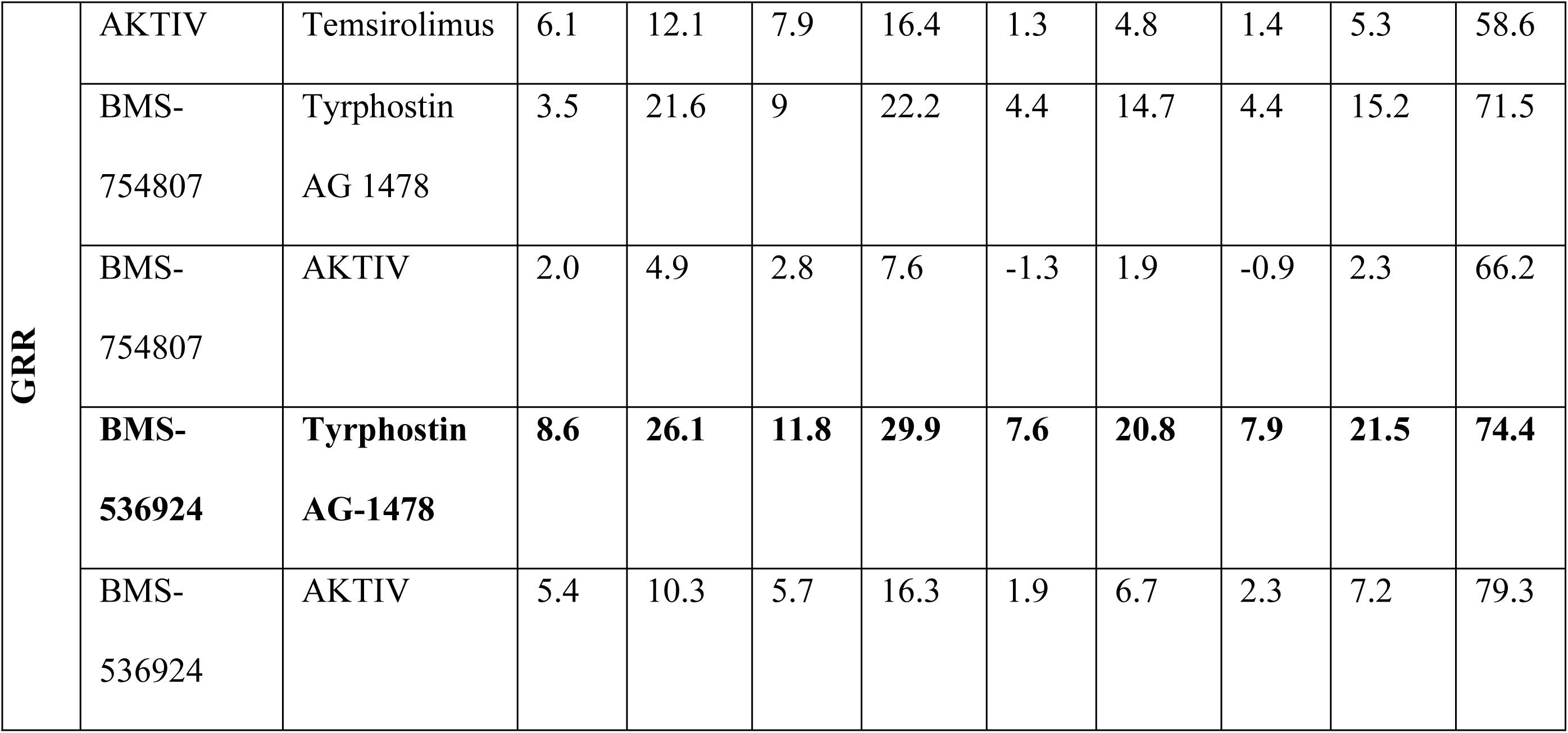
Synergy scores for the secondary drug combinations (8×8 Matrix) in 2D cell models across four reference models.

In the GRR combination analysis, the BMS-536924 and tyrphostin AG-1478 combination exceeded the 5-score threshold at both the overall (ZIP: 7.9, HSA: 11.8, Bliss: 7.6, Loewe: 8.6) and MSA (ZIP: 21.5, HSA: 29.9, Bliss: 20.8, Loewe: 26.1) δ scoring levels, with a 74.4% CSS score for efficacy indicating substantial therapeutic potential through dual pathway inhibition. Three other combinations (BMS-536924 + AKTIV; temsirolimus + AKTIV; and BMS-754807 + tyrphostin AG-1478). showed synergistic effects at the MSA level (exceeding the 5-score threshold), while only one showed (BMS-754807 + AKTIV) additive effects. The contour plots of the 2D drug combination experiments are shown in **Fig 5**. The corresponding overall synergy (δ) scores and MSA δ scores from SynergyFinder3 for all the GRR combinations are shown in **Table 3**. All the remaining LWAS and GRR combinations not shown had δ <5 for all models at both the overall and MSA level.

**Fig 5.**
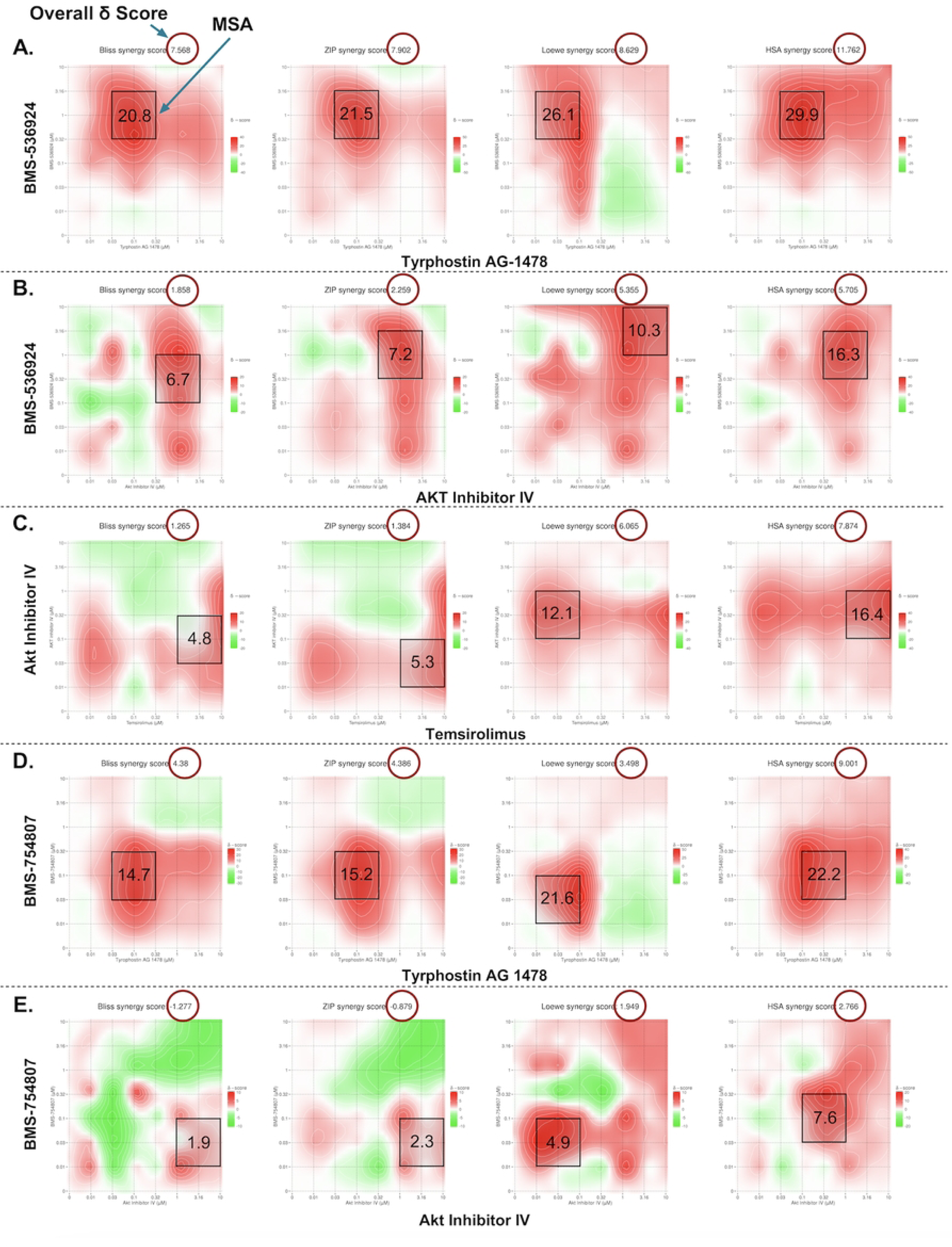
Contour plots of selected GRR drug combinations. Synergy scores across Bliss, ZIP, Loewe and HSA reference models were obtained from SynergyFinder3. (A) BMS-536924 + tyrphostin AG 1478, (B) BMS-536924 + AKT Inhibitor IV, (C) Temsirolimus + Akt Inhibitor IV, (D) BMS-754807 + tyrphostin AG 1478, and (E) BMS-754807 + Akt Inhibitor IV. In each plot, x and y axes represent drug concentrations. Red areas indicate synergy, while green areas indicate antagonism. Black boxes highlight the MSA, with scores shown. Red circles in the top right corner of each plot indicate the overall δ score. Color scales on the right of each plot show the range of synergy scores. The data represented the average of three independent experiments (n = 3).

### Evaluation of selected GRR drug combinations for effects on SUM149 spheroids

To more accurately mimic IBC tumors, we utilized SUM149 spheroids as a 3D tumor model [33, 50]. In this assay, the SUM149 spheroids were allowed to form over 72 h before adding each of the three selected drug combinations identified from the 2D study. After an additional 72 h of treatment, cell viability was then evaluated using CellTiter-Glo, an ATP-based luminescence assay. To ensure consistency and allow direct comparison between 2D and 3D experiments, we used the same 8×8 dose-response matrix that was used in our 2D experiments.

In our 3D spheroid study the BMS-536924 and tyrphostin AG 1478 combination yielded overall synergy scores of (ZIP: 7.1, HSA: 12.2, Bliss: 6.9, and Loewe: 9.9), while MSA level scores were (ZIP: 11.8, HSA: 23.9, Bliss: 12.1, and Loewe: 17.9). BMS-536924 and AKTIV combination showed overall synergy scores of (ZIP: 4.1, Loewe: 0.5, Bliss: 3.4, and HSA: 7.8), with MSA level scores reaching (ZIP: 10.7, Loewe: 6.4, Bliss: 10.5, and HSA: 22.9). Lastly, for the temsirolimus and AKTIV combination, overall synergy scores were (ZIP: 6.1, Loewe: −8.2, Bliss: 5.8, and HSA: 7.7), with MSA level scores of (ZIP: 18.9, Loewe: 5.2, Bliss: 18.2, and HSA: 19.9). The corresponding contour plots are shown in **Fig 6**. Based on the accepted criteria of a synergy score exceeding 5 across all reference models [49], these results indicate a synergistic effect at the MSA level for all tested combinations. Notably, as in the 2D study, the BMS-536924 and tyrphostin AG 1478 combination also achieved synergy at the overall level in the 3D study. It is worth noting that the Loewe model yielded a negative synergy score for the combination of temsirolimus and AKTIV. This result may be attributed to the fact that the Loewe model relies on well-defined dose-response curves for each agent [51, 52], a condition not satisfied by temsirolimus in our 3D assay, where it failed to produce a consistent dose–response relationship.

**Fig 6.**
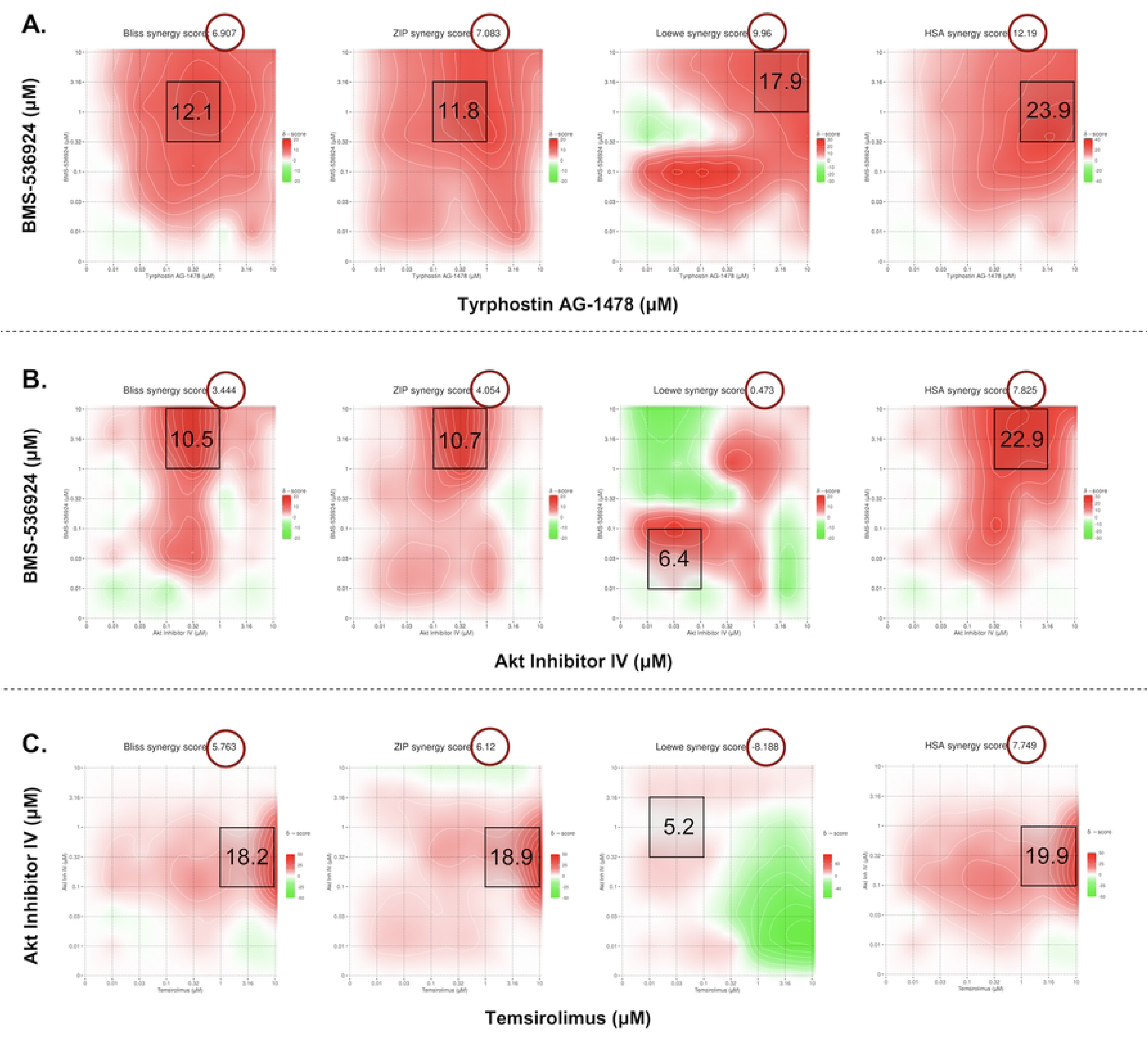
Analysis of drug combinations in 3D SUM149 spheroid model. Contour plots and their synergy scores across Bliss, ZIP, Loewe and HSA reference models obtained from SynergyFinder3. (A) BMS-536924 + tyrphostin AG 1478 (B) BMS-536924 + AKT Inhibitor IV and (C) Temsirolimus + AKT Inhibitor IV combinations. In each plot, x and y axes represent drug concentrations. Red areas indicate synergy, while green areas indicate antagonism. Black boxes highlight the MSA, with scores shown. Red circles in the top right corner of each plot indicate the overall δ score. Color scales on the right of each plot show the range of synergy score. The data are represented as the average of three independent experiments (n = 3).

SynergyFinder+ was used to analyze the synergy scores and inhibition percentages (%) based on percent-effect of DMSO control at specific concentrations. There was good agreement between our 3D and 2D results, with comparable synergy scores across the four reference models (**Table 4**). For the inhibition comparison, the BMS-536924 (1 µM) and tyrphostin AG 1478 (0.1 µM) combination showed nearly identical % inhibition in both 2D (78.6%) and 3D (79.1%) models. The BMS-536924 (1 µM) and AKTIV (1 µM) combination was more effective in 3D, achieving 92.3% inhibition compared to 79.4% in 2D. The temsirolimus (10 µM) and AKTIV (0.1 µM) combination achieved 61.2% inhibition in the 3D model, which is comparable to the 54.2% inhibition observed in 2D (**S6 Fig**). The 3D results also revealed stronger inhibitory effects from the combinations than from single agents. The combination of BMS-536924 (1 µM) with Tyrphostin AG 1478 (0.1 µM) achieved 79.1% inhibition, which is higher than each agent alone (BMS-536924: 66.6%; tyrphostin AG-1478: 3.7%). Similarly, combining BMS-536924 (1 µM) with AKTIV (1 µM) resulted in 92.3% inhibition, surpassing the effects of individual treatments (BMS-536924: 65.1%; AKTIV: 73.1%). While the temsirolimus (10 µM) and AKTIV (0.1 µM) combination showed the lowest inhibition (61.2%), it still demonstrated improved efficacy compared to single-agent treatments (temsirolimus: −12.9%; AKTIV: 16.4%). Although the synergy metrics yielded high scores for this combination (**Table 4**), those values can be misleading in the 3D context as these calculations assume both drugs retain intrinsic activity, where their combined effect should be greater than the additive expectation. However, temsirolimus showed minimal to no activity in 3D; therefore, this interaction might be better described as enhancement or potentiation of AKTIV activity, which increased growth inhibition from 16.4 % with AKTIV alone to 61.2 % in combination.

**Table 4.**
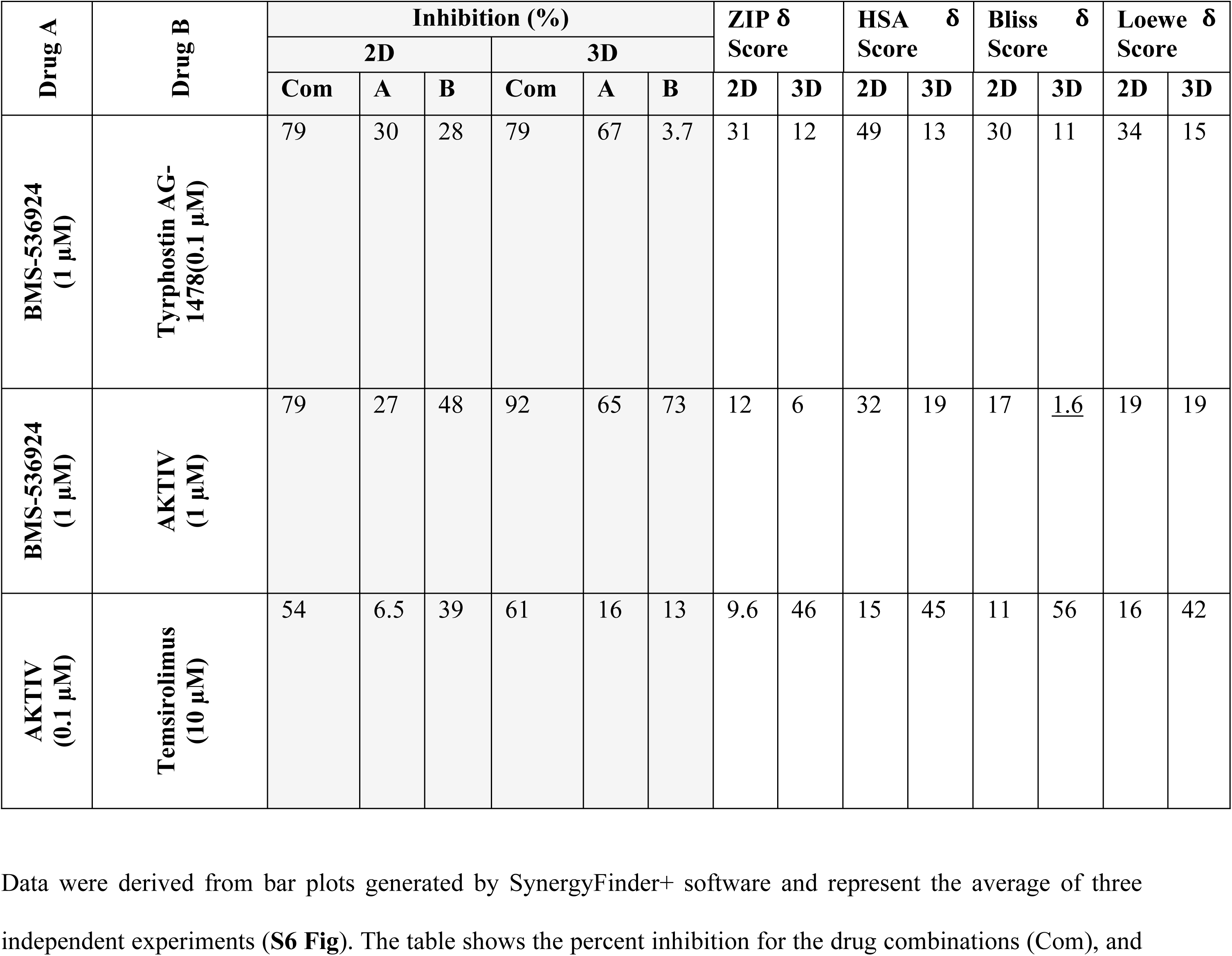
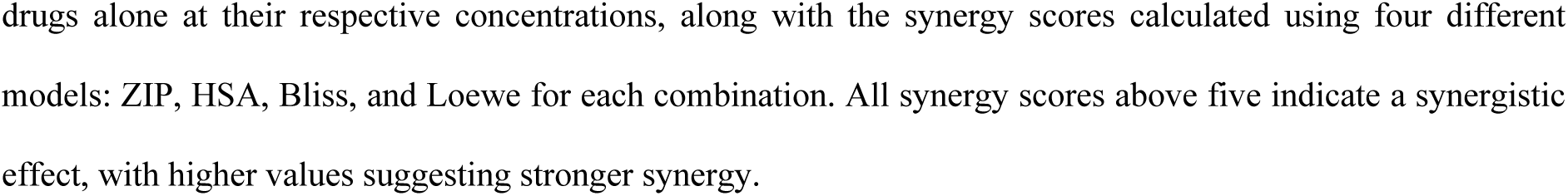
Summary table of synergy scores in SUM149 2D and 3D cell models.

To further understand the effects of drug combinations on SUM149 spheroids, we used the SYTOX™ Red Dead Cell Stain to visualize the spheroids using the IncuCyte system. In all experimental panels, 0.2% DMSO was used as a negative (solvent) control, while staurosporine (10 μM) was used as a positive inhibitor control [50], which, as expected, caused significant spheroid disruption after 72 hours of treatment. The effects of individual drugs and their combinations are shown in **Fig 7**. Some combinations, especially those involving BMS-536924, appear to induce more pronounced changes in spheroid morphology compared to single-agent treatments. The images reveal varying degrees of spheroid integrity and morphological alterations in response to combination treatment.

**Fig 7.**
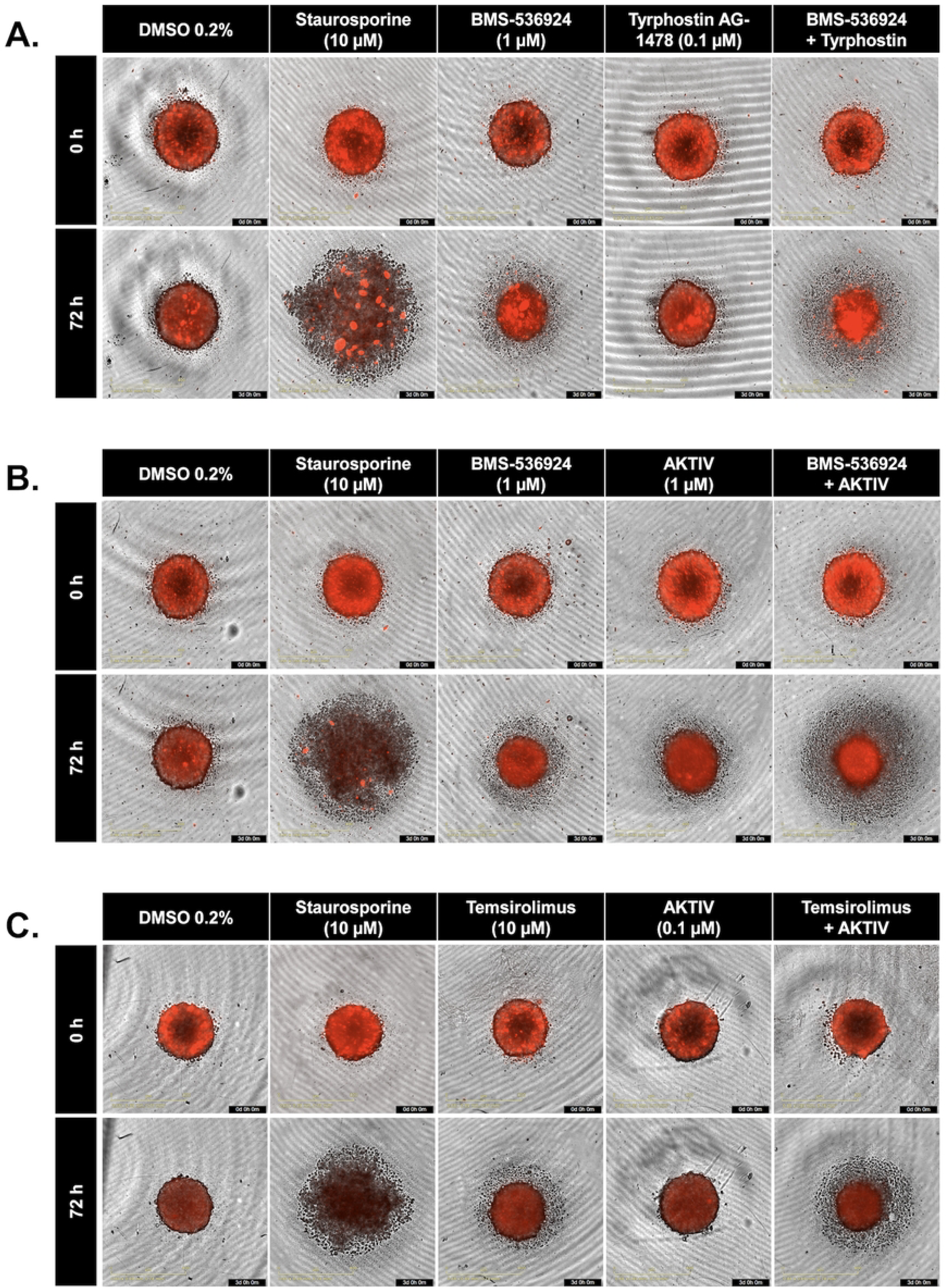
Representative SUM149 spheroids showing the effects of individual drugs and drug combinations. (A) BMS-536924 (1 µM) and tyrphostin AG-1478 (0.1 µM) combination. (B) BMS-536924 (1 µM) and AKTIV (1 µM) combination. (C) Temsirolimus (10 µM) and AKTIV (0.1 µM) combination. In each set, the top row shows spheroids at 0 h, while the bottom row shows the same spheroids after 72 h of treatment. DMSO 0.2% represents a negative control while staurosporine (10 μM) was used as a positive control.

Taken together, the drugs we identified from both the 2D and 3D combination screening pipeline included inhibitors BMS-536924 [53] targeting IGF1-R, tyrphostin AG-1478 [54, 55] targeting EGFR, Akt inhibitor IV [56] targeting Akt phosphorylation, and temsirolimus [57] targeting mTOR.

### Assessment of selected GRR drug combinations on SUM149 motility

To evaluate the identified drugs for effects on cell migration, a critical parameter in cancer cell behavior and metastasis, we used the “wound healing” assay, utilizing IncuCyte’s precision WoundMaker tool. For this assay, a wound is created within a confluent cell monolayer, and cell movement back into the wound measured in real time using automated IncuCyte imaging. Cells were plated overnight and then the “wound” was created, followed by selected combination drug treatments (as listed in **Table 4**). Relative Wound Density (RWD) %, which considers any effects of cell proliferation, was calculated to quantify cell movement into the wound area relative to initial wound size over 24 h.

The DMSO vehicle control demonstrated progressive wound closure over the 24 h observation period (shown in **Fig 8**). SUM149 treatment with the IGF1-R inhibitor BMS-536924 (1 μM) did not result in any significant effect on migration kinetics compared to control (*p* > 0.05), while treatment with the mTOR inhibitor tyrphostin AG-1478 (0.1 μM) had a significant effect on slowing wound closure with *p* < 0.05. When both BMS-526924 and tyrphostin AG-1478 were combined at those concentrations, migration was significantly reduced by 30% at 24 h (*p* < 0.001). The combination treatment was not significantly different from tyrphostin AG-1478 treatment alone. SUM149 cells treated with the Akt inhibitor AKTIV (1 μM) had significantly impaired wound closure, reducing migration by > 50% compared to control at 24 h (p < 0.001). When both BMS-526924 and AKTIV were combined at these concentrations, migration was reduced by 30% at 24 h (*p* < 0.001). The combination treatment was not significantly different from AKTIV treatment alone. Treatment of SUM149 cells with temsirolimus (1 μM), an mTOR complex 1 inhibitor, significantly reduced migration by ∼ 40% relative to control at 24 h (p < 0.05). The combination of temsirolimus and low dose AKTIV (0.1 μM) did produce significant effects (p < 0.001) compared to the control and single treatments. Representative wound healing images and corresponding statistical analyses are shown in **Fig 8**, with each experimental condition performed in eight replicates (n = 8).

**Fig 8.**
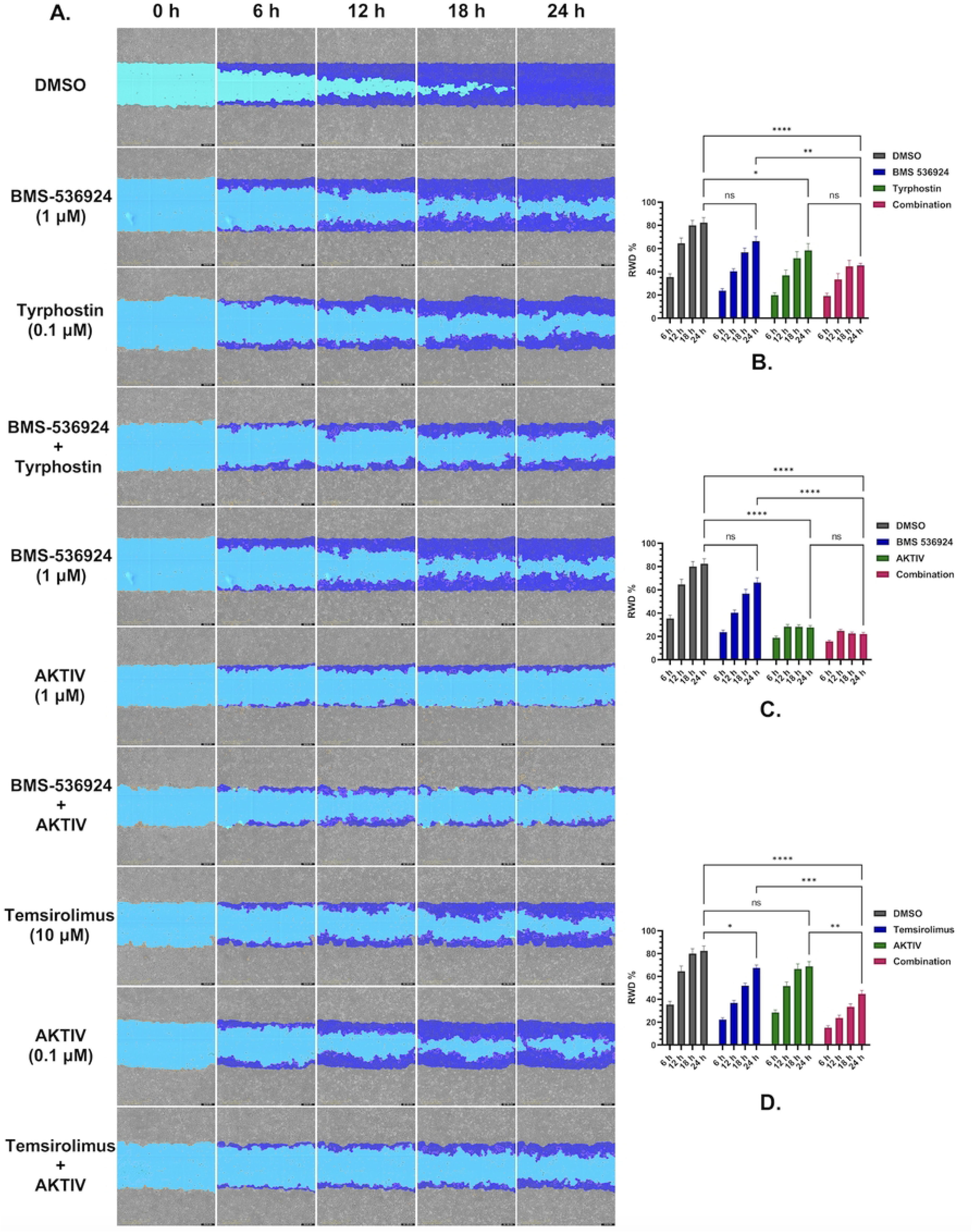
Effect of GRR selected drugs on SUM149 motility. Relative wound density (RWD %) was measured at 6 h intervals from 0 to 24 h. The effects of individual drugs and their combinations are compared to the DMSO control. (A) Representative images of wound closure over 24 h treatment for each condition. (B) to (D) Quantification of the RWD % relative to each combination. (B) BMS 536924 (1 µM) and tyrphostin AG 1478 (0.1 µM). (C) BMS 536924 (1 µM) and AKTIV (1 µM). (D) Temsirolimus (10 µM) and AKTIV (0.1 µM). Statistical significance is indicated by asterisks (*p < 0.05, **p < 0.01, ***p < 0.001, ****p < 0.0001, ns: not significant). Error bars represent mean ± SEM from eight technical replicates.

### Selected GRR combinations mediate the reduction of phosphoprotein expression

We next evaluated the effects of the selected three drug combinations on key signaling proteins in SUM149 cells. The analysis focused on expression levels and phosphorylation status of Akt, EGFR, IGF1-R, and mTOR proteins in response to single- and combination-drug treatments. Western blot analysis was performed after 24 h of drug treatment, and densitometric analysis for all treatment conditions, including individual drugs and corresponding combinations, and representative blots are presented in **Fig 9**. Densitometric analyses of triplicate experiments (representative whole blots shown in **S7 Fig**) confirmed the statistical significance of these phosphorylation changes, with β-actin serving as the loading control across all conditions. The maintenance of total protein levels across treatments suggests that the observed effects were specific to protein phosphorylation rather than overall protein expression. Due to technical challenges using the Li-COR system for IGF1-R and pIGF1-R, HRP-tagged secondary antibodies and iBRIGHT imaging was used for these proteins (**S7 Fig**). All four of the proteins targeted by our selected GRR combinations were detectable in the SUM149 IBC cell line by Western blot (**Fig 9**), and the drug combinations effectively suppressed their target pathways (**Fig 9** and **S7 Fig**). The combination of AKTIV (0.1 µM) with temsirolimus (10 µM) demonstrated the strongest inhibitory effects, achieving a significant reduction of both p-Akt (*p* < 0.001) and p-mTOR (*p* < 0.01) (**Fig 9A**). The second combination, AKTIV (1 µM) with BMS-536924 (1 µM), showed a significant decrease in p-Akt levels (*p* < 0.01) (**Fig. 9B**), which appeared to be primarily driven by BMS-536924 rather than AKTIV. The combination of tyrphostin AG-1478 (0.1 µM) and BMS-536924 (1 µM) resulted in a reduction in both phosphorylated EGFR (p-EGFR) levels and phosphorylated IGF1-R (p-IGF1-R), but did not meet statistical significance, while maintaining stable total protein levels for EGFR and IGF1-R (**Fig. 9C**).

**Fig 9.**
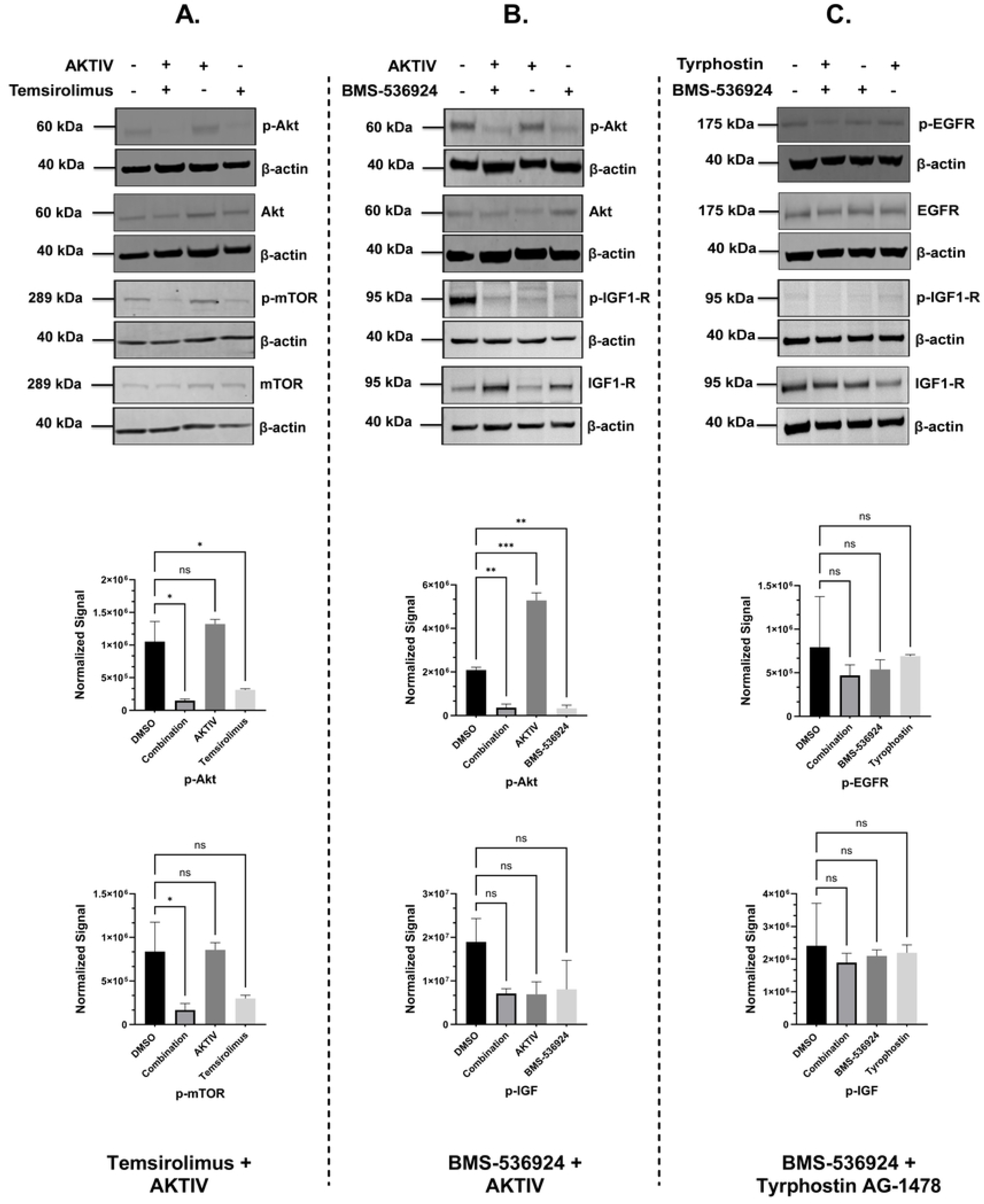
Western blot analysis of protein expression levels in response to the tested drugs and combinations. Representative Western blots for (A) Temsirolimus (10 µM) and Akt inhibitor IV (0.1 µM) combination effects on the expression of p-Akt, total Akt, p-mTOR, and total mTOR; (B) Akt Inhibitor IV (1 µM) and BMS-536924 (1 µM) combination effects on the expression of p-Akt, total Akt, p-IGF1-R, and total IGF1-R; and (C) Tyrphostin AG-1478 (0.1 µM) and BMS-536924 (1 µM) combination effects on the expression of p-EGFR, total EGFR, p-IGF1-R, and total IGF1-R. The presence (+) or absence (-) of each treatment is indicated, with β-actin serving as a loading control. Quantification of DMSO bands vs each treatment bands. Representative whole-blot images of triplicate samples (n = 3) are shown in **S7 Fig**.

### Assessment of differential gene expression changes in SUM149 cells treated with the selected GRR combinations

To assess if the selected GRR combinations we identified herein with efficacy in multiple IBC cell models significantly reversed IBC gene expression, we undertook a RNA Seq analysis. For this, SUM149 cells were cultured and treated for 24 h (n = 3) with the selected GRR combinations, as well as DMSO vehicle control. RNA was isolated and RNA-Seq conducted as in materials and methods. All three combination treatments caused significant global gene expression changes in SUM149 (see heatmaps and volcano plots in **S8 Fig**). Analysis of the expression of the 297 IBC-GES genes only in SUM149 cells treated with the three dual-drug combinations revealed distinct transcriptional reversal patterns. For those IBC-GES genes detected as significantly expressed (*p*adj < 0.05) in SUM149 cells (blue columns in **Fig 10**), the combinations BMS-536924 + AKTIV, BMS-536924 + tryrphostin, and temsirolimus + AKTIV, reversed 50%, 52% and 48% of the IBC-GES genes detected, respectively (orange columns **Fig 10**).

**Fig 10.**
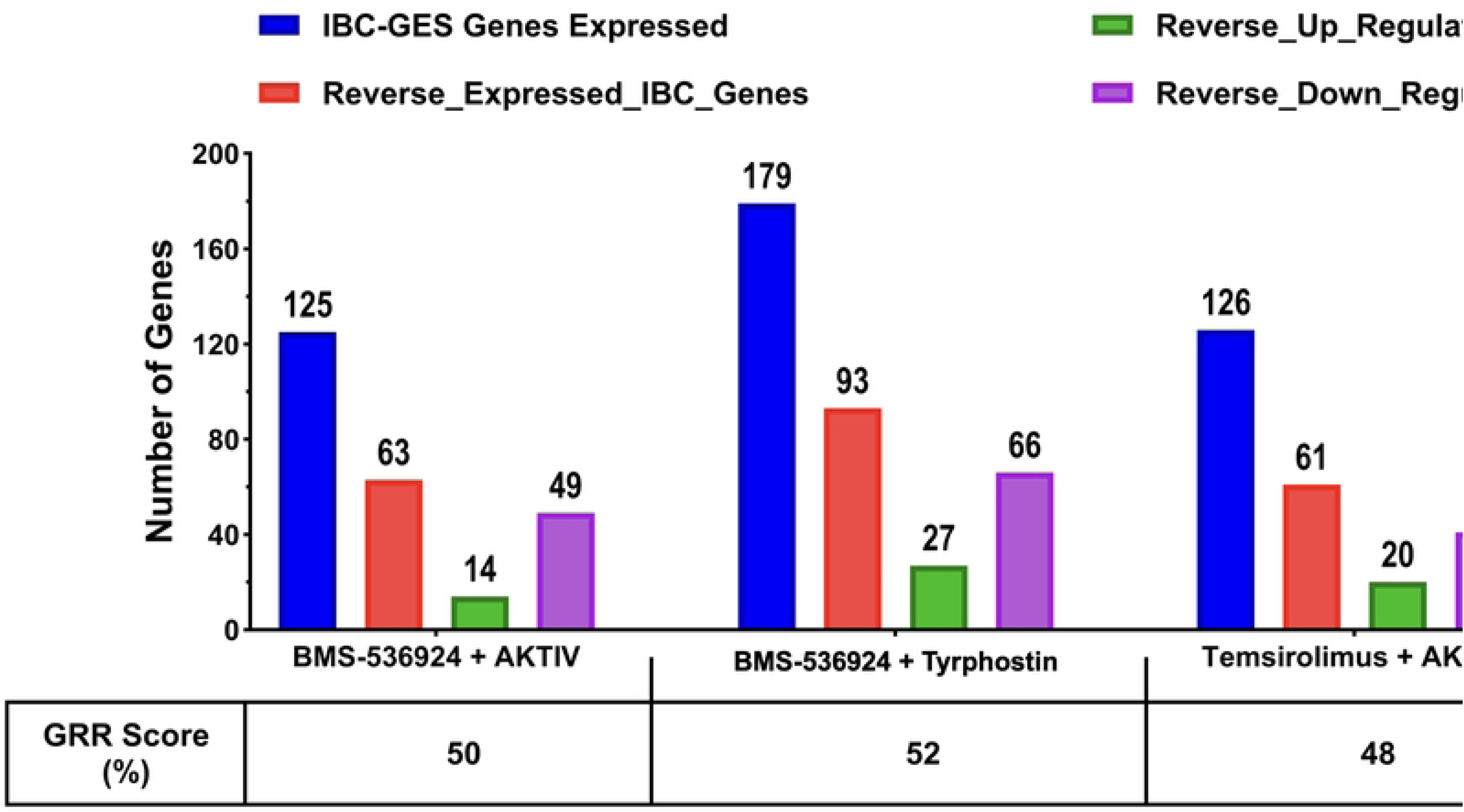
IBC gene expression signature changes induced by GRR combinations in SUM149 cells. SUM149 cells were incubated with GRR combinations or DMSO alone for 24 h, and RNA isolated for RNA-Seq analysis (n = 3 biological replicates). Left: DMSO (0.2%) vs (BMS-536924 (1 μM) + AKTIV (1 μM)); middle: DMSO (0.2%) vs (BMS-536924 (1 μM)) + tyrphostin AG1478 (0.1 μM)); right: DMSO (0.2%) vs (AKTIV (0.1 μM) + temsirolimus (10 μM)). Data was analyzed to identify genes significantly expressed in SUM149 cells (*p*adj < 0.05). Data are plotted as number of IBC-GES genes: detected in SUM149 cells (blue columns), number of detected IBC-GES genes reversed by the combination treatment (orange columns), and the number reversed up (green columns) and reversed down (purple columns).

## Discussion

Due to the limited availability of therapies targeting IBC, particularly for the triple-negative (TNBC) subtype, there is a critical need to identify novel targets and drugs to halt its rapid progression and aggressive metastasis [2, 58]. Drug repurposing is one approach to identify therapies for rare diseases such as IBC [22, 24, 59]. We have previously used two computational approaches: text mining (LWAS) and gene expression reversal rate analysis (GRR), as predictive tools for identifying novel drug candidates for IBC [27, 28].

To gauge the predictive value of our computational approaches for identifying new drugs to repurpose for IBC, we tested the drugs and compounds for anti-proliferative efficacy in four breast cancer cell lines, including SUM149, an IBC model. The oncology-focused LWAS method [27], showed a higher success rate, with 17 of 24 compounds (70%) demonstrating efficacy in the cell line models. For the GRR approach, which had identified both non-oncology and oncology drugs [28], the success rate was lower, with 6 out of 16 compounds (∼38%) showing efficacy in the cell line models. Among the LWAS-predicted drugs for IBC, paclitaxel, docetaxel, and doxorubicin showed the highest efficacy with nanomolar IC_50_ potencies in SUM149 (**Tables 1 and 5**). These drugs are FDA-approved for breast cancer and are also among those used as IBC treatment options [14]. All 11 of the LWAS drugs that have been tested in clinical trials for IBC (**S1 Table**), showed efficacy in our SUM149 Hoechst assay. Of the other 13 LWAS drugs/compounds not yet tested in clinical trials for IBC (**S1 Table**), 6 demonstrated activity in our SUM149 Hoechst assay (46% success), while 7 showed no activity. For the 7 with no activity, there were no reports on these drugs having activity in IBC models. Four novel drug candidates with in vitro efficacy were identified, which, to our knowledge, had not previously been investigated for IBC (as determined from a review of PubMed and clinicaltrials.gov (**S1 Table**)). These included topotecan, a topoisomerase I inhibitor, and cytarabine, an antimetabolite, which had potency across all tested cell lines, including IC_50_ values of 0.008 µM and 0.025 µM in SUM149, respectively. In our initial computational LWAS analysis [27], topotecan appeared among the drugs linked to medulloblastoma, while cytarabine was associated with acute lymphocytic leukemias, both diseases that clustered closely to IBC [27]. This semantic co-occurrence analysis suggests that other drugs targeting these “IBC-clustered” diseases, such as daunorubicin and vincristine, would also have activity in IBC models, and we did observe efficacy for these two drugs *in vitro* (**Table 5**).

**Table 5.**
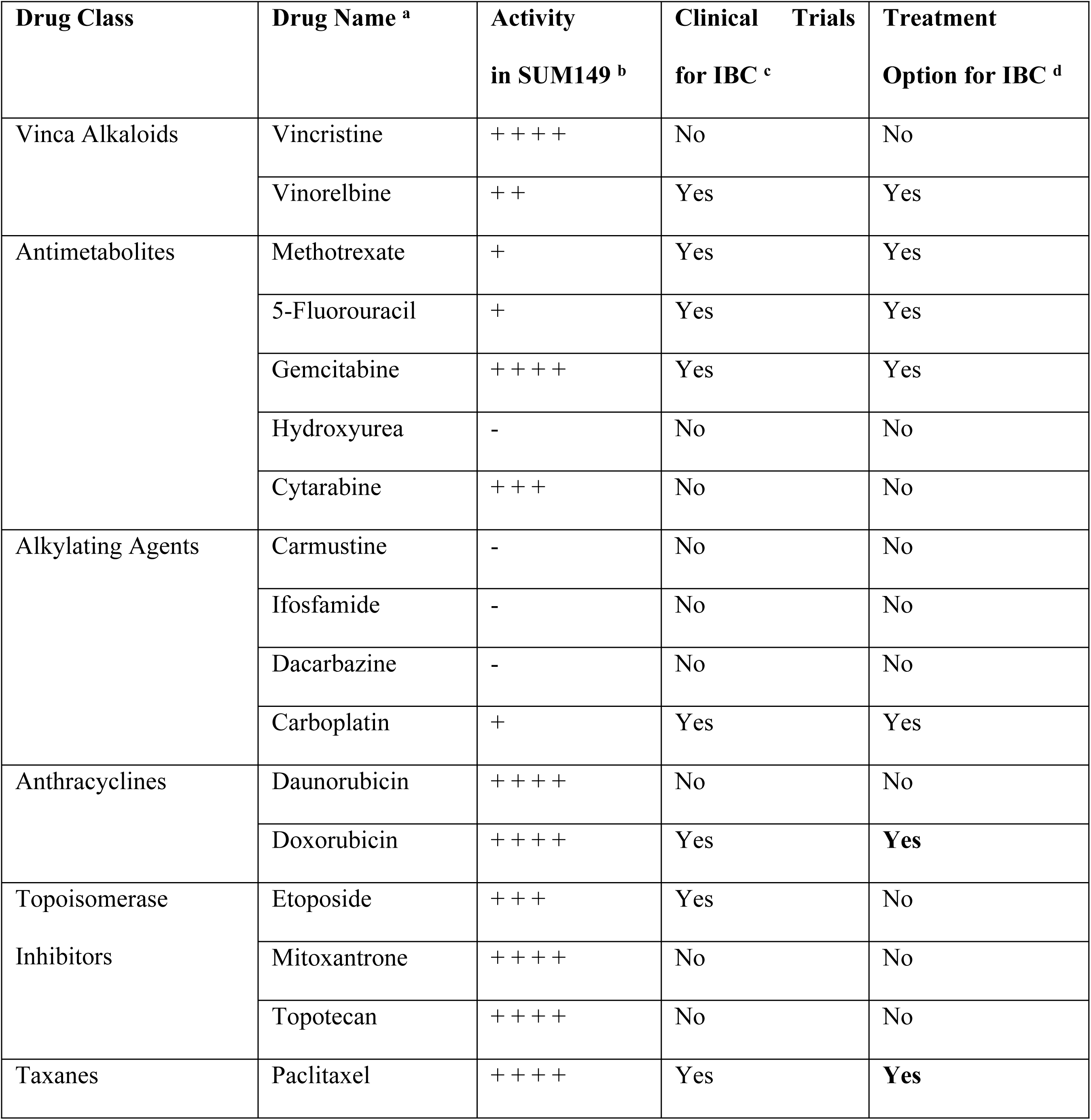
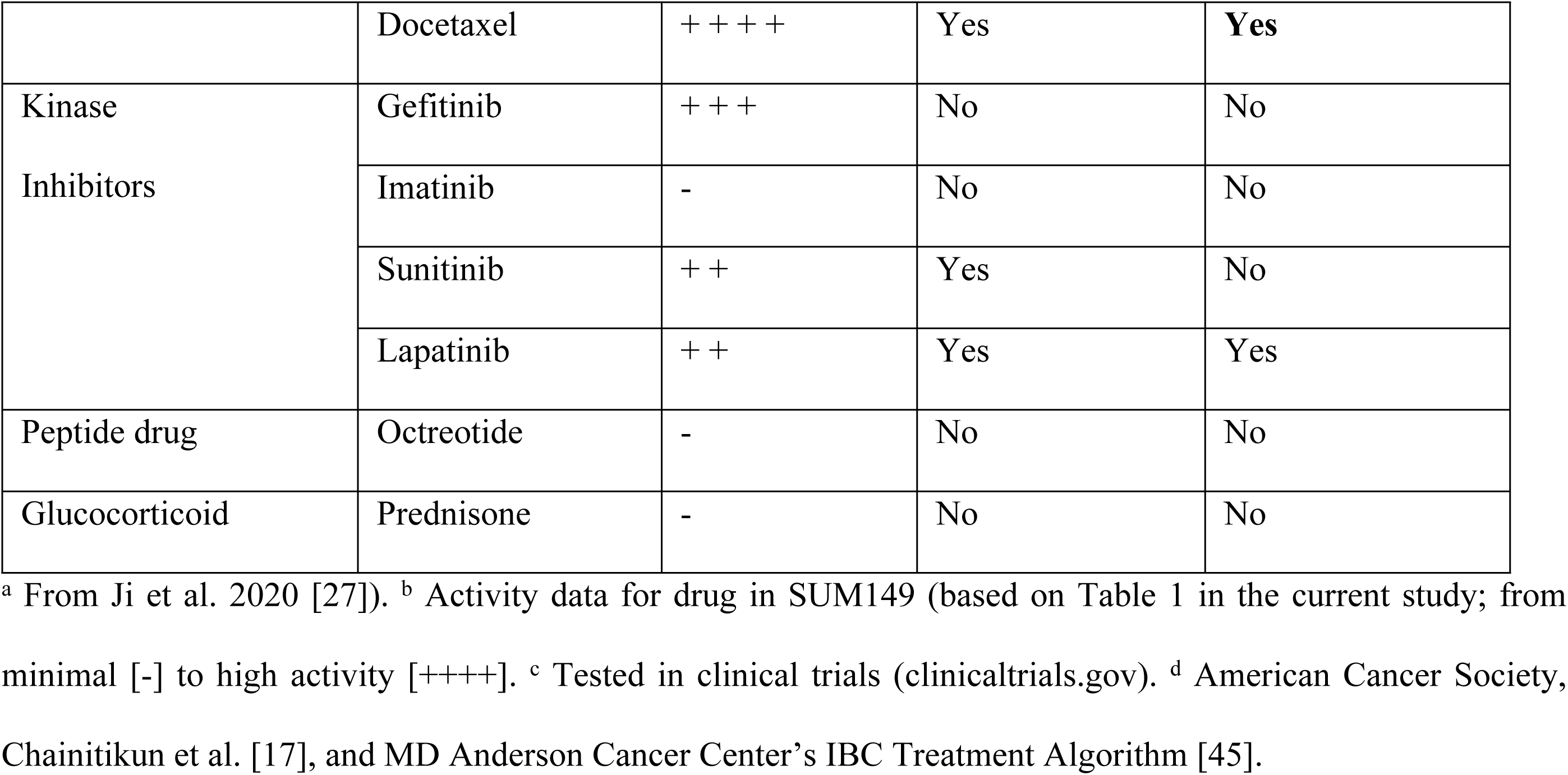
Clinical status of LWAS-predicted drug candidates in IBC.

TKIs, such as lapatinib, gefitinib, and sunitinib, showed varying potency and efficacy against SUM149, with gefitinib having the lowest IC_50_ value of 0.4 µM, and lapatinib achieving the highest efficacy ∼ 90%, at 10 µM. These two drugs target EGFR, which is highly expressed in the SUM149 cell line [60, 61], and, along with the finding that EGFR signaling plays a role in IBC progression and metastasis and can be blocked by EGFR inhibition [62], likely explains the effectiveness of these drugs in SUM149. While lapatinib has been tested clinically for IBC and showed limited success [63, 64], adverse events and toxicity were observed [17, 65], and there are currently no ongoing clinical trials for lapatinib.

Our validation of the GRR list identified novel effective drugs previously untested against IBC, primarily TKIs targeting multiple downstream pathways. Notably, tipifarnib, a farnesyltransferase inhibitor (FTI), demonstrated the highest potency (IC_50_ = 0.08 μM) in SUM149. This finding aligns with the contributing role of RhoC GTPase, a Ras superfamily member, in IBC pathogenesis [66, 67]. Since RhoC activation requires farnesylation, a post-translational modification, it can be targeted by FTIs. Tipifarnib has also progressed to Phase II clinical trials in locally advanced breast cancer (LABC), both as monotherapy and in combination with hormonal therapy [66, 67]. These results validate our approach and support the potential of other GRR effective compounds as repurposed therapeutic candidates for IBC treatment.

The IGF1-R inhibitors BMS-754807 and BMS-536924 showed IC_50_ values of 0.5 μM and 1.9 μM in the Hoechst assay, respectively. Differences in potency may be attributed to BMS-536924’s selective binding characteristics compared to BMS-754807’s broader IGF1-R/IR inhibition profile [68, 69]. BMS-536924 has previously been shown to have anti-growth activity in SUM149 and MCF7 cells [70]. While these IGF1-R inhibitors have been tested in preclinical breast cancer models [69, 71] and in clinical trials for breast cancer [72], they remain unexplored in clinical trials for IBC, suggesting potential novel therapeutic opportunities, although there are no FDA-approved small-molecule inhibitors of IGF1-R. Temsirolimus, an mTORC1 inhibitor [57], showed moderate potency for cytotoxicity in SUM149 (absolute IC_50_ = 4.3 μM), but a high potency for proliferation (relative IC_50_ = 0.1 nM). While FDA-approved for renal cell carcinoma, and assessed in clinical trials for metastatic breast cancer [73], temsirolimus has not been specifically investigated for IBC treatment, presenting an additional therapeutic avenue worth exploring both as monotherapy or in combinations. Another mTOR inhibitor, everolimus, which did not show up in our GRR analysis, has been FDA-approved for some types of metastatic breast cancer [74, 75]. Tyrphostin AG 1478, an EGFR inhibitor [54, 55], and AKTIV [56, 76] showed comparable potency with IC_50_ values of 0.23 μM in SUM149 cells, yet neither has been evaluated for IBC treatment. These results suggest promising therapeutic potential for expanding treatment options in IBC, with the possibility of evaluating additional FDA-approved EGFR inhibitors and the approved Akt inhibitor, both as single agents and in combination. The mechanisms of action for the effective LWAS and GRR candidates, as well as corresponding pathways, are illustrated in **Fig 11**.

**Fig 11.**
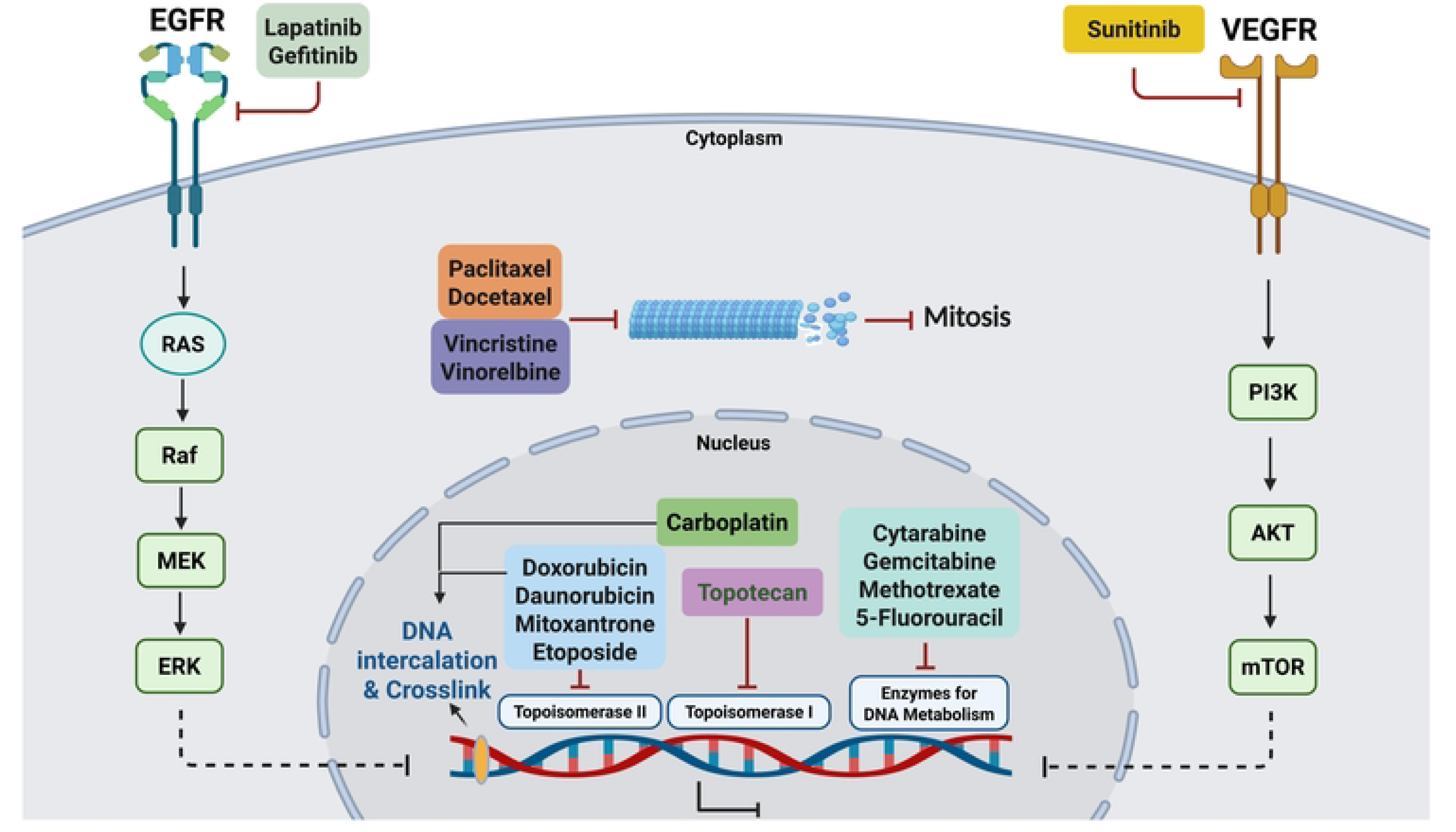
Cellular signaling pathways targeted by effective drugs previously identified through computational approaches. (A) LWAS 17 effective compounds, representing diverse drug classes. These include targeted therapies such as tyrosine kinase inhibitors (TKIs) and different chemotherapeutic agents categorized by their mechanisms: microtubule-targeting drugs, DNA-damaging agents (e.g., topoisomerase I and II inhibitors), platinum-based drugs, and antimetabolites. (B) GRR 6 effective compounds, all of which are TKIs, targeting various receptors and their downstream signaling pathways such as RAS/RAF/MEK/ERK and PI3K/AKT/mTOR. These pathways regulate critical cellular processes, including proliferation, differentiation, cell cycle, invasion, and epithelial-mesenchymal transition (EMT).

Due to challenges with single-drug approaches in cancer therapy, including drug resistance, we also examined drug combinations targeting multiple pathways. We focused on the most promising synergistic drug combinations identified in our initial screening. Further, due to the reported limitations of 2D cultures in identifying clinically relevant drug treatments, we also tested the synergistic combinations in 3D SUM149 tumor models [50], aiming to mimic the tumor architecture more accurately. We used a high-throughput approach to first screen combinations at single dose and then in a more extensive 8×8 matrix (**Fig 3**). A comparable approach to identify novel drug combinations with synergy for other cancers has been reported [77]. Our initial combinations were carried out by evaluating each drug at its EC_25_ concentration, with the Bliss method used to calculate Bliss excess scores. This resulted in identifying 25 combinations from the LWAS list and 9 combinations from the GRR list (*p* < 0.05). Then, we performed an 8×8 matrix analysis focusing on combinations with the highest Bliss excess scores. Our synergy analysis using SynergyFinder3 revealed three synergistic combinations meeting our stringent criteria, with scores greater than 5 across all reference models (Bliss, Loewe, HSA, and ZIP) at the MSA level. SynergyFinder+ was used to confirm these results at the micro visualization scale, with detailed bar plots presented in **S6 Fig**.

Interestingly, all three combinations originate from the GRR list that involves novel targeted compounds for IBC treatment. Four targeted agents were prominent within these identified combinations: BMS-536924 [53], AKTIV [56, 76], tyrphostin AG-1478 [54, 55], and temsirolimus [57]. Each of these drugs targets specific nodes within the PI3K/Akt/mTOR signaling cascade, supporting their potential in addressing the aggressive characteristics of IBC.

Our synergyfinder+ analysis on a micro visualization scale identified three synergistic combinations: termed our selected GRR combinations. The combinations include EGFR inhibitor, IGF1-R/IR inhibitor, Akt inhibitor, and mTOR inhibitor that can directly or indirectly target different components of the PI3K/Akt/mTOR pathway (**Fig 12**) a crucial signaling cascade in several cancers [78–80].

**Fig 12.**
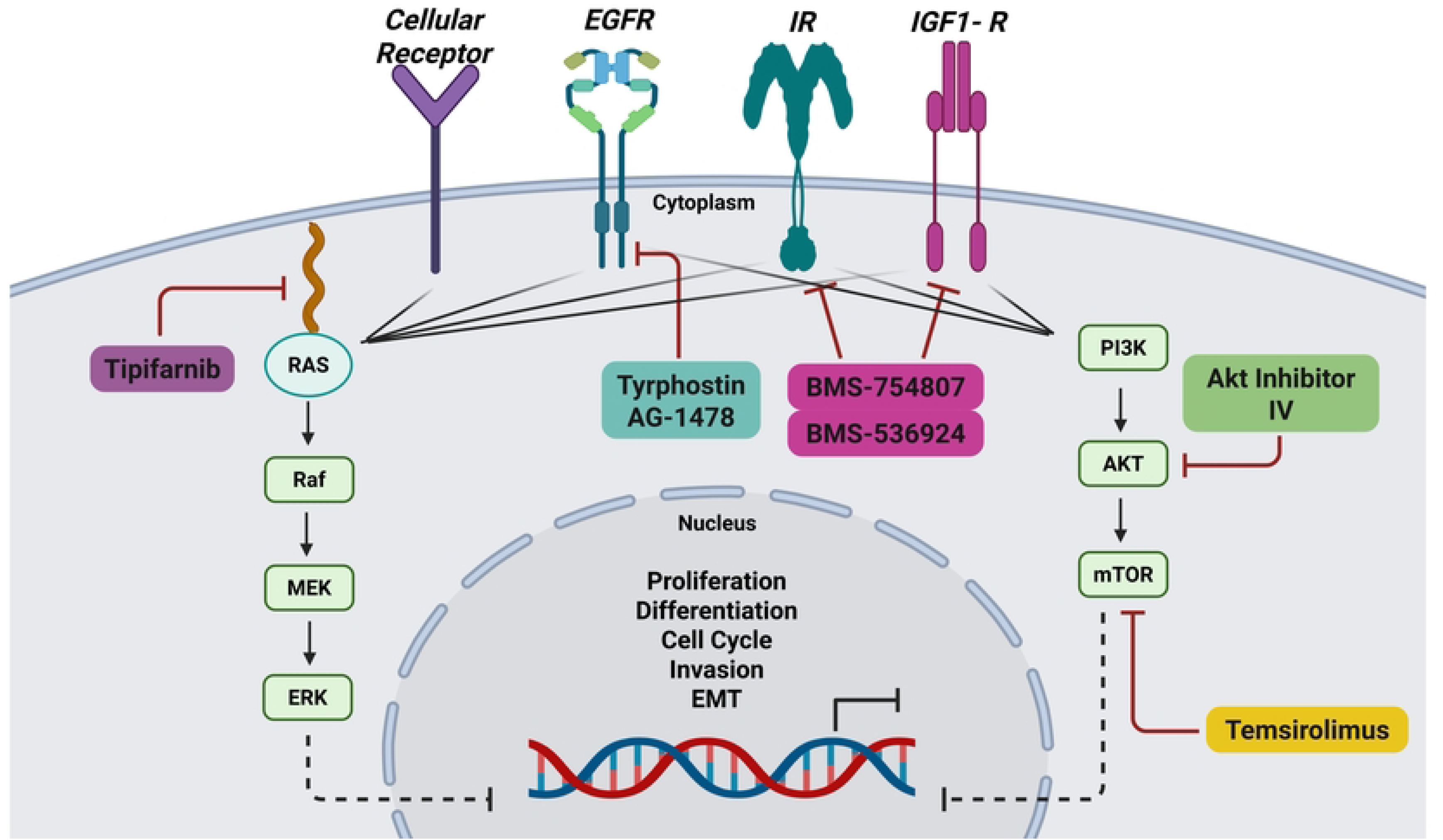
Schematic representation of PI3K/AKT/mTOR downstream signaling pathway targeted by the selected GRR combinations. The diagram shows the crosstalk between EGFR and IGF1-R with the PI3K/AKT/mTOR pathway to regulate the cell cycle. The three GRR drug combinations are highlighted: tyrphostin AG-1478 + BMS-536924, AKT inhibitor IV + BMS-536924, and AKT inhibitor IV + temsirolimus.

The first combination of BMS-536924 (1 µM) and tyrphostin AG-1478 (0.1µM) stands as a novel and promising approach, yielding the highest synergy score in both the primary combination screening with Bliss excess score = −0.266 as well as the secondary combination matrix screening in 2D and 3D models (Synergy scores summarized in **Table 4**). This combination primarily targets the crosstalk between EGFR and IGF1-R [81]. High levels of IGF-1R can cause resistance across different cancers [72, 82], and cancer cells can become resistant to IGF-1R inhibitors, by activating PI3K/mTOR pathways through alternative EGFR signaling [83, 84]. It was also reported that the interaction between IGF-1R and different receptor tyrosine kinases, such as EGFR, can create complex compensatory mechanisms that can limit treatment efficacy and promote resistance when targeting individual pathways [85]. These interactions occur through multiple mechanisms, including direct receptor associations and downstream signaling networks [86]. These compensatory mechanisms provide a strong rationale for dual targeting of both EGFR and IGF-1R with a consideration of reducing the cytotoxicity of the combined drugs by using the lowest concentrations that can achieve high synergy scores.

The IGF1-R/Akt signaling crosstalk represents a promising therapeutic target, addressed by our second identified drug combination of BMS-536924 (1 µM) and AKTIV (1 µM). Akt, also known as protein kinase B, mediates signal transduction from receptor tyrosine kinases, including IGF1-R and IR, via the PI3K pathway. This signaling cascade promotes G1/S cell cycle transition, which is crucial for cellular proliferation [78, 87]. The significant overlap between IGF1-R and Akt signaling provides a mechanistic rationale for the simultaneous targeting of both molecules.

Our studies identified another combination, temsirolimus (10 µM) and AKTIV (0.1 µM), that can simultaneously target mTOR and Akt. mTOR, a downstream effector, functions as a central regulatory kinase that operates multiple essential cellular survival processes through two different protein complexes: mTORC1 and mTORC2. Through these complexes, mTOR can execute its regulatory functions by phosphorylating key downstream proteins, including Akt [88, 89]. We suggest that targeting both proteins through our identified combination of temsirolimus, an mTORC1 inhibitor [57, 90], alongside AKTIV may effectively reduce the likelihood of drug resistance development while achieving more complete pathway suppression. Capivasertib was recently approved by FDA as the first Akt inhibitor drug for treating hormone receptor (HR)-positive, human epidermal growth factor 2 (HER2)-negative advanced or metastatic breast cancer [91, 92]. This approval validates Akt as a promising clinical target and supports our findings that Akt inhibitors, which appeared in two of the three identified combinations, could be potentially effective in treating IBC.

We also tested all the selected combinations in 3D cell models. Although all these combinations showed synergistic effects at the proliferation level in 2D, the temsirolimus and AKTIV combination showed a significant shift in 3D cell models, where temsirolimus loses its single-agent efficacy. This phenomenon may represent a mechanistic transition from classical synergy observed in 2D cultures (where both agents demonstrate independent activity) to a potentiator or enhancer effect in 3D models, as described previously [93]. While temsirolimus alone appeared to be ineffective, it significantly sensitized cells to AKTIV, enhancing the growth inhibition from 16.4% as an AKTIV single agent to 61.2% in combination. Mechanistically, this transition reflects fundamental aspects of 3D tumor biology, where cells in spheroids can activate survival pathways (e.g., mTOR), which has been reported to have a greater effect on survival in 3D than in 2D breast cancer cell cultures and may contribute to drug resistance [94]. This shift allows for more accurate modeling of tumor behavior and drug responses, making 3D cultures a valuable tool in cancer research and drug development. This could potentially be translated to enhance therapeutic efficacy in clinical applications, particularly in IBC, where these pathways are known to drive tumor growth and survival [2, 16].

To investigate how these drug combinations might affect IBC cell migration, we performed wound-healing assays to assess drug effects on cell motility. Our results demonstrated that the combinations of (BMS-536924 + tyrphostin AG-1478) and (temsirolimus + AKTIV) enhanced the inhibitory effects of the individual drugs on cell motility and migration. This aligns with studies reporting the role of the PI3K/Akt/mTOR pathway in cell motility/invasion, and its crosstalk with IGF1-R and EGFR signaling, either in SUM149 [95, 96] or in TNBC [97–100]. In our study, the combination co-targeting Akt upstream (by AKTIV) and mTORC1 downstream (by temsirolimus) may effectively shut down the PI3K/Akt/mTOR axis and prevent any compensatory rebound activation. In essence, Akt inhibition blocks the bulk of signaling through the pathway, while mTORC1 inhibition ensures that any Akt activity not fully suppressed (or any Akt-independent mTOR activation) is also blocked. These results demonstrate that effective suppression of cell migration in SUM149 requires a multi-targeted approach, given the interconnected nature of growth factor receptor signaling, and involves the PI3K/Akt/mTOR pathway.

We also assessed the GRR combinations for direct effects on signaling pathways in SUM149 cells, and we observed that the combination of AKTIV and temsirolimus suppressed phosphorylation of both Akt and mTOR (**Fig. 9A**). While temsirolimus appeared to be the primary driver of this suppression, the combination showed enhanced suppressive effects compared to control, indicating potential synergistic interactions within the Akt/mTOR signaling pathway. The combination of AKTIV and BMS-536924 also suppressed Akt phosphorylation, predominantly by BMS-536924 (**Fig. 9B**). While IGF1-R phosphorylation trended lower with AKTIV and BMS-536924 treatment, it did not reach significance due to some variability across experiments. This was also complicated by our observation that BMS-536924 appeared to increase total IGF1-R expression, perhaps indicating the activation of compensatory mechanisms in response to pathway inhibition, as reported previously [101, 102]. While in cell-based efficacy assays, AKTIV demonstrated anti-proliferative effects with an IC_50_ of 0.2 µM, both 1 µM and 0.1 µM, AKTIV did not appear to reduce p-Akt expression, perhaps due to differences in assay timing (24 h for Western blot compared to 72 h for the cell assays), or indicating an off-target effect in the cell assays. In the BMS-536924 + tyrphostin combination (**Fig. 9C**), although not significant, pEGFR and pIGFR levels trended lower compared to controls.

The gene reversal approach involves identifying drugs capable of reversing disease-associated gene expression signatures back to a ‘normal/healthy’ transcriptomic state [31]. In our previous computational GRR study, we had first generated a 387 IBC-gene expression signature (IBC-GES) compiled from published studies using IBC patient samples (see citations in [28]). 297 of these 387 genes had drug data in the LINCS database, and our GRR analysis [28] identified drug perturbations that significantly reversed the expression of genes in the IBC-GES list. For example, tyrphostin AG-1478 had a GRR score of 59% [28], meaning 175 of the 297 genes were reversed. In our RNA-Seq study herein assessing the three drug combinations in SUM149 cells, ∼half of the detectable IBC-GES genes were reversed in expression by the drug combinations, suggesting some reprogramming of IBC-associated transcriptional profiles.

An assessment of the global gene expression changes in our RNA-Seq data identified significant downregulation of cell cycle regulators, DNA replication enzymes, and survival factors across all treatment groups. For example, both BMS-536924 + tyrphostin and the BMS-536924 + AKTIV treatments reduced expression of RRM2, a key driver of DNA synthesis and proliferation. RRM2, which encodes ribonucleotide reductase, is overexpressed in aggressive basal-like and TNBC, where elevated levels correlate with adverse clinical features including larger tumor burden, lymph node involvement, and disease recurrence [103, 104]. Another gene, cyclin E2 (CCNE2) was found to be downregulated by two combinations: AKTIV + temsirolimus and BMS-536924 + AKTIV. CCNE2, an S phase cyclin is frequently overexpressed in breast cancer [105] with previous studies reported the overexpression of Cyclin E including CCNE2 in IBC, with correlation of poor prognosis in IBC patients [106].

All these results highlight the therapeutic promise of the identified combinations in combating IBC, and to our knowledge, we are the first to report these novel combinations. The three most effective combinations all came from drugs identified in the GRR computational analysis emphasizing the need for multiple approaches to identify effective novel combinations. While these initial findings are promising, significant studies are still needed to examine their long-term efficacy, potential resistance mechanisms, and validation in additional IBC models and patient-derived samples before advancing to clinical trials.

## Conclusions

This study bridges the gap between computational predictions and experimental validation in drug repurposing for IBC. Our findings validate the potential of our *in-silico* approaches and identified novel, potentially synergistic drug combinations that require further exploration for IBC treatment. By combining computational methods with experimental validation, we have opened new avenues for IBC therapy that could enhance therapeutic responses for individuals affected by this aggressive breast cancer subtype. Future research directions will be needed to validate the identified drug combinations in additional IBC cell models beyond SUM149 as well as *in vivo* studies to assess the different combination’s therapeutic efficacy, establish optimal dosing regimens, and characterize toxicity profiles in a physiologically relevant context. This systematic approach aimed to uncover novel IBC-specific therapeutic targets that could lead to new effective treatment strategies.

## Author Contributions

Conceptualization, K.P.W., X.J. and W.Z.; methodology, E.A.S., M.T., and X.J..; software, E.A.S. and J.E.S.; formal analysis, E.A.S., J.E.S. and K.P.W.; investigation, EA.S., M.T. and X.J.; writing—original draft preparation, K.P.W., J.E.S. and E.A.S.; writing—review and editing, E.A.S., X.J., M.T., M.S.D., W.Z., J.E.S. and K.P.W.; visualization, E.A.S.; supervision, K.P.W. and J.E.S.; funding acquisition, K.P.W. All authors have read and agreed to the published version of the manuscript.

## Data Availability Statement

The raw data supporting the conclusions of this article will be made available by the authors on request.

## Acknowledgements

Schematic and Pathway Figures, and graphical abstract were created with BioRender.com.

## Supporting information

### Supplemental Figure Legends

**S1 Fig. Representative dose response curves for the LWAS drug candidates with efficacy in breast cancer cell lines using the Hoechst assay.** (A) SUM159, (B) MDA-MB-231, and (C) MCF-7. Each graph shows the relative cell count (%) plotted against drug concentration (μM) on a logarithmic scale. Plots are grouped by drug mechanisms: antimetabolites, microtubule inhibitors, topoisomerase inhibitors, and tyrosine kinase Inhibitors. Error bars indicate ± SD from replicate experiments.

**S2 Fig. Representative dose response curves of the GRR effective compounds with efficacy in different breast cancer cell lines using the Hoechst assay.** (A) SUM159, (B) MDA-MB-231 and (C) MCF-7s. The x axis represents the concentration of the compound in µM while the y axis represents the cell count % normalized to DMSO control. All the experiments were done in technical and independent experiments (n=3). Error bars indicate ± SD from replicate experiments.

**S3 Fig. Drug sensitivity heatmap profiles and dose response curves for LWAS drugs in IBC and non-IBC breast cancer cell line panel by MTT assay.** (A) Drug response heatmaps across all the breast cancer cell lines. The x-axis represents the concentration of each drug delivered in a dose-response, while the y-axis lists the compounds. The color gradient indicates the level of inhibition, with red indicating higher inhibition and green indicating lower inhibition. Representative dose response curves grouped by drug class: antimetabolites, microtubule inhibitors, topoisomerase inhibitors, and tyrosine kinase inhibitors (TKIs), for (B) SUM149, (C) SUM159, (D) MDA-MB-231 and (E) MCF-7. Inhibition data plotted as the percentage (%) of cell proliferation relative to control versus drug concentration (µM). Error bars indicate ± SD from replicate experiments.

**S4 Fig**. **Drug sensitivity heatmap profiles and dose response curves for GRR drugs in IBC and non-IBC breast cancer cell line panel by MTT assay.** (A) Drug response heatmaps across all the breast cancer cell lines. The x-axis represents the concentration of each drug delivered in a dose-response, while the y-axis lists the compounds. The color gradient indicates the level of inhibition, with red indicating higher inhibition and green indicating lower inhibition. Representative dose response curves for each drug: BMS-536924, BMS-754807, Akt Inhibitor IV, temsirolimus, tipifranib, and tyrphostin AG 1478 in the four breast cancer cell lines with (B) SUM149, (C) SUM159, (D) MDA-MB-231 and (E) MCF-7. Inhibition data plotted as the percentage (%) of cell proliferation relative to control versus drug concentration (µM). Error bars indicate ± SD from replicate experiments.

**S5 Fig**. **Contour Plots of selected LWAS drug combinations with their synergy scores across Bliss, ZIP, Loewe and HSA reference models obtained from SynergyFinder3.** Synergy scores across Bliss, ZIP, Loewe and HSA reference models were obtained from SynergyFinder3. (A) Lapatinib + doxorubicin (B) Lapatinib + carboplatin (C) Carboplatin + etoposide (D) Carboplatin + topotecan (E) Vincristine + docetaxel, (F) Gefitinib + carboplatin and (G) Carboplatin + cytarabine. In each plot, x and y axes represent drug concentrations (µM). Red areas indicate synergy, while green areas indicate antagonism. Black boxes highlight the MSA, with scores shown. Red circles in the top right corner of each plot indicate the overall δ score. Color scales on the right of each plot show the range of synergy scores. The data represented the average of three independent experiments (n = 3).

**S6 Fig. Quantitative synergy analysis of GRR drug combinations in SUM149 2D and 3D SUM149 cell models.** Synergy scores were computed using SynergyFinder+ across four established reference models (ZIP, HSA, Bliss, and Loewe). Results are shown as bar plots for each combination. (**A**) BMS-536924 and Tyrphostin AG-1478 combination, (**B**) BMS-536924 and Akt Inhibitor IV combination, and (**C**) Temsirolimus and Akt Inhibitor IV combination. The upper figure for each combination represents the results from the 2D experiments, while the lower figure represents the results from the 3D experiments. Bar plots represent the drug combination landscape, where red bars indicate positive synergy scores, and green bars indicate antagonistic effects. Blue bars highlight the optimal combinations achieving maximal synergy scores and growth inhibition while minimizing individual drug concentration, which were: (1 µM BMS-536924 + 0.1 µM tyrphostin AG-1478), (1 µM BMS-536924 + 1 µM Akt Inhibitor IV), and (10 µM temsirolimus + 0.1 µM Akt Inhibitor IV). The data shown represents the average of three independent experiments.

**S7 Fig. Western blot analysis of protein expression levels in response to the selected GRR drugs and combinations.** Representative whole-blot Western blots for replicate DMSO and combination incubations with SUM149 cells. (A) Temsirolimus (10 µM) and Akt inhibitor IV (0.1 µM) combination effects on the expression of p-Akt, total Akt, p-mTOR, and total mTOR; (B) Akt Inhibitor IV (1 µM) and BMS-536924 (1 µM) combination effects on the expression of p-Akt, total Akt, p-IGF1-R, and total IGF1-R; and (C) Tyrphostin AG-1478 (0.1 µM) and BMS-536924 (1 µM) combination effects on the expression of p-EGFR, total EGFR, p-IGF1-R, and total IGF1-R. β-actin served as a loading control. Right panels show densitometric band quantification of DMSO relative to each treatment. The Li-COR system was used for all proteins except for IGF1-R and pIGF1-R, for which we used HRP-tagged secondary antibodies and iBRIGHT imaging. Statistical significance set as *P < 0.05, **P < 0.01, ***P < 0.001.

**S8 Fig. Global gene expression changes induced by GRR combinations in SUM149 cells.** SUM149 cells were incubated with GRR combinations or DMSO alone for 24 h, and RNA isolated for RNA-Seq (n = 3 biological replicates). RNA-Seq data were analyzed to identify genes significantly differentially expressed in SUM149 cells (padj < 0.05). (A) Differential gene expression (DGE) analysis. The number of genes differentially expressed in the combination treatment relative to the DMSO control. Total (black columns), up-regulated (red columns) and down-regulated (blue columns). (B) Volcano plots display log2 fold change (x-axis) versus −log10 p value (y-axis); vertical red lines mark the fold-change cutoffs, and the horizontal line denotes the significance threshold (FDR-adjusted *p* < 0.05). Red points indicate upregulated transcripts, and blue points indicate downregulated transcripts; representative top hits are labeled. (C) Heat maps show normalized expression values (log scale; blue = lower, red = higher) for the top 25 up-regulated and top 25 down-regulated protein-coding genes across biological triplicates. (i) to (iii) Treatment comparison: (i) DMSO (0.2%) vs BMS-536924 (1 μM) + tyrphostin AG1478 (0.1 μM) (BMS24-Tyrpho). (ii) DMSO (0.2%) vs BMS-536924 (1 μM) + AKTIV (1 μM) (BMS24-Akt). (iii) DMSO (0.2%) vs AKTIV (0.1 μM) + temsirolimus (10 μM) (Tem-Akt).

Compared with DMSO, the BMS-536924 + tyrphostin AG1478 group showed the strongest transcriptional response, with 2868 DEGs, followed by AKTIV + temsirolimus with 2434 DEGs. Together, these results show that each combination affects the cells differently, with BMS-536924 + tyrphostin AG1478 having the greatest overall impact. Heatmap analysis revealed that this combination induced significant changes in genes such as GSTA2, OMD, and HIST1H2AJ. Volcano plots further highlight important gene shifts, including decreases in F3, UHRF1, and RRM2, and increases in PDK4, MMP7, and ELF3, confirming the substantial impact of the BMS-536924 + tyrphostin AG1478 combination.

## Supplemental Tables

**S1 Table. The 24 LWAS identified drugs for IBC (from Ji et al., 2020).** The table presents the corresponding drug class, PubMed entry, Clinical Trials entry, and NCT number.

**S2 Table. The 19 GRR identified drugs/compounds (from Ji et al., 2023)**. The table shows with the corresponding drug class for each drug and its mechanism of action.

**S3 Table. Efficacy of LWAS and GRR predicted drugs and compounds in four breast cancer cell lines using the MTT assay.** The table lists the IC_50_ values for each drug determined in the MTT assay across the cell lines, with values reported as the mean ± SD from 2 independent experiments.

**S4 Table. Selected drugs and compounds from LWAS and GRR active lists for combination studies.** The table shows their corresponding EC_25_ values and the highest concentration used for the 8 x 8 matrix.

