## Supplemental figures and tables for "From *in silico* prediction to experimental validation: Identification of drugs and novel synergistic combinations that inhibit growth of inflammatory breast cancer cells"

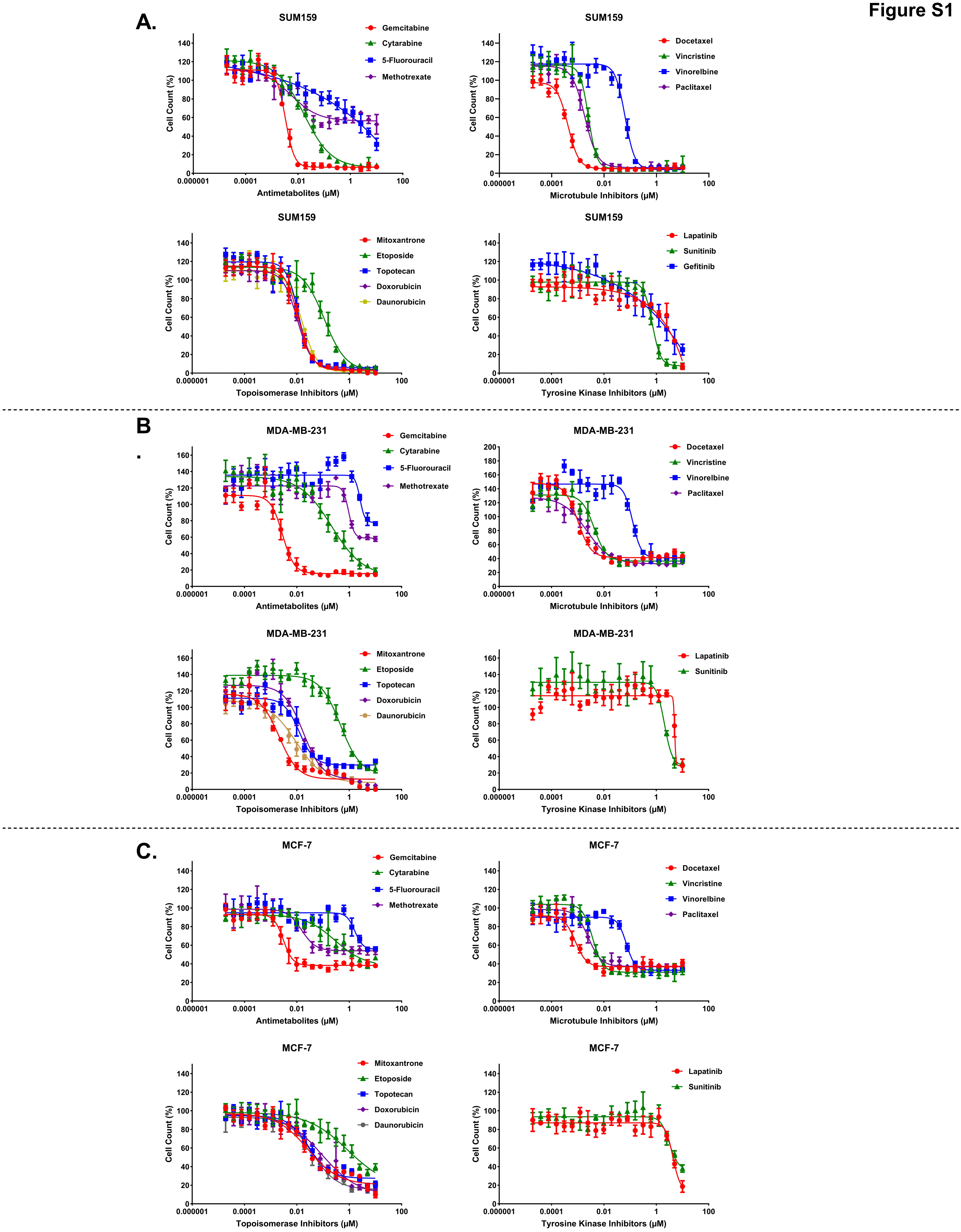
A.
SUM159
Gemcitabine
Cytarabine
5-Fluorouracil
Methotrexate
Cell Count (%)
Antimetabolites (µM)
SUM159
Docetaxel
Vincristine
Vinorelbine
Paclitaxel
Microtubule Inhibitors (µM)
SUM159
Mitoxantrone
Etoposide
Topotecan
Doxorubicin
Daunorubicin
Topoisomerase Inhibitors (µM)
SUM159
Lapatinib
Sunitinib
Gefitinib
Tyrosine Kinase Inhibitors (µM)
B
MDA-MB-231
Gemcitabine
Cytarabine
5-Fluorouracil
Methotrexate
Antimetabolites (µM)
MDA-MB-231
Docetaxel
Vincristine
Vinorelbine
Paclitaxel
Microtubule Inhibitors (µM)
MDA-MB-231
Mitoxantrone
Etoposide
Topotecan
Doxorubicin
Daunorubicin
Topoisomerase Inhibitors (µM)
MDA-MB-231
Lapatinib
Sunitinib
Tyrosine Kinase Inhibitors (µM)
C.
MCF-7
Gemcitabine
Cytarabine
5-Fluorouracil
Methotrexate
Antimetabolites (µM)
MCF-7
Docetaxel
Vincristine
Vinorelbine
Paclitaxel
Microtubule Inhibitors (µM)
MCF-7
Mitoxantrone
Etoposide
Topotecan
Doxorubicin
Daunorubicin
Topoisomerase Inhibitors (µM)
MCF-7
Lapatinib
Sunitinib
Tyrosine Kinase Inhibitors (µM)
0.000001
0.0001
0.01
1
100

**Figure S1**

Figure S2

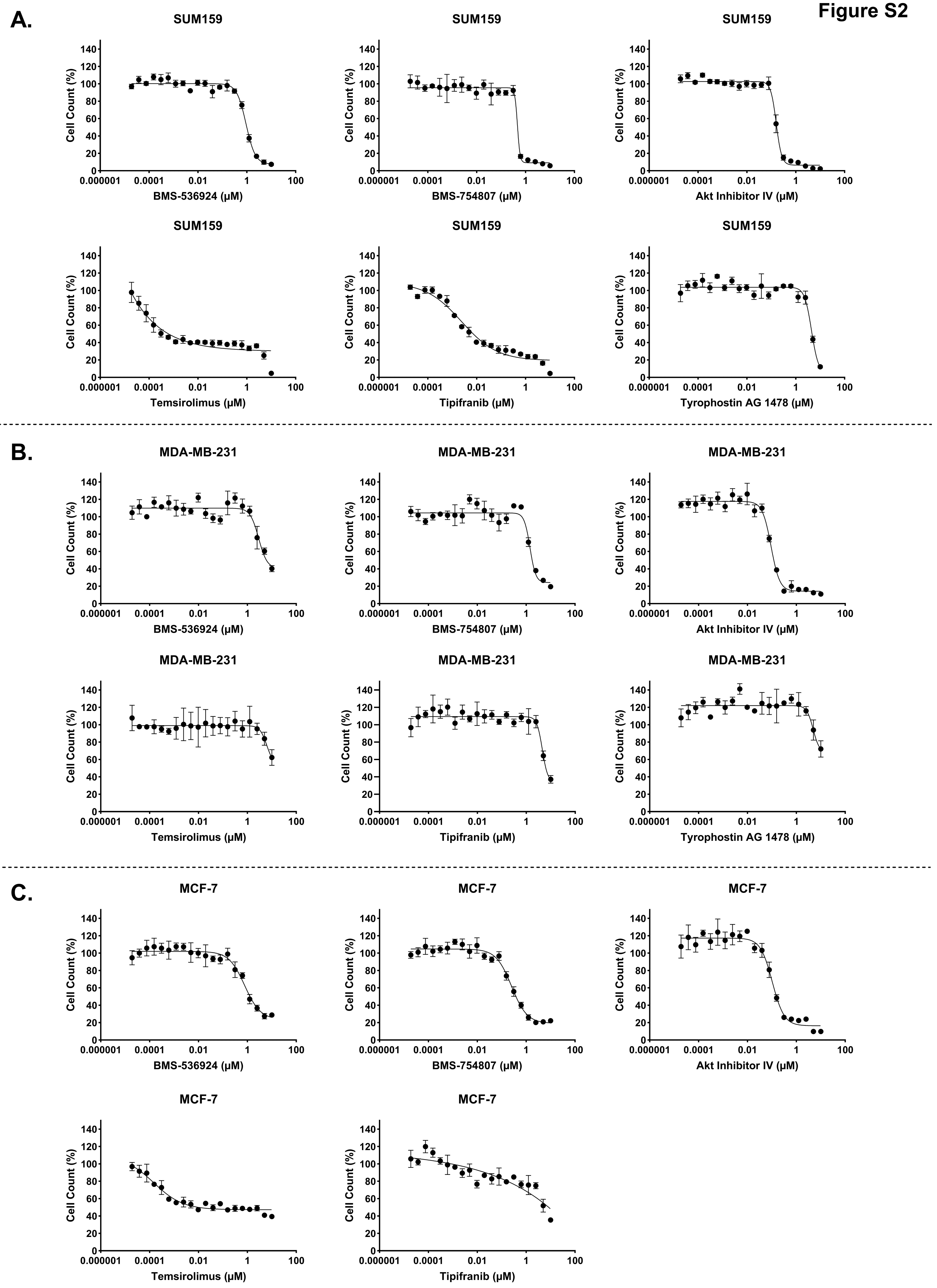
A.
SUM159
Cell Count (%)
BMS-536924 (μM)
SUM159
Cell Count (%)
BMS-754807 (μM)
SUM159
Cell Count (%)
Akt Inhibitor IV (μM)
SUM159
Cell Count (%)
Temsirolimus (μM)
SUM159
Cell Count (%)
Tipifranib (μM)
SUM159
Cell Count (%)
Tyrophostin AG 1478 (μM)
B.
MDA-MB-231
Cell Count (%)
BMS-536924 (μM)
MDA-MB-231
Cell Count (%)
BMS-754807 (μM)
MDA-MB-231
Cell Count (%)
Akt Inhibitor IV (μM)
MDA-MB-231
Cell Count (%)
Temsirolimus (μM)
MDA-MB-231
Cell Count (%)
Tipifranib (μM)
MDA-MB-231
Cell Count (%)
Tyrophostin AG 1478 (μM)
C.
MCF-7
Cell Count (%)
BMS-536924 (μM)
MCF-7
Cell Count (%)
BMS-754807 (μM)
MCF-7
Cell Count (%)
Akt Inhibitor IV (μM)
MCF-7
Cell Count (%)
Temsirolimus (μM)
MCF-7
Cell Count (%)
Tipifranib (μM)

**Figure S3A,B**

**A.**

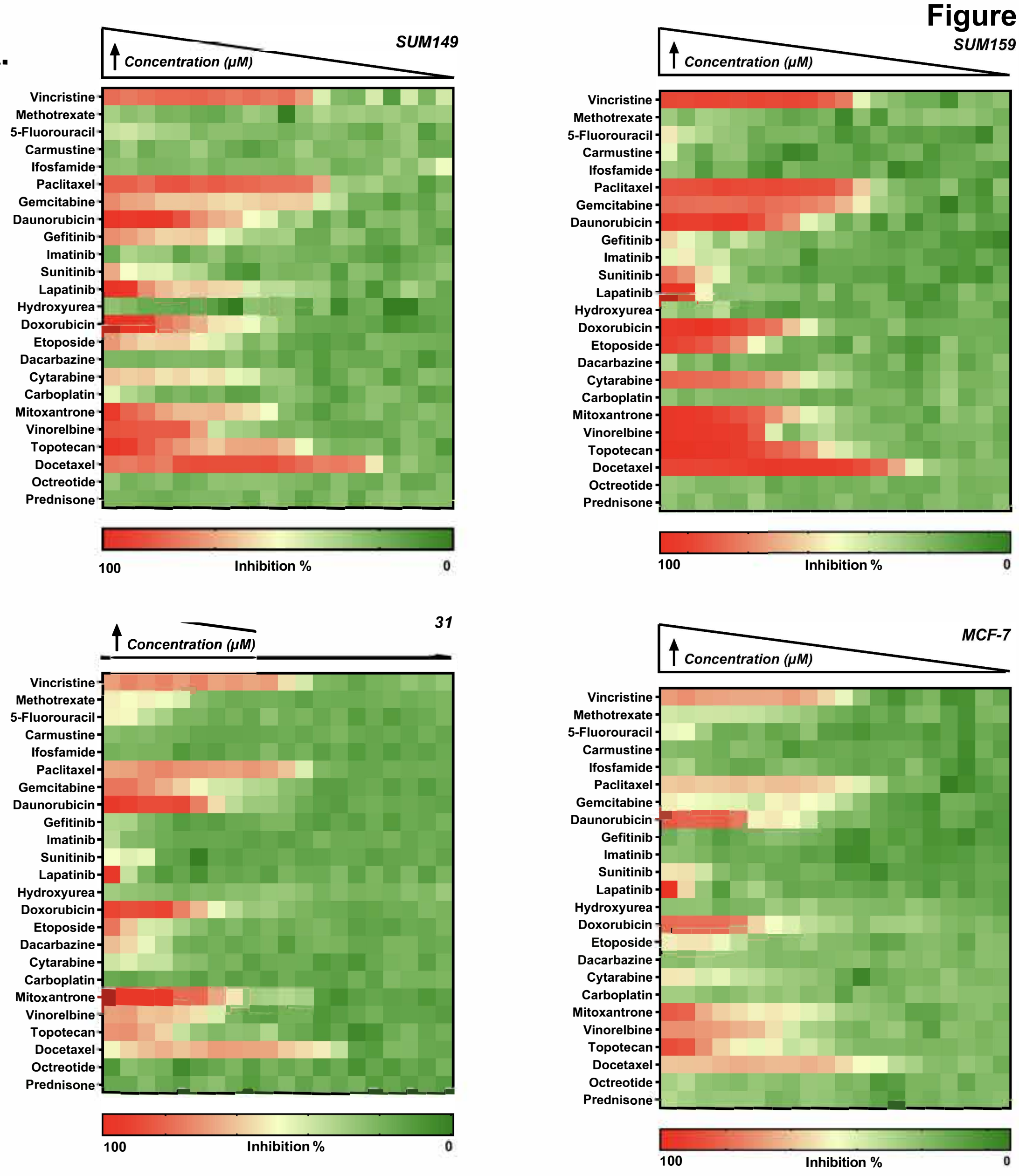


**B.**

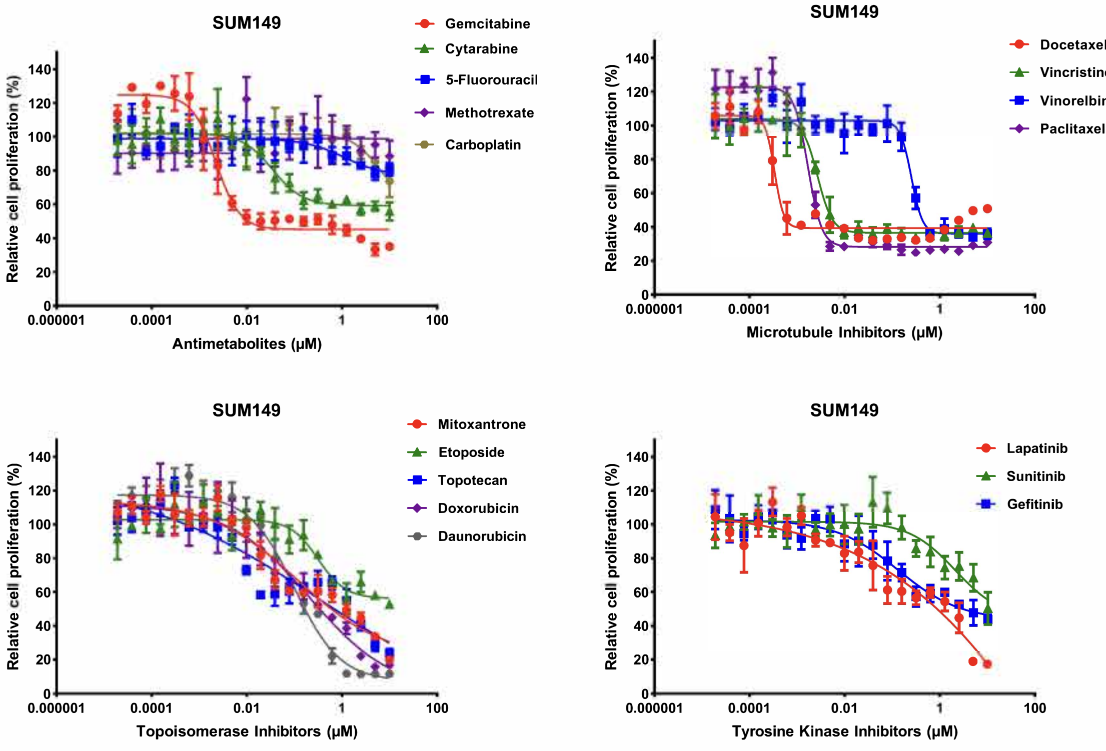

**Figure S3C-E**

C.

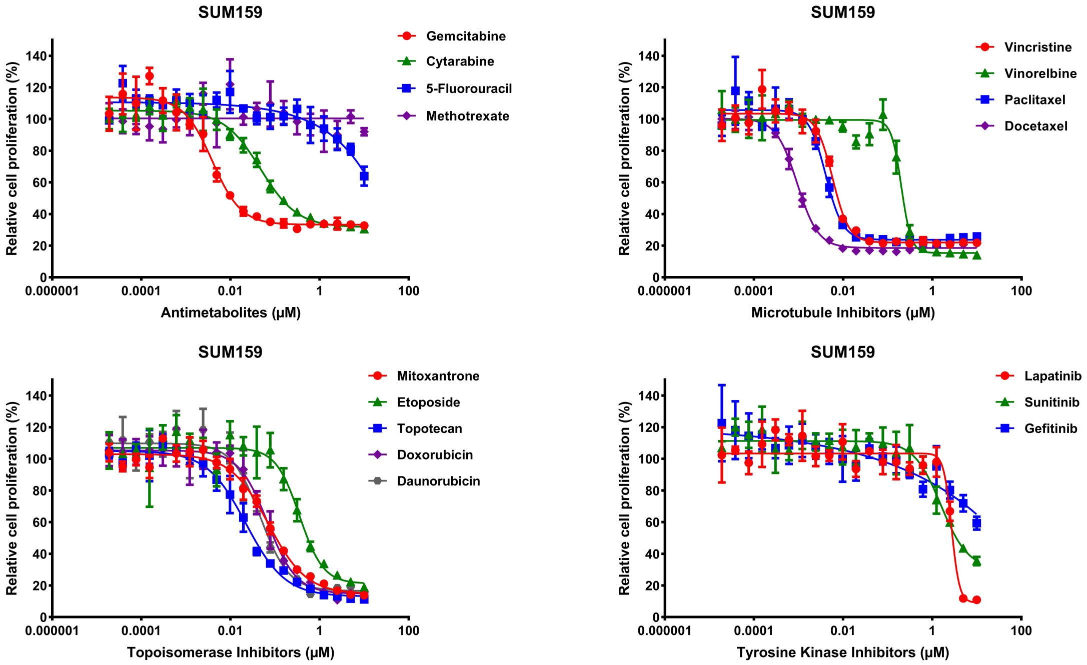


D.

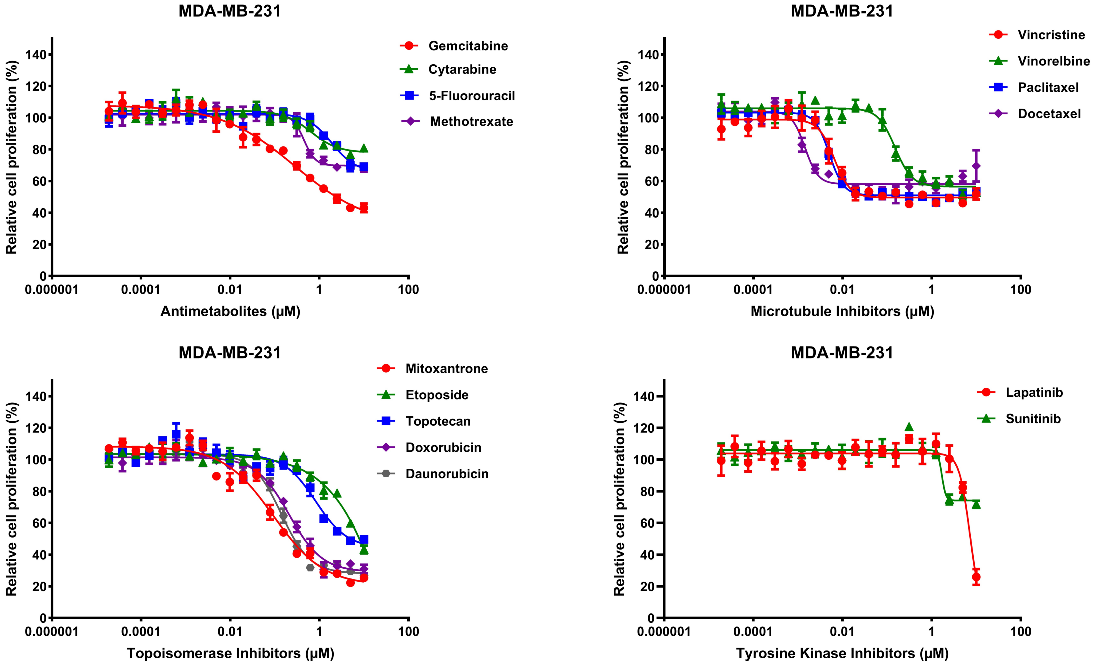


E.

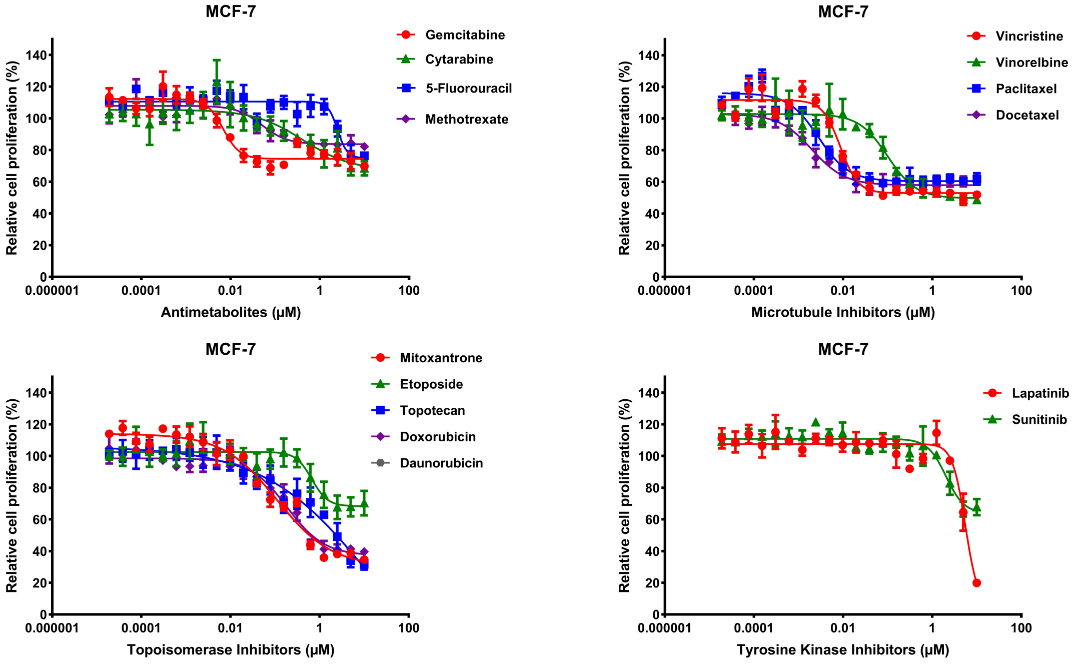

Figure S4A,B

A.

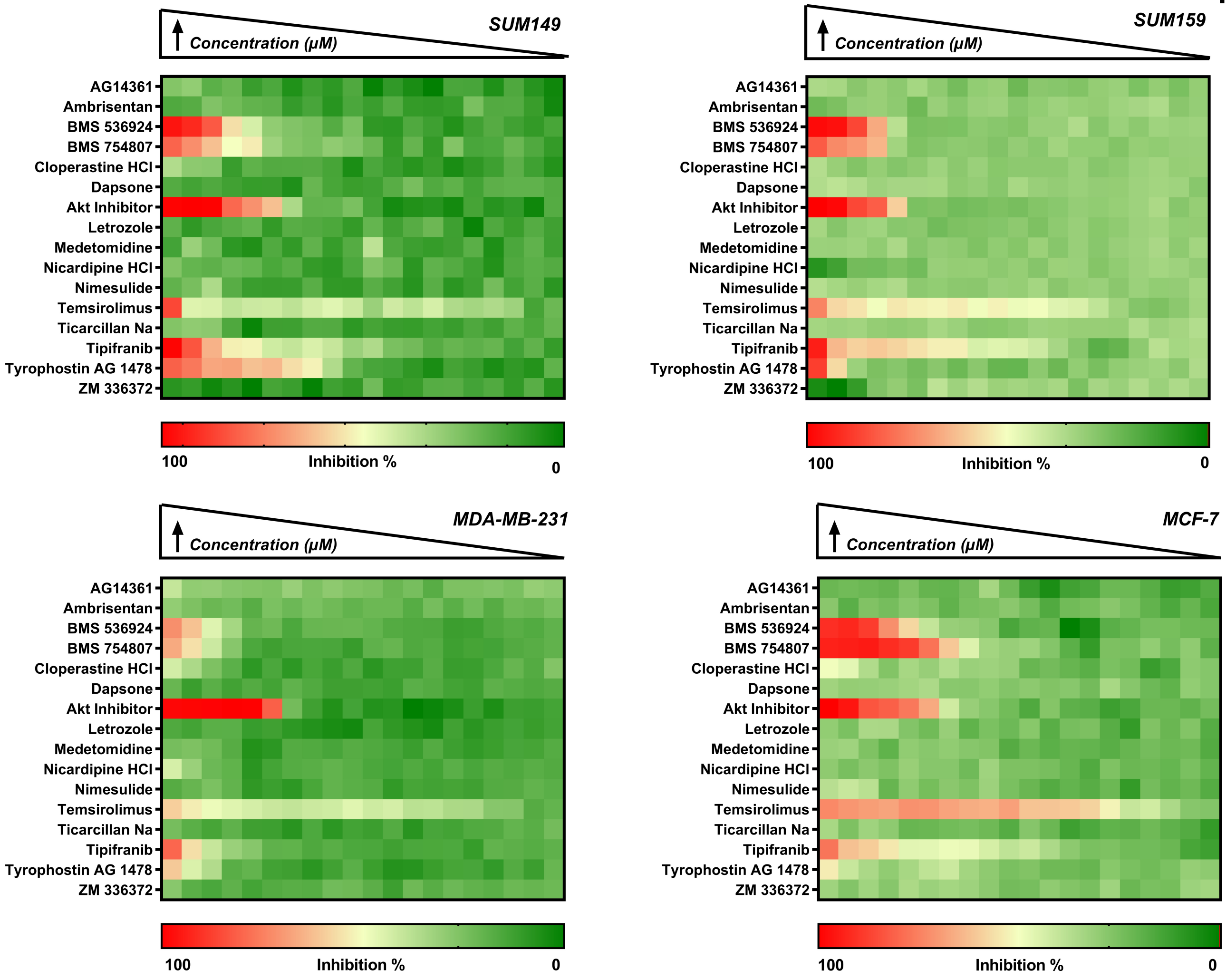


B.

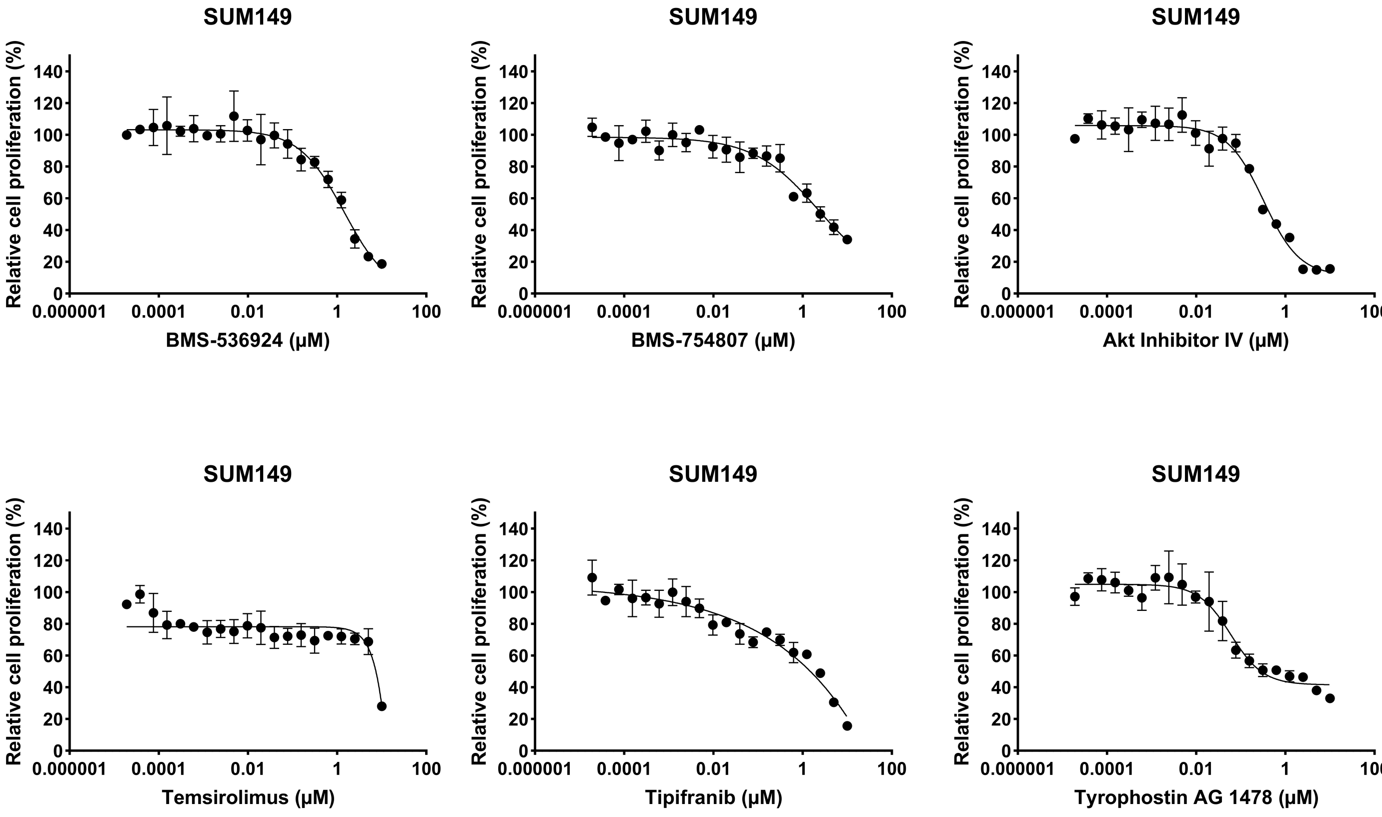

**Figure S4C-E**

C.

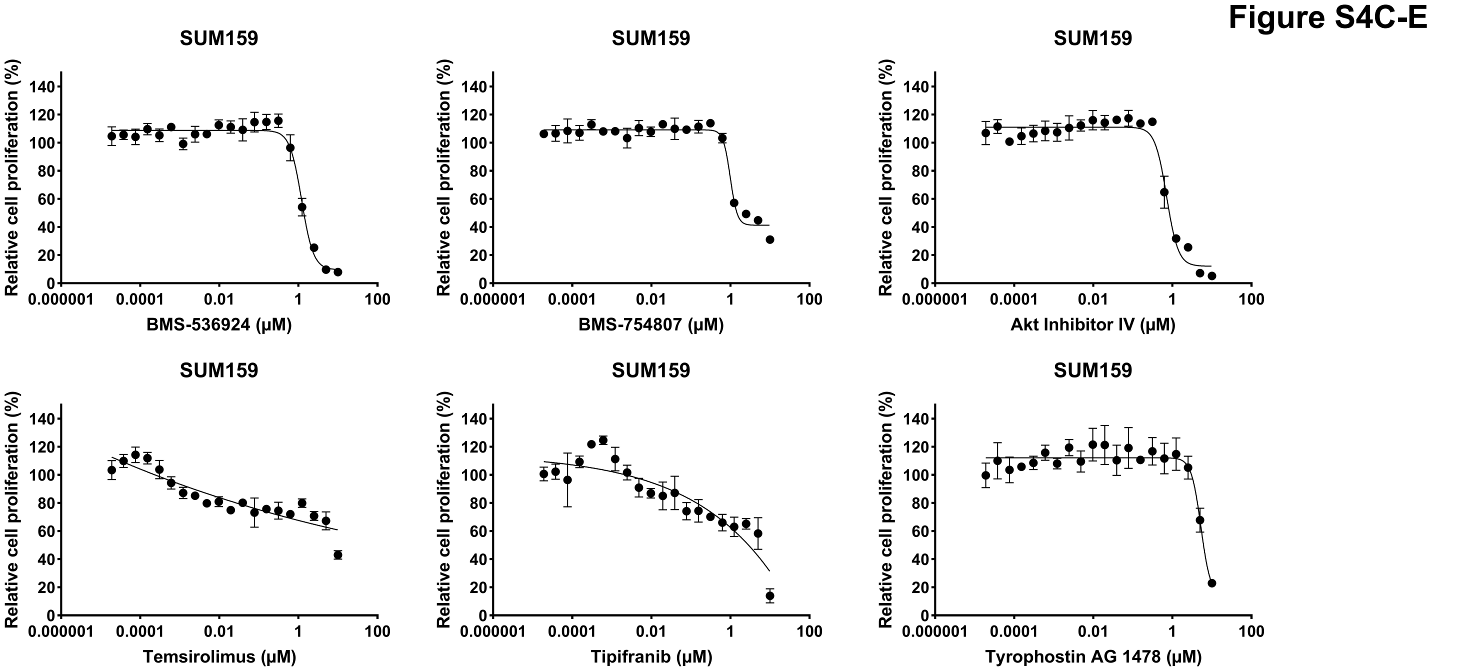


D.

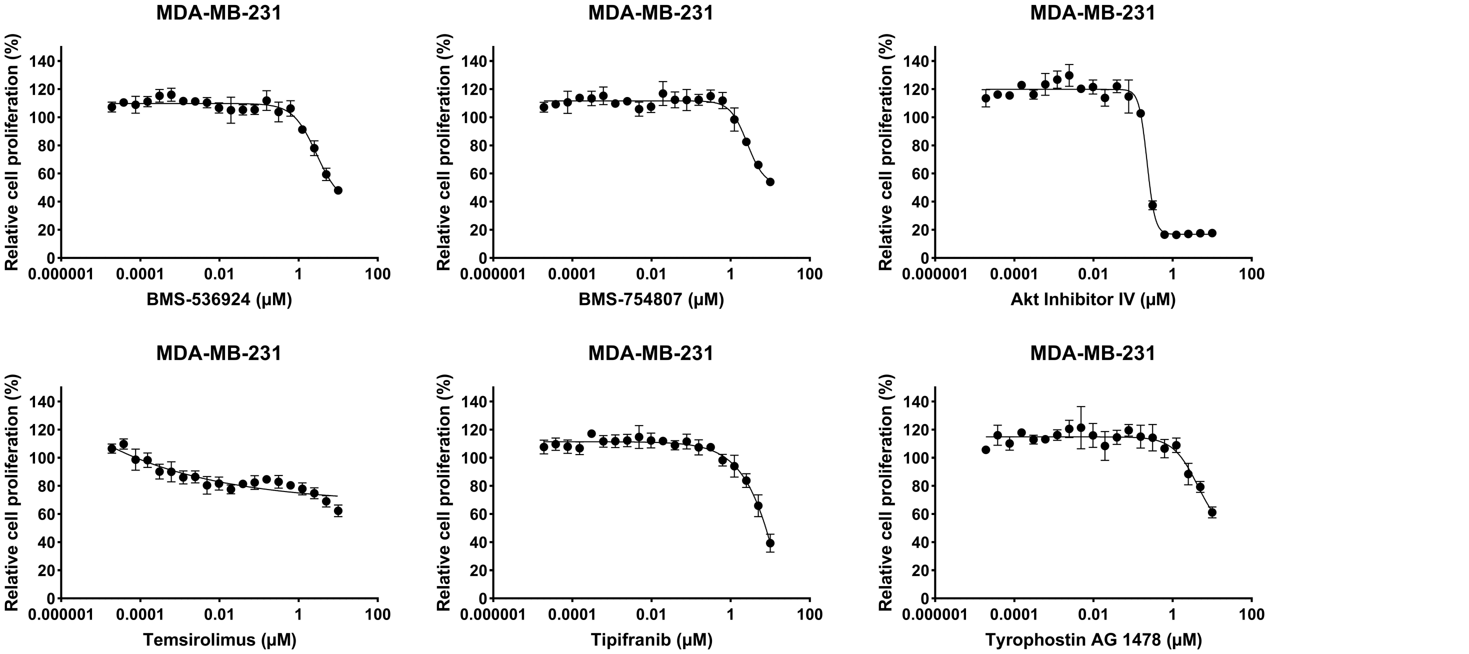


E.

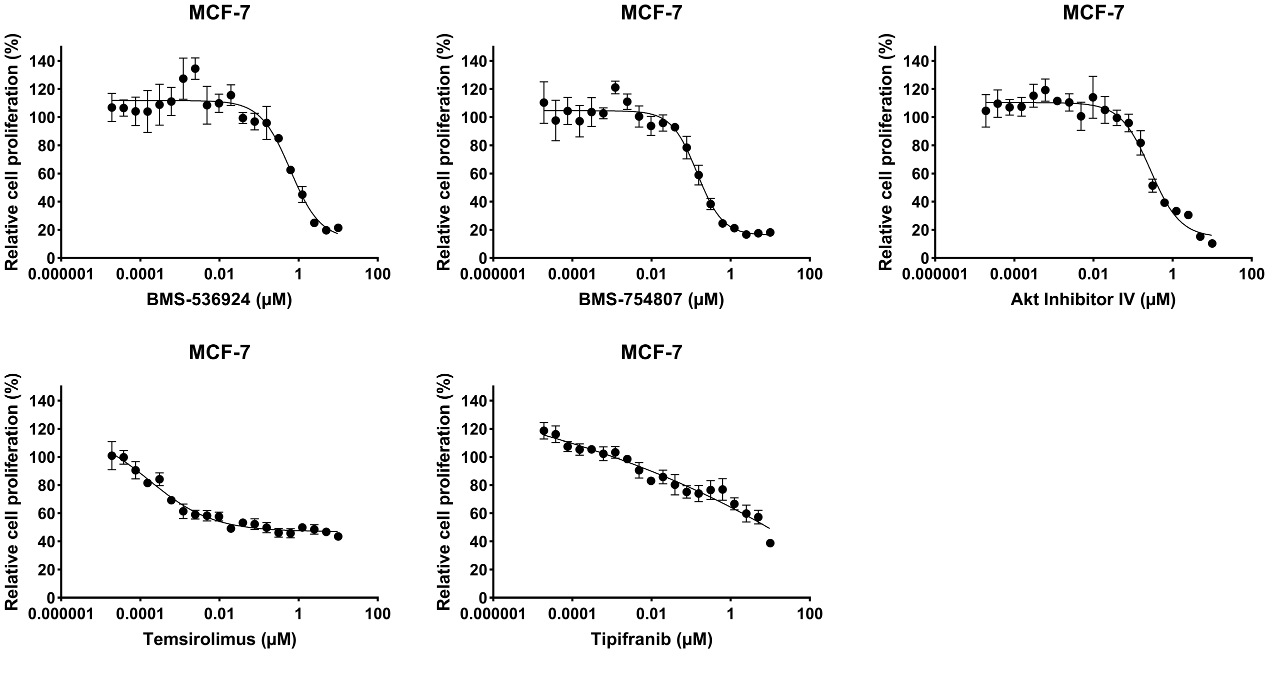

**Figure S5A-E**

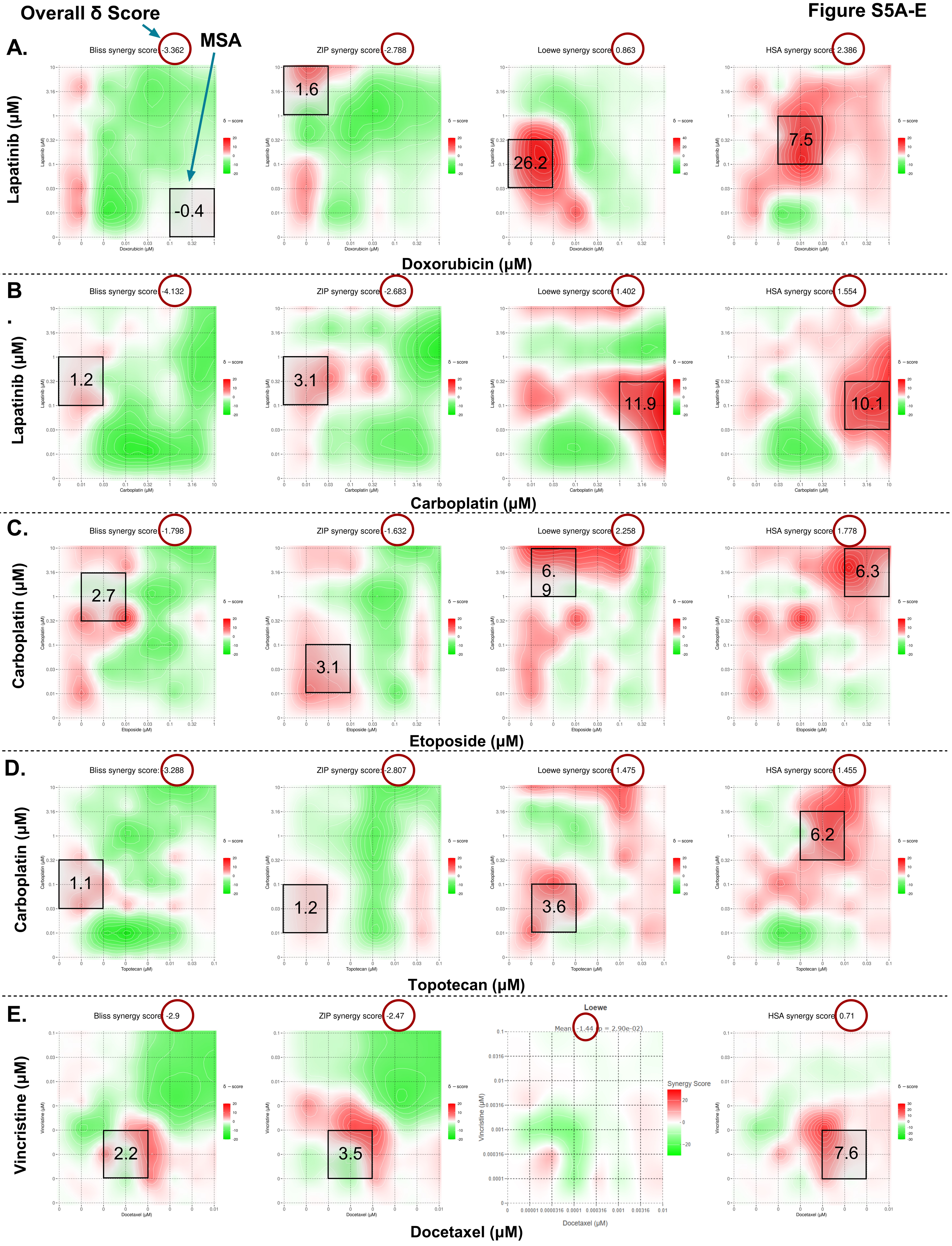
Overall δ Score
MSA
A.
Bliss synergy score: -3.362
ZIP synergy score: -2.788
Loewe synergy score: 0.863
HSA synergy score: 2.386
-0.4
1.6
26.2
7.5
Lapatinib (μM)
Doxorubicin (μM)
B
Bliss synergy score: -4.132
ZIP synergy score: -2.683
Loewe synergy score: 1.402
HSA synergy score: 1.554
1.2
3.1
11.9
10.1
Lapatinib (μM)
Carboplatin (μM)
C.
Bliss synergy score: -1.798
ZIP synergy score: -1.632
Loewe synergy score: 2.258
HSA synergy score: 1.778
2.7
3.1
6.9
6.3
Carboplatin (μM)
Etoposide (μM)
D.
Bliss synergy score: -3.288
ZIP synergy score: -2.807
Loewe synergy score: 1.475
HSA synergy score: 1.455
1.1
1.2
3.6
6.2
Carboplatin (μM)
Topotecan (μM)
E.
Bliss synergy score: -2.9
ZIP synergy score: -2.47
Loewe
Mean: -1.44 (p = 2.90e-02)
Synergy Score
HSA synergy score: 0.71
2.2
3.5
7.6
Vincristine (μM)
Docetaxel (μM)
δ – score

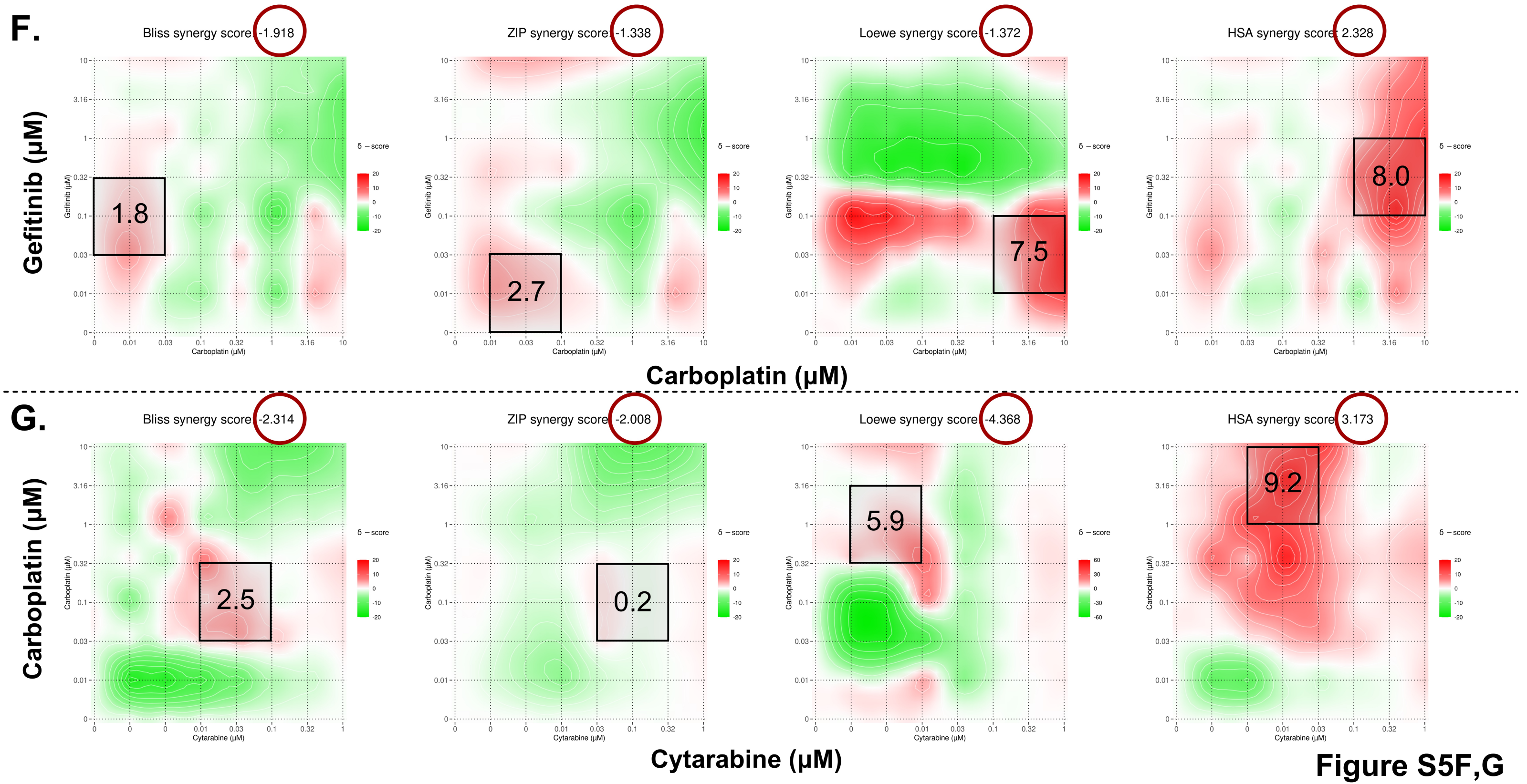

F.
Bliss synergy score: -1.918
ZIP synergy score: -1.338
Loewe synergy score: -1.372
HSA synergy score: 2.328
Gefitinib (μM)
1.8
2.7
7.5
8.0
Carboplatin (μM)
G.
Bliss synergy score: -2.314
ZIP synergy score: -2.008
Loewe synergy score: -4.368
HSA synergy score: 3.173
Carboplatin (μM)
2.5
0.2
5.9
9.2
Cytarabine (μM)


Figure S5F,G

**Figure S6A-C**

**A.**

## 2D

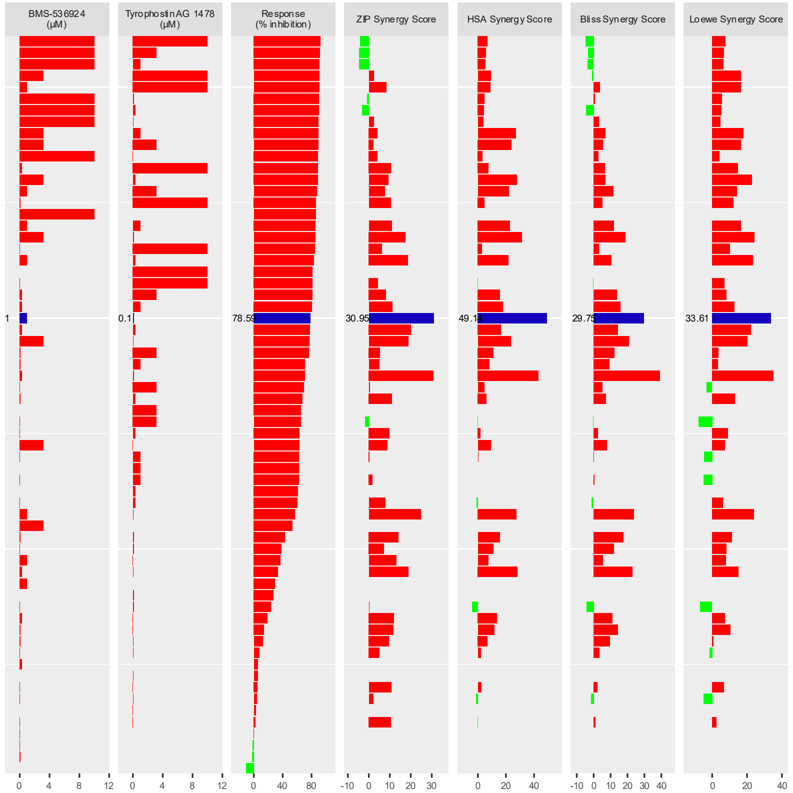


## 3D

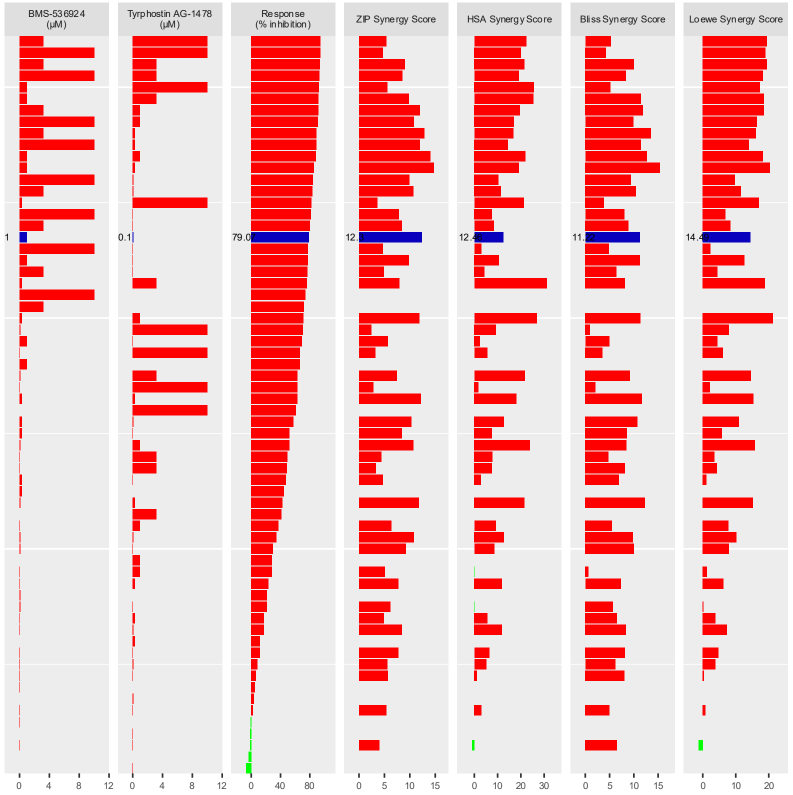

**B.**

## 2D

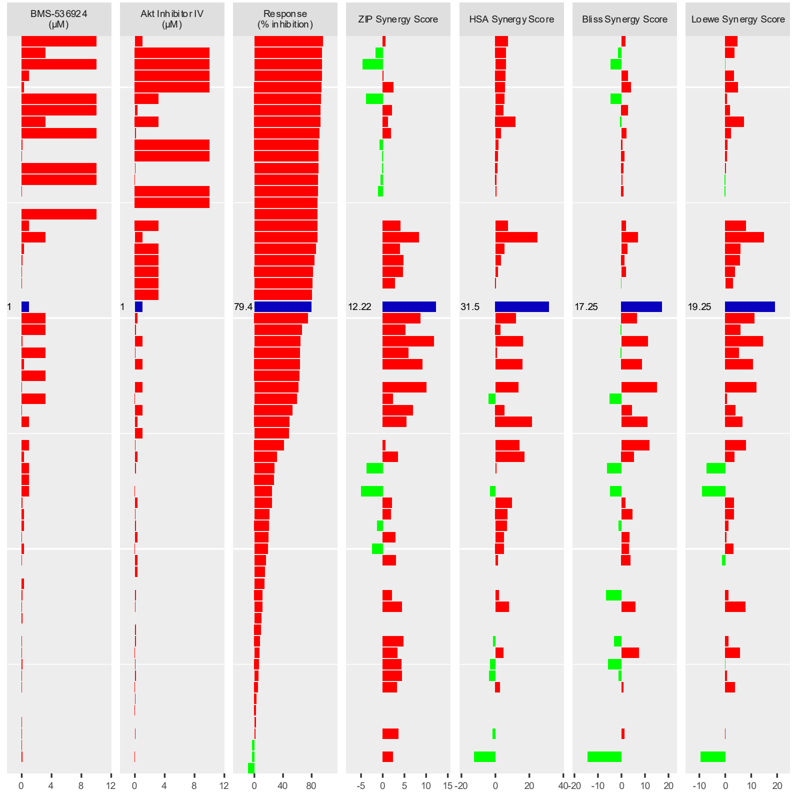

BMS-536924 (μM)
Akt Inhibitor IV (μM)
Response (% inhibition)
ZIP Synergy Score
HSA Synergy Score
Bliss Synergy Score
Loewe Synergy Score
1
1
79.4
12.22
31.5
17.25
19.25


## 3D

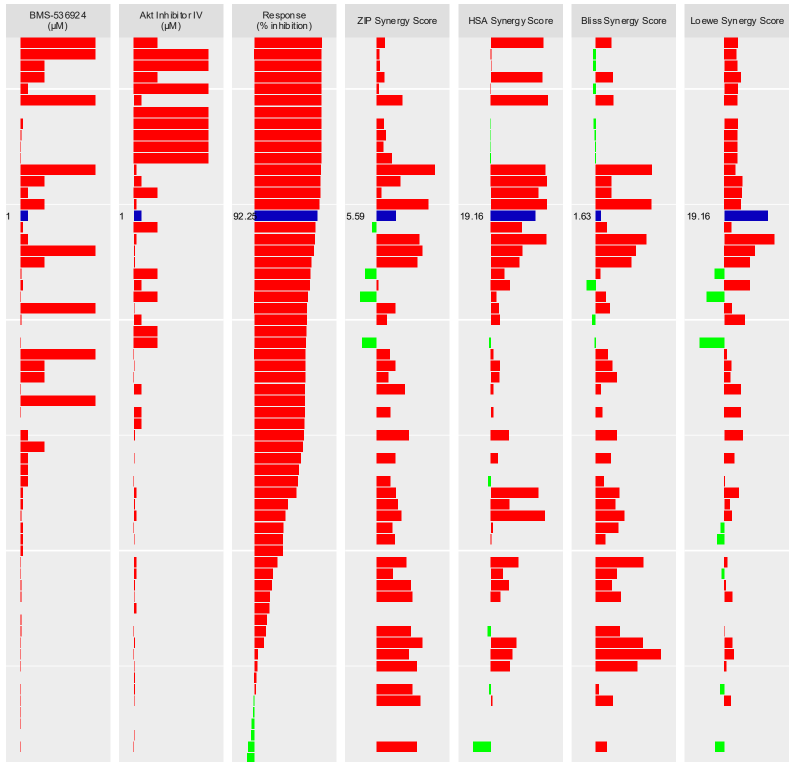

BMS-536924 (μM)
Akt Inhibitor IV (μM)
Response (% inhibition)
ZIP Synergy Score
HSA Synergy Score
Bliss Synergy Score
Loewe Synergy Score
1
1
92.25
5.59
19.16
1.63
19.16

**C.** **2D**

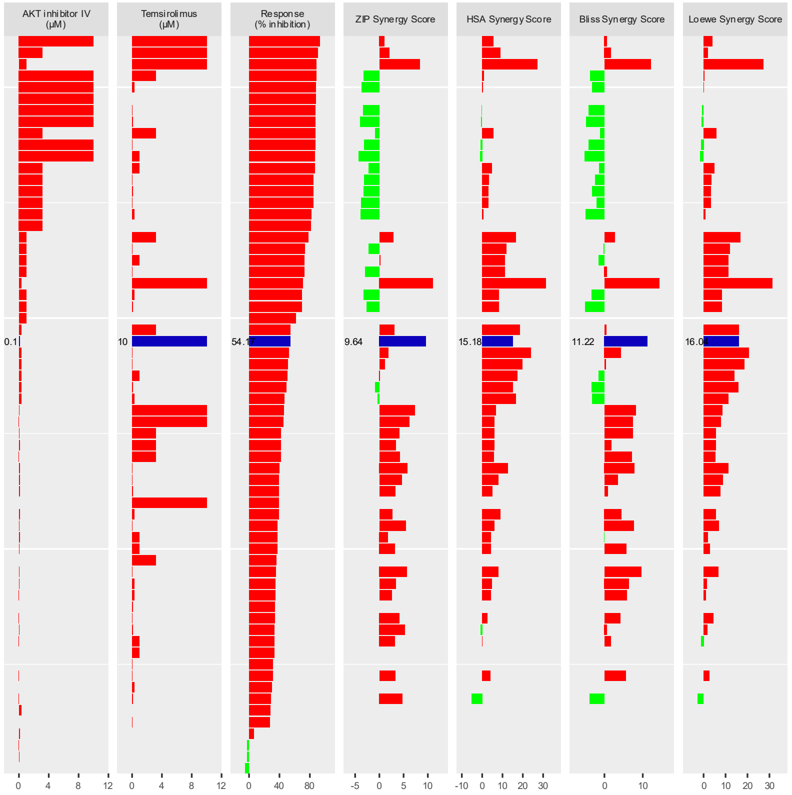
AKT inhibitor IV (µM)
Temsirolimus (µM)
Response (% inhibition)
ZIP Synergy Score
HSA Synergy Score
Bliss Synergy Score
Loewe Synergy Score
0.1
10
54.1
9.64
15.18
11.22
16.08
0 4 8 12
0 4 8 12
0 40 80
-5 0 5 10
-10 0 10 20 30
0 10
0 10 20 30

**3D**

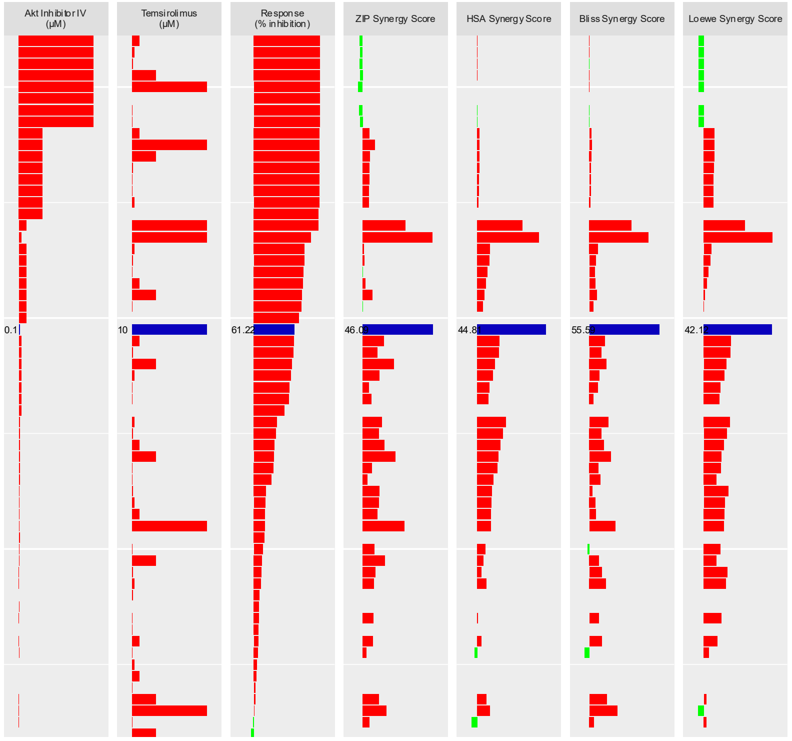
Akt Inhibitor IV (µM)
Temsirolimus (µM)
Response (% inhibition)
ZIP Synergy Score
HSA Synergy Score
Bliss Synergy Score
Loewe Synergy Score
0.1
10
61.22
46.0
44.8
55.5
42.1

### Figure S7

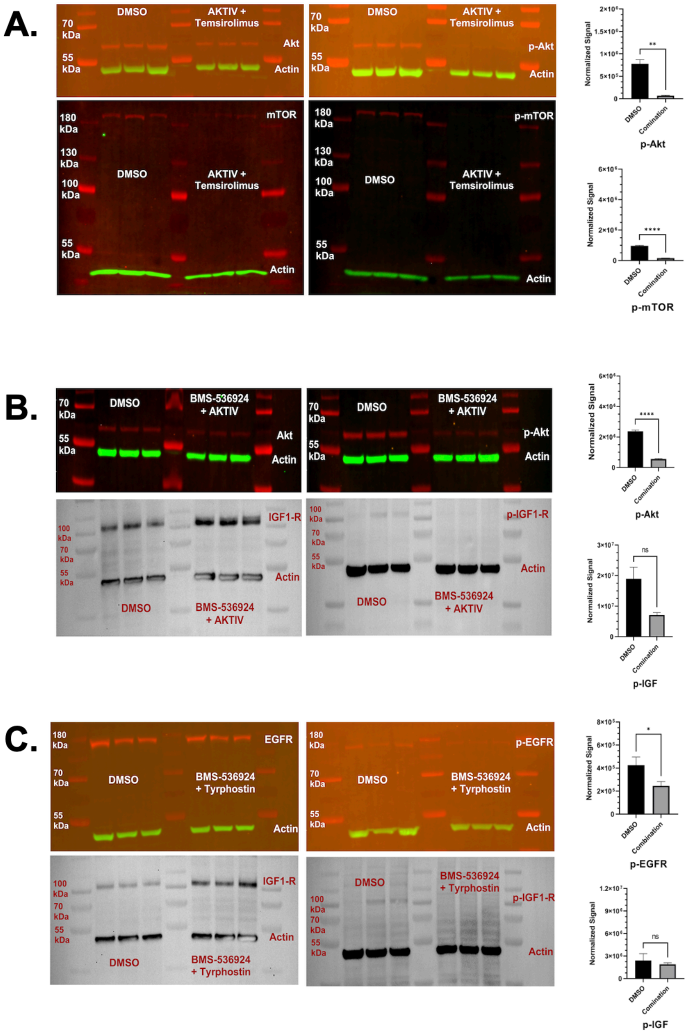
A.
DMSO
AKTIV + Temsirolimus
70 kDa
55 kDa
Akt
Actin
p-Akt
180 kDa
130 kDa
100 kDa
mTOR
p-mTOR
Normalized Signal
**
****
DMSO
Comination
p-Akt
p-mTOR
B.
BMS-536924 + AKTIV
IGF1-R
p-IGF1-R
ns
p-IGF
C.
EGFR
p-EGFR
BMS-536924 + Tyrphostin
*
Combination
p-EGFR
p-IGF

### Figure S8

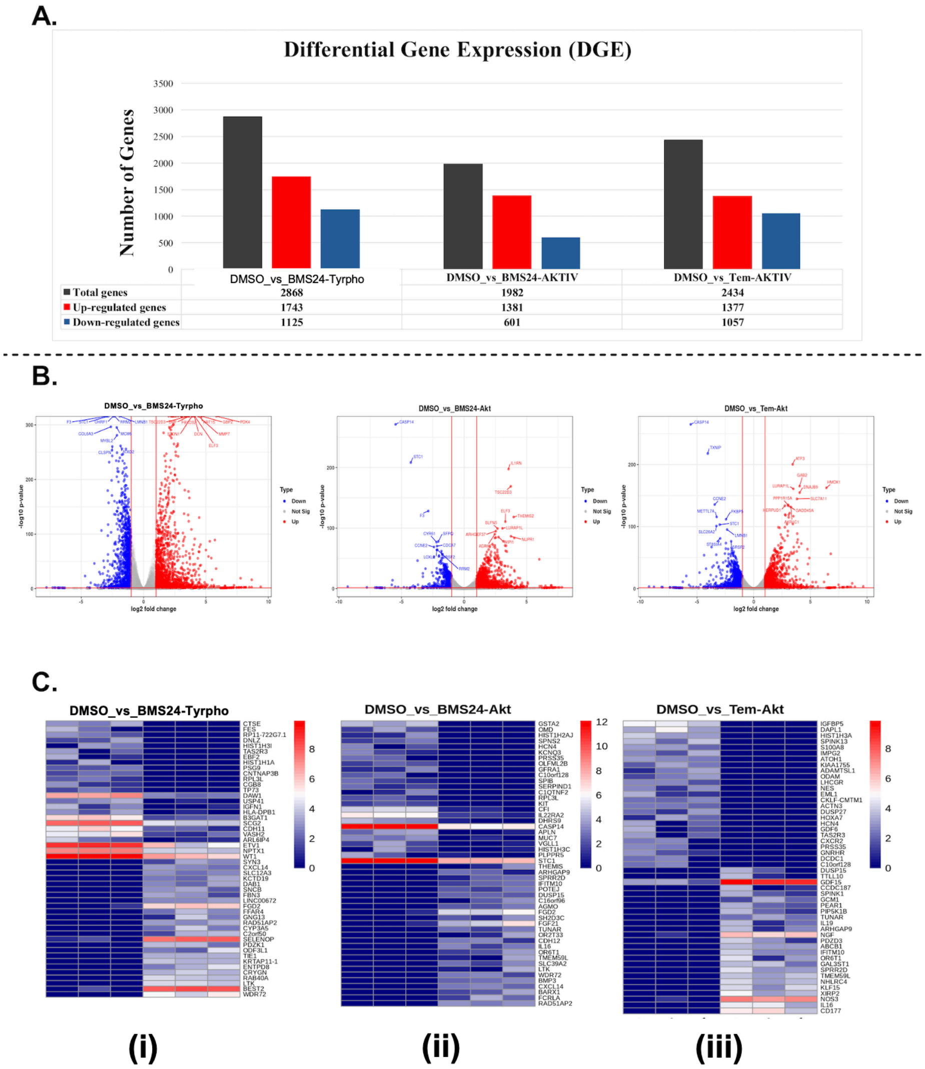
A.
Differential Gene Expression (DGE)
Number of Genes
3500
3000
2500
2000
1500
1000
500
0
DMSO_vs_BMS24-Tyrpho
DMSO_vs_BMS24-AKTIV
DMSO_vs_Tem-AKTIV
Total genes 2868 1982 2434
Up-regulated genes 1743 1381 1377
Down-regulated genes 1125 601 1057
B.
DMSO_vs_BMS24-Tyrpho
DMSO_vs_BMS24-Akt
DMSO_vs_Tem-Akt
-log10 p-value
log2 fold change
Type
Down
Not Sig
Up
C.
DMSO_vs_BMS24-Tyrpho
DMSO_vs_BMS24-Akt
DMSO_vs_Tem-Akt
(i)
(ii)
(iii)

### Supporting Information

**S1 Table. The 24 LWAS-predicted drugs for IBC (from Ji *et al.*, 2020).** The table presents the corresponding drug class, PubMed entry, Clinical Trials entry, and NCT number.

| | Drug Name | Drug Bank ID [1] | Drug class (MOA) [2] | PubMed for IBC [3] | Clinical Trials for IBC [4] | NCT Number [5] |
|---|---|---|---|---|---|---|
| **Vinca alkaloids** | Vincristine | DB00541 | Microtubule destabilizer | Yes | No | -- |
| | Vinorelbine | DB00361 | Microtubule destabilizer | Yes | Yes | NCT01325428 |
| **Antimetabolite** | Methotrexate | DB00563 | DNA/RNA damage/inhibits dihydrofolate reductase (DHFR) | Yes | Yes | NCT00003680 |
| | 5-Fluorouracil | DB00544 | DNA/RNA damage/inhibits thymidylate synthase (TS) | Yes | Yes | NCT01036087<br>NCT00976989<br>NCT02132949 |
| | Gemcitabine | DB00441 | Inhibits DNA synthesis | Yes | Yes | NCT00193050<br>NCT00193206 |
| | Hydroxyurea | DB01005 | Inhibits DNA synthesis | No | No | -- |
| | Cytarabine | DB00987 | DNA replication inhibitor | No | No | -- |
| **Alkylating Agents** | Carmustine | DB00262 | Alkylates DNA | Yes | No | -- |
| | Ifosfamide | DB01181 | Alkylates DNA | Yes | No | -- |
| | Dacarbazine | DB00851 | Alkylates DNA | No | No | -- |
| | Carboplatin | DB00958 | Platinum-based alkylating agent (DNA crosslinker) | Yes | Yes | NCT01036087<br>NCT00118053<br>NCT05093387<br>NCT00251329 |
| **Anthracycline** | Daunorubicin | DB00694 | Intercalates DNA/ topoisomerase (topo) 2 inhibitor/DNA breakage | Yes | No | -- |
| | Doxorubicin | DB00997 | Intercalates DNA/ topo 2 inhibitor/DNA breakage | Yes | Yes | NCT00004925<br>NCT00005822<br>NCT00016406<br>NCT00005800 |
| **Topoisomerase inhibitor** | Etoposide | DB00773 | Plant alkaloids (topo 2 inhibitor) | Yes | Yes | NCT00001507 |
| | Mitoxantrone | DB01204 | (Intercalates DNA/DNA damage/topo inhibitor) | Yes | No | -- |
| | Topotecan | DB01030 | Topoisomerase 1 inhibitor | No | No | -- |
| **Taxane** | Paclitaxel | DB01229 | Tubulin stabilizer/mitotic inhibitor | Yes | Yes | NCT00111787<br>NCT01036087<br>NCT00001507<br>NCT02132949 |
| | Docetaxel | DB01248 | Tubulin stabilizer/mitotic inhibitor | Yes | Yes | NCT00193050<br>NCT00066443<br>NCT00118053<br>NCT00017095 |

| | | | | | | |
|---|---|---|---|---|---|---|
| **Kinase inhibitor** | Gefitinib | DB00317 | EGFR Tyrosine Kinase (TK) inhibitor | Yes | No | -- |
| | Imatinib | DB00619 | TK inhibitor (BCR-ABL, KIT, PDGF-R) | Yes | No | -- |
| | Sunitinib | DB01268 | TK inhibitor (VEGFR/PDGFR/c-kit) | No | Yes | NCT00513695 |
| | Lapatinib | DB01259 | TK inhibitor (EGFR/HER2) | Yes | Yes | NCT00105950<br>NCT00558103<br>NCT00111787<br>NCT00450892 |
| **Peptide drug** | Octreotide | DB00104 | For metastatic carcinoid tumors and vasoactive intestinal peptide secreting tumors | Yes | No | -- |
| **Glucocort-icoids** | Prednisone | DB00635 | Inhibit NF-Kappa B and other inflammatory transcription factors | Yes | No | -- |

[1] Drug Bank ID; [2] Mechanism of action (MOA); [3] Status in PubMed for IBC; [4] Clinical Trials for IBC; [5] National Clinical Trial Identifier Number.

**S2 Table. The 19 GRR-predicted drugs/compounds (from Ji et al., 2023)**. The table shows with the corresponding drug class for each drug and its mechanism of action.

| Drug Name | Drug ID (PubChem) [1] | Drug Class [2] |
|---|---|---|
| **AG-14361** | 9840076 | DNA damage, anti-cancer → PARP-1 inhibitor |
| **AKTIV** | 5719375 | Cytotoxic and antiproliferative → Akt protein kinase Inhibitor |
| **Ambrisentan** | 6918493 | Vasodilator Endothelin → Receptor, GPCR & G Protein |
| **AZD-7545** | 16741245 | Adenocarcinoma → Selective inhibitor of PDHK2 |
| **BMS-536924** | 135440466 | Anti-cancer → IGF1-R kinase and IR inhibitor |
| **BMS-754807** | 24785538 | Anti-cancer → reversible IGF1-R/IR inhibitor |
| **Butalbital** | 2481 | CNS depressant → JAK/STAT signaling, P53 signaling, and NOTCH signaling pathway |
| **Clobenpropit** | 2790 | Anti-tumor → Histamine H3 receptor, PI3K/AKT pathway |
| **COT-10b** | | Acute myeloid leukemia → Serine/threonine MAP3 kinase |
| **Dapsone** | 2955 | Antibacterial → Sulfone drug<br>Anti-inflammatory → (Not fully understood) |
| **Letrozole** | 3902 | Aromatase Inhibitor<br>Antineoplastic Agent |
| **Medetomidine** | 68602 | Neurological Disease/Psychotic Disorders → selective α2-adrenoceptor agonist |
| **Nicardipine** | 4474 | Cardiovascular Disease → Calcium channel blocker |
| **Nimesulide** | 4495 | Selective COX-2 inhibitor (NSAID) |
| **Temsirolimus** | 6918289 | Antineoplastic (mTOR inhibitor) Immunomodulating agent |
| **Ticarcillin** | 36921 | Infection → Beta lactam antibiotic |
| **Tipifarnib-P2** | 159324 | Antineoplastic → Farnesyltransferase inhibitors |
| **Tyrphostin-AG-1478** | 2051 | Histiocytic Lymphoma → EGFR Tyrosine Kinase Inhibitor |
| **ZM336372** | 5730 | Histiocytic Lymphoma → Inhibitor of the MAP protein kinase c-Raf, JAK/STAT signaling. |

[1]PubChemID, [2]Drug Class.

**S3 Table. Efficacy of LWAS and GRR predicted drugs and compounds in four breast cancer cell lines using the MTT assay.** The table lists the $IC_{50}$ values determined for each drug from the MTT assay across the cell lines, with values reported as the mean ± SD from 2 independent experiments.

| Study | Drug | SUM149 ($IC_{50}$ μM) | SUM159 ($IC_{50}$ μM) | MDA-MB-231 ($IC_{50}$ μM) | MCF-7 ($IC_{50}$ μM) |
|---|---|---|---|---|---|
| LWAS [1] | Docetaxel | 0.0003 ± 2.3E-05 | 0.0009 | 0.002 ± 0.0009 | 0.002 |
| | Paclitaxel | 0.002 ± 0.0001 | 0.004 ± 0.0008 | 0.007 ± 0.002 | 0.003 |
| | Vincristine | 0.003 ± 0.001 | 0.005 ± 0.0009 | 0.004 ± 0.003 | 0.009 |
| | Vinorelbine | 0.2 ± 0.4 | 0.2 | 0.2 ± 0.03 | 0.1 |
| | Lapatinib | 0.7 ± 0.5 | 3.6 ± 1.2 | 7.3 | 5.6 |
| | Sunitinib | 2.1 ± 0.05 | 1.8 ± 0.3 | 2.1 ± 0.6 | 2.04 |
| | Gefitinib | 0.1 ± 0.02 | 11 ± 1.9 | -- | -- |
| | Doxorubicin | 0.3 ± 0.1 | 0.07 | 0.3 ± 0.1 | 0.2 |
| | Daunorubicin | 0.09 ± 0.01 | 0.05 ± 0.003 | 0.2 ± 0.08 | 0.1 |
| | Mitoxantrone | 0.08 ± 0.04 | 0.07 | 0.1 ± 0.05 | 0.05 |
| | Etoposide | 0.2 ± 0.1 | 0.35 | 7.4 | 0.7 |
| | Topotecan | 0.2 ± 0.2 | 0.02 | 0.8 ± 0.08 | 0.5 |
| | Gemcitabine | 0.002 ± 0.0003 | 0.004 ± 0.0004 | 0.3 ± 0.03 | 0.01 |
| | Cytarabine | 0.04 ± 0.004 | 0.05 | 0.8 ± 0.2 | 0.4 |
| | Carboplatin | 4.5 ± 2.2 | -- | -- | -- |
| | Methotrexate | 0.07 ± 0.09 | 0.1 | 0.5 ± 0.1 | 0.05 |
| | 5-Fluorouracil | 1.1 | 15.7 ± 3.8 | 1.9 | 2.5 |
| GRR [2] | BMS-536924 | 1.5 ± 0.1 | 1.2 ± 0.2 | 6.7 | 0.8 ± 0.1 |
| | BMS-754807 | 3.6 ± 0.9 | 2.3 ± 0.7 | 8.0 | 0.2 ± 0.1 |
| | AKTIV | 0.6 ± 0.2 | 0.7 ± 0.2 | 0.3 | 0.4 ± 0.1 |
| | Temsirolimus | 7.3 ± 1.7 | 0.0005 ± 0.0005 | 0.0003 | 0.0005 ± 0.0004 |
| | Tipifarnib | 1.3 ± 0.1 | 2.1 ± 0.7 | 6.4 | 3.8 ± 4.8 |
| | Tyrphostin AG-1478 | 0.4 ± 0.2 | 5.7 ± 0.3 | 10.9 | -- |

[1] Drugs identified by LWAS in Ji et al., 2020; [2] Drugs identified by GRR in Ji et al., 2023.

**S4 Table. Drugs selected for combination studies.** Selected compounds from LWAS and GRR active lists for combination studies in SUM149, along with their corresponding $EC_{25}$ values and the highest concentration used for the 8 x 8 matrix.

| | Compounds | $IC_{50}$ Average "Hoechst Assay" (μM) | $EC_{25}$ (μM) | Highest Conc. in 8 × 8 Matrix (μM) |
|---|---|---|---|---|
| **LWAS** | Doxorubicin | 0.01 | 0.004 | 1 |
| | Mitoxantrone | 0.005 | -- | -- |
| | Etoposide | 0.07 | 0.023 | 1 |
| | Topotecan | 0.008 | 0.003 | 0.1 |
| | Docetaxel | 0.0004 | 0.0002 | 0.01 |
| | Vincristine | 0.004 | 0.002 | 0.1 |
| | Carboplatin | 4.4 | 1.5 | 10 |
| | Fluorouracil | 4.8 | -- | -- |
| | Lapatinib | 0.63 | 0.12 | 10 |
| | Gefitinib | 0.4 | 0.2 | 10 |
| | Sunitinib | 1.4 | -- | -- |
| | Gemcitabine | 0.0012 | -- | -- |
| | Cytarabine | 0.025 | 0.008 | 1 |
| **GRR** | BMS 536924 | 1.9 | 0.7 | 10 |
| | BMS 754807 | 0.5 | 0.2 | 10 |
| | AKTIV | 0.22 | 0.1 | 10 |
| | Temsirolimus | 4.3 | 1.5 | 10 |
| | Tipifarnib | 0.075 | 0.03 | 10 |
| | Tyrphostin AG-1478 | 0.23 | 0.1 | 10 |
